# Evolutionary superscaffolding and chromosome anchoring to improve *Anopheles* genome assemblies

**DOI:** 10.1101/434670

**Authors:** Robert M. Waterhouse, Sergey Aganezov, Yoann Anselmetti, Jiyoung Lee, Livio Ruzzante, Maarten J.M.F. Reijnders, Romain Feron, Sèverine Bérard, Phillip George, Matthew W. Hahn, Paul I. Howell, Maryam Kamali, Sergey Koren, Daniel Lawson, Gareth Maslen, Ashley Peery, Adam M. Phillippy, Maria V. Sharakhova, Eric Tannier, Maria F. Unger, Simo V. Zhang, Max A. Alekseyev, Nora J. Besansky, Cedric Chauve, Scott J. Emrich, Igor V. Sharakhov

**Author notes:** Correspondence should be addressed to (RMW), (IVS).

## Abstract

**Background:** New sequencing technologies have lowered financial barriers to whole genome sequencing, but resulting assemblies are often fragmented and far from ‘finished’. Updating multi-scaffold drafts to chromosome-level status can be achieved through experimental mapping or re-sequencing efforts. Avoiding the costs associated with such approaches, comparative genomic analysis of gene order conservation (synteny) to predict scaffold neighbours (adjacencies) offers a potentially useful complementary method for improving draft assemblies.

**Results:** We employed three gene synteny-based methods applied to 21 *Anopheles* mosquito assemblies to produce consensus sets of scaffold adjacencies. For subsets of the assemblies we integrated these with additional supporting data to confirm and complement the synteny-based adjacencies: six with physical mapping data that anchor scaffolds to chromosome locations, 13 with paired-end RNA sequencing (RNAseq) data, and three with new assemblies based on re-scaffolding or Pacific Biosciences long-read data. Our combined analyses produced 20 new superscaffolded assemblies with improved contiguities: seven for which assignments of non-anchored scaffolds to chromosome arms span more than 75% of the assemblies, and a further seven with chromosome anchoring including an 88% anchored *Anopheles arabiensis* assembly and, respectively, 73% and 84% anchored assemblies with comprehensively updated cytogenetic photomaps for *Anopheles funestus* and *Anopheles stephensi*.

**Conclusions:** Experimental data from probe mapping, RNAseq, or long-read technologies, where available, all contribute to successful upgrading of draft assemblies. Our comparisons show that gene synteny-based computational methods represent a valuable alternative or complementary approach. Our improved *Anopheles* reference assemblies highlight the utility of applying comparative genomics approaches to improve community genomic resources.

## Introduction

Reduced costs of new sequencing technologies have facilitated the rapid growth of draft genome assemblies from all kingdoms of life. Nevertheless, the process of progressing from draft status to that of a ‘finished’ reference genome—a near-complete and near-contiguous chromosome-level assembly—remains the exclusive accomplishment of relatively few species. Chromosomal ordering and orienting of contigs or scaffolds may be achieved by experimental approaches including fluorescence *in situ* hybridization (FISH) (Bauman et al. 1980), genetic linkage mapping (Hahn et al. 2014; Fierst 2015), optical (restriction site) mapping (Levy-Sakin and Ebenstein 2013), or analysis of chromatin interaction frequency data (Kaplan and Dekker 2013; Burton et al. 2013). When resources allow, combined approaches can produce excellent results, e.g. for Brassicaceae plants (Jiao et al. 2017), the three-spined stickleback (Peichel et al. 2017), and the mosquitoes, *Aedes aegypti* and *Culex quinquefasciatus* (Dudchenko et al. 2017; Matthews et al. 2018).

While many research applications may not strictly require such high-quality assemblies, improvements in completeness, contiguity, and chromosome anchoring or assignments can substantially add to the power and breadth of biological and evolutionary inferences from comparative genomics or population genetics analyses. For example, extensive contiguity and chromosome-level anchoring are clearly important when addressing questions concerning karyotype evolution or smaller-scale inversions and translocations, re-sequencing analyses of population-level samples, reconstructing rearrangement-based phylogenies, identifying and characterising genes that localise within quantitative trait loci (QTL), examining genomic sexual conflicts, or tracing drivers of speciation. In many such studies, assembly improvements were critical to enable more robust analyses, e.g. QTL analysis with rape mustard flowering-time phenotypes (Markelz et al. 2017); contrasting genomic patterns of diversity between barley cultivars (Mascher et al. 2017); defining rearrangements of the typical avian karyotype (Damas et al. 2017); detecting chromosome fusion events during butterfly evolution (Davey et al. 2016); characterising the ancestral lepidopteran karyotype (Ahola et al. 2014); identifying the chromosomal position and structure of the male determining locus in *Ae. aegypti* (Matthews et al. 2018); and characterising a melon fly genetic sexing strain as well as localising the sexing trait (Sim and Geib 2017).

Available genome assemblies for anopheline mosquitoes vary considerably in contiguity and levels of chromosome anchoring. Sequencing the first mosquito genome produced an assembly for the *Anopheles gambiae* PEST strain with 8’987 scaffolds spanning 278 megabasepairs (Mbp), where 303 scaffolds spanned 91% of the assembly and physical mapping assigned 84% of the genome to chromosome arms (Holt et al. 2002). Additional FISH mapping and orienting of 28 scaffolds and bioinformatics analyses later facilitated an assembly update by removing haplotype scaffolds and bacterial sequences and anchoring a third of previously unmapped scaffolds to chromosomes (Sharakhova et al. 2007). Since then, more than 20 new assemblies have been built for the anophelines, several with mapping efforts that enabled at least partial chromosome anchoring. Sequencing of the *A. gambiae* Pimperena S form and *Anopheles coluzzii* (formerly *A. gambiae* M form) produced assemblies with 13’050 and 10’525 scaffolds, respectively, with 89% of each of these assemblies alignable to the closely related PEST assembly (Lawniczak et al. 2010). The much smaller 174 Mbp assembly of the more distantly related neotropical vector, *Anopheles darlingi*, comprised 8’233 scaffolds, but they remained unanchored (Marinotti et al. 2013). Physical mapping assigned 62% of the 221 Mbp *Anopheles stephensi* Indian strain assembly (23’371 scaffolds) (Jiang et al. 2014) and 36% of the *Anopheles sinensis* Chinese strain assembly (9’597 scaffolds) (Zhou et al. 2014; Wei et al. 2017) to polytene chromosomes. The *Anopheles* 16 Genomes Project (Neafsey et al. 2013) produced assemblies ranging from a few hundred to several thousand scaffolds and used mapping data from four species to anchor *Anopheles funestus* (35%), *Anopheles atroparvus* (40%), *A. stephensi* SDA-500 strain (41%), and *Anopheles albimanus* (76%) genomes to chromosome arms (Neafsey et al. 2015). Additional physical mapping data for *A. atroparvus* subsequently improved this initial assembly to 90% chromosome anchoring (Artemov et al. 2018a), and for *A. albimanus* to 98% (Artemov et al. 2017).

For a genus such as *Anopheles* with already more than 20 genome assemblies available (Ruzzante et al. 2018), cross-species comparisons to identify potentially neighbouring scaffolds could facilitate assembly upgrades with improved contiguities. While genome rearrangements can and do occur, multiple homologous genomic regions with conserved orders and orientations, i.e. regions with maintained synteny, offer an evolutionarily guided approach for assembly improvement. Specifically, employing orthologous genes as conserved markers allows for the delineation of maintained syntenic blocks that provide support for putative scaffold adjacencies. Here we present results from applying three synteny-based computational approaches to perform evolutionarily guided assembly improvements of multiple *Anopheles* genomes. The consensus predictions offer well-supported sets of scaffold adjacencies that lead to improved assembly contiguities without the associated costs or time-investments required for experimental superscaffolding. Integrating these predictions with experimental data for subsets of the anophelines supported many adjacencies and highlighted the complementarity of experimental and computational approaches. Providing support for experimental results, complementary data to enhance improvements, or independent evidence for assembly validations, these evolutionarily guided methods offer a handy set of utensils in any genome assembly toolbox—here applied to improve available genomic resources of *Anopheles* mosquitoes.

## Results

### New reference genome assemblies and chromosome maps

New genome assemblies with scaffolds and superscaffolds anchored or assigned to chromosome arms were generated by leveraging evolutionary relationships to predict scaffold adjacencies and combining these with additional experimental data for subsets of the anophelines (**Fig. 1**). Integrating results from three gene synteny-based computational approaches to build superscaffolds from all scaffold neighbours and reconciling these with the experimental datasets resulted in 20 new assemblies with improved contiguities (**Table 1**), as well as chromosome mapping spanning 88% of the *Anopheles arabiensis* assembly, and updated chromosome maps for six other anophelines (**Table 2**). The synteny-based adjacencies were used to define well-supported consensus sets, which were then validated with and complemented by physical mapping and/or RNAseq and/or re-sequencing data for 14 assemblies. This followed a reconciliation workflow to integrate the different sets of scaffold adjacencies from synteny, physical mapping, RNAseq, or alignment data for each assembly (**see Methods; Supplementary Online Material: Figure S1**). Applying this integrative approach produced updated reference assemblies with increased scaffold N50 values (a median-like metric where half the genome is assembled into scaffolds of length N50 or longer) and reduced scaffold counts (**Table 1**). Although superscaffold contiguity levels remain variable, e.g. *A. atroparvus* 200 Mbp in 34 superscaffolds and *A. funestus* 182 Mbp in 113 superscaffolds, the total span of scaffolds that now form part of superscaffolds comprises more than half of ten of the assemblies, ranging from 113 Mbp for *Anopheles minimus* to 222 Mbp for *A. arabiensis* (**Supplementary Online Material: Figure S2**).

**Table 1.**
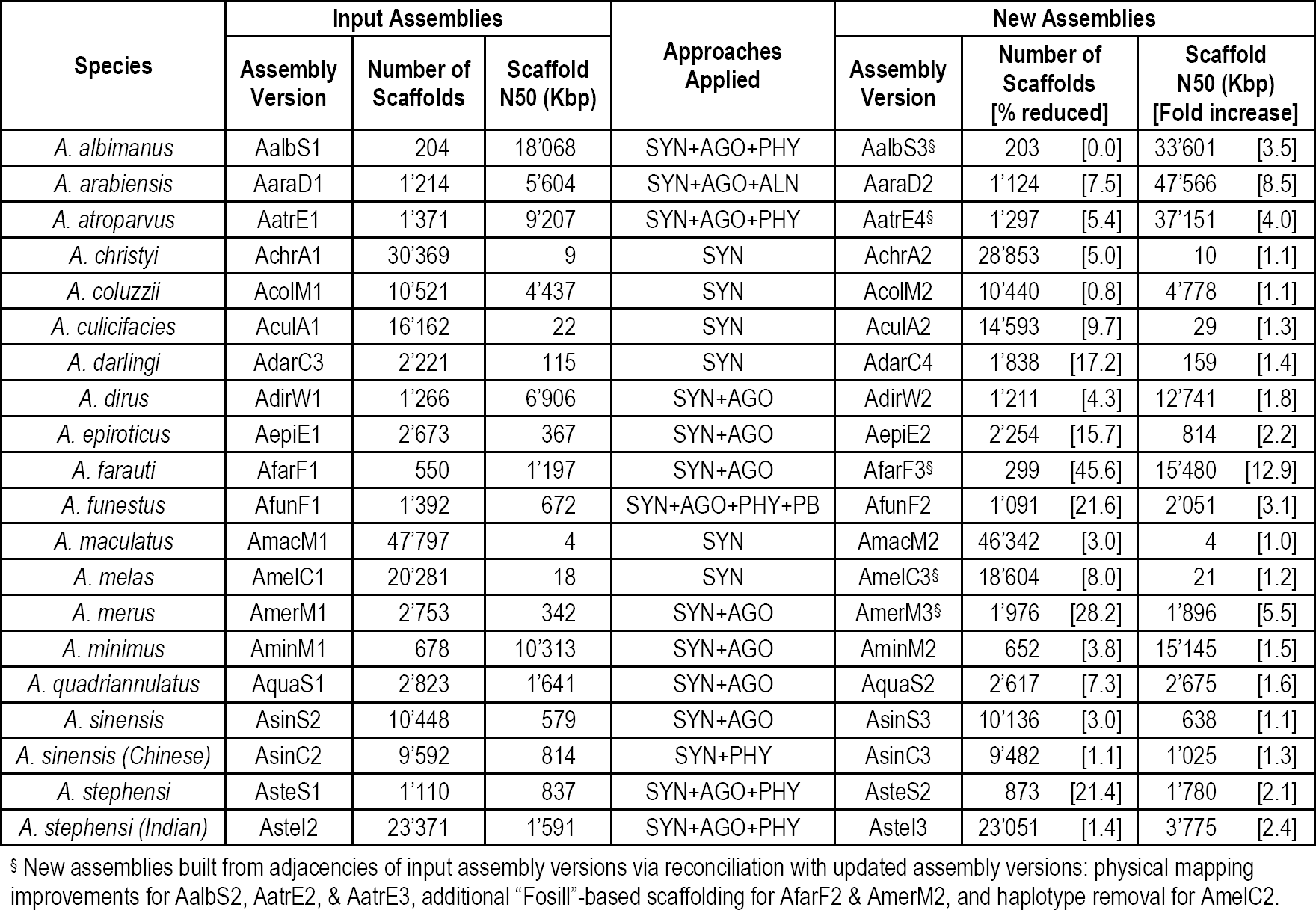
Summary statistics of the 20 input and new improved *Anopheles* assemblies. Summary statistics of scaffold counts and N50 values of the 20 input and improved *Anopheles* assemblies after applying synteny-based (SYN), and/or RNAseq AGOUTI-based (AGO), and/or alignment-based (ALN), and/or physical mapping-based (PHY), and/or PacBio sequencing-based (PB) approaches.

**Table 2.** Summary of anchoring improvements for seven anophelines with chromosome mapping data. Summary of scaffold counts and genomic spans added to the initial chromosome maps from synteny-based (SYN) and RNAseq AGOUTI-based (AGO) adjacencies, and counts of chromosome-mapped scaffolds that gained oriented neighbours after incorporating the SYN and AGO scaffold adjacencies.

| Assembly | Mapped scaffolds | Scaffolds added to map by: |  |  | Total scaffolds added | Mapped scaffolds now with oriented neighbours | Total basepairs added | % of assembly added | Total % of assembly anchored |
| --- | --- | --- | --- | --- | --- | --- | --- | --- | --- |
|  |  | Synteny | AGOUTI | SYN+AGO |  |  |  |  |  |
| *A. albimanus* | 31 | 0 | 2 | 0 | 2 | 0 | 2'160 | 0.00 | 98.26 |
| *A. arabiensis* | 51 | 4 | 2 | 0 | 6 | 0 | 256'948 | 0.10 | 87.84 |
| *A. atroparvus* | 46 | 5 | 7 | 3 | 9 | 0 | 870'748 | 0.39 | 89.75 |
| *A. funestus* | 202 | 89 | 45 | 34 | 100 | 81 | 26'434'544 | 11.73 | 72.91 |
| *A. sinensis (Chinese)* | 52 | 18 | NA | NA | 18 | 14 | 5'791'225 | 2.62 | 40.41 |
| *A. stephensi* | 99 | 102 | 52 | 45 | 110 | 77 | 47'779'259 | 21.20 | 61.96 |
| *A. stephensi (Indian)* | 118 | 76 | 47 | 33 | 90 | 92 | 10'975'818 | 4.96 | 83.66 |

**Figure 1.**
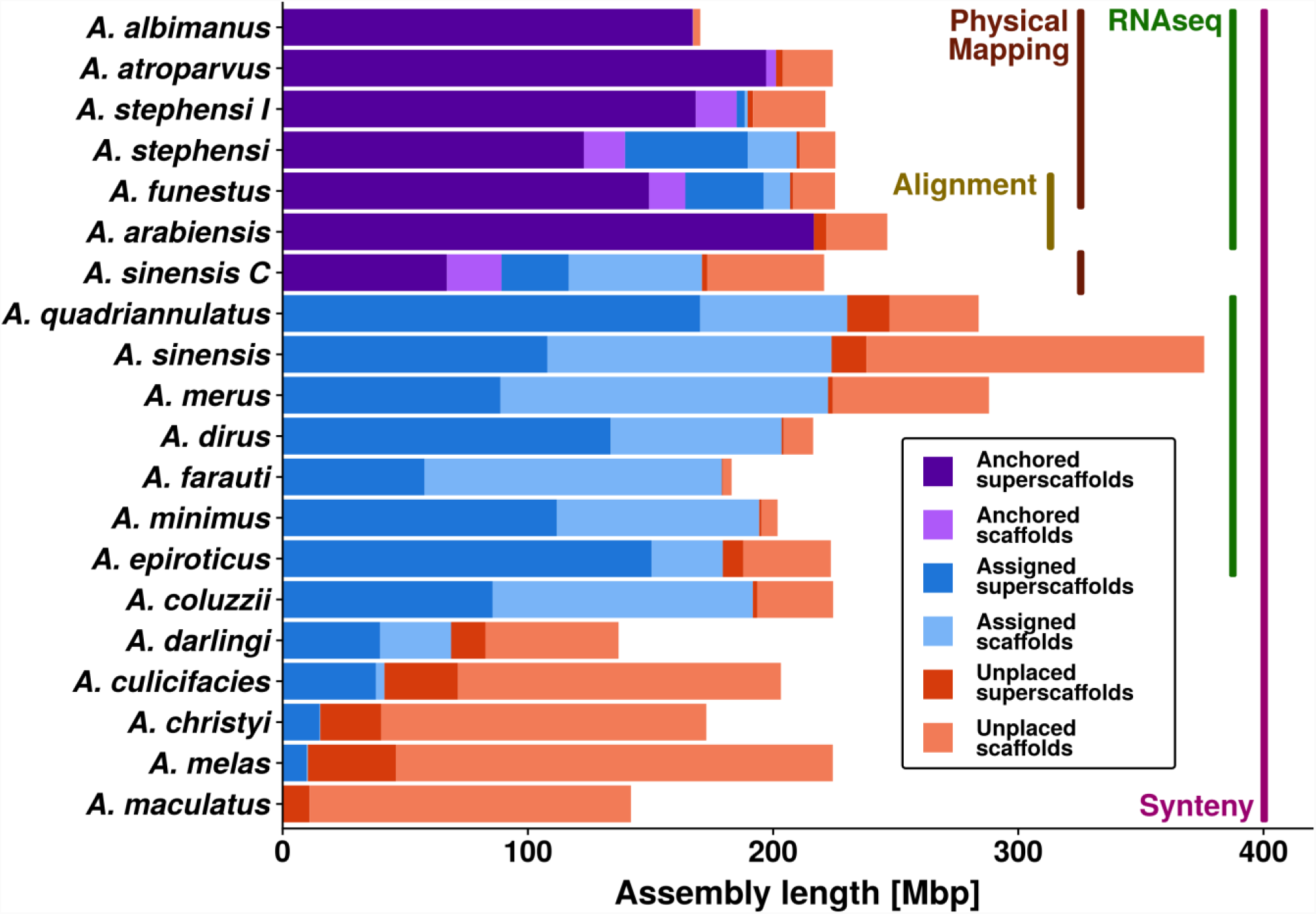
Genomic spans of scaffolds and superscaffolds with and without chromosome anchoring or arm assignments for 20 improved *Anopheles* assemblies. Consensus gene synteny-based methods were employed across the 21-assembly input dataset (also including *Anopheles gambiae*) to delineate scaffold adjacencies and build new superscaffolded assemblies with improved contiguities. These were integrated with results from additional complementary approaches for subsets of the anophelines including transcriptome (RNAseq) and genome sequencing data, whole-genome alignments, and chromosome anchoring data from physical mapping of probes. Chromosome mapping data for seven assemblies enabled anchoring of superscaffolds and scaffolds to their chromosomal locations (purple colours). Enumerating shared orthologues further enabled the assignment of non-anchored superscaffolds and scaffolds to chromosome arms (blue colours). Unplaced superscaffolds and scaffolds (orange colours) still comprise the majority of the least contiguous input assemblies, but they make up only a small proportion of the assemblies for which the available data allowed for substantial improvements to assembly contiguity and/or anchoring and/or arm assignments. Results for two strains are shown for *Anopheles sinensis*, SINENSIS and Chinese (C), and *Anopheles stephensi*, SDA-500 and Indian (I).

The greatest reductions in the total numbers of scaffolds were achieved for some of the least contiguous input assemblies including *Anopheles christyi, Anopheles culicifacies, Anopheles maculatus*, and *Anopheles melas* (**Table 1**). Given the heterogeneity of the input assemblies the relative changes highlight some of the most dramatic improvements, e.g. the *A. funestus* and *A. stephensi* (SDA-500) scaffold counts both dropped by almost 22% and the newly anchored *A. arabiensis* assembly resulted in an 8.5-fold larger N50 value (**Table 1**). For the anophelines with chromosome mapping data, the contributions of the synteny-based and/or RNAseq-based adjacencies to the numbers and genomic spans of anchored scaffolds were largest for *A. stephensi* (SDA-500) and *A. funestus*, but negligible or low for the recently updated *A. albimanus* (Artemov et al. 2017), *A. atroparvus* (Artemov et al. 2018a), and *A. sinensis* (Chinese) (Wei et al. 2017) assemblies (**Table 2**). The two *A. stephensi* assemblies achieved updated assembly anchoring of 62% and 84% (both improvements of more than 20%) and *A. funestus* more than doubled to reach 73% anchored and a further 17% with chromosome arm assignments (**Fig. 1; Table 2**).

The seven updated assemblies with additional chromosome anchoring data (**Table 2**), together with the chromosome-level *A. gambiae* (PEST) genome, provided the opportunity to confidently assign non-anchored scaffolds and scaffolds from non-anchored assemblies to chromosome arms (**see Methods; Supplementary Online Material: Table S1**). This resulted in total anchoring or arm assignments of 90-92% for the *A. funestus* and *A. stephensi* (SDA-500) assemblies, as well as assignments for the non-anchored assemblies of 96-97% for *A. minimus* and *Anopheles farauti* and 75% or more for a further five assemblies (**Fig. 1**). All of the new improved *Anopheles* genome assemblies and their updated gene annotations, as well as the corresponding chromosome maps of all anchored scaffolds and superscaffolds, have been submitted to VECTORBASE (www.vectorbase.org) for processing and incorporation as new reference assemblies for the benefit of the entire research community.

### Synteny contributions to improved assembly contiguities

Applying only the synteny-based approaches to build two-way consensus sets of well-supported predicted scaffold adjacencies resulted in substantial improvements for several assemblies (**Fig. 2**). These employed orthologues delineated across 21 anopheline gene sets (**Supplementary Online Material: Table S2**) and combined the results from three approaches, ADSEQ (Anselmetti et al. 2018), GOS-ASM (Aganezov and Alekseyev 2016), and ORTHOSTITCH (**see Methods; Supplementary Online Material: Figures S3, S4 and Tables S3, S4**). The two-way consensus adjacencies were required to be predicted by at least two of the approaches with no third-method conflicts. Improvements were quantified in terms of the absolute (**Fig. 2A**) and relative (**Fig. 2B**) increases in scaffold N50 values and decreases in scaffold counts, considering only scaffolds with annotated orthologous genes used as input data for the scaffold adjacency predictions.

**Figure 2.**
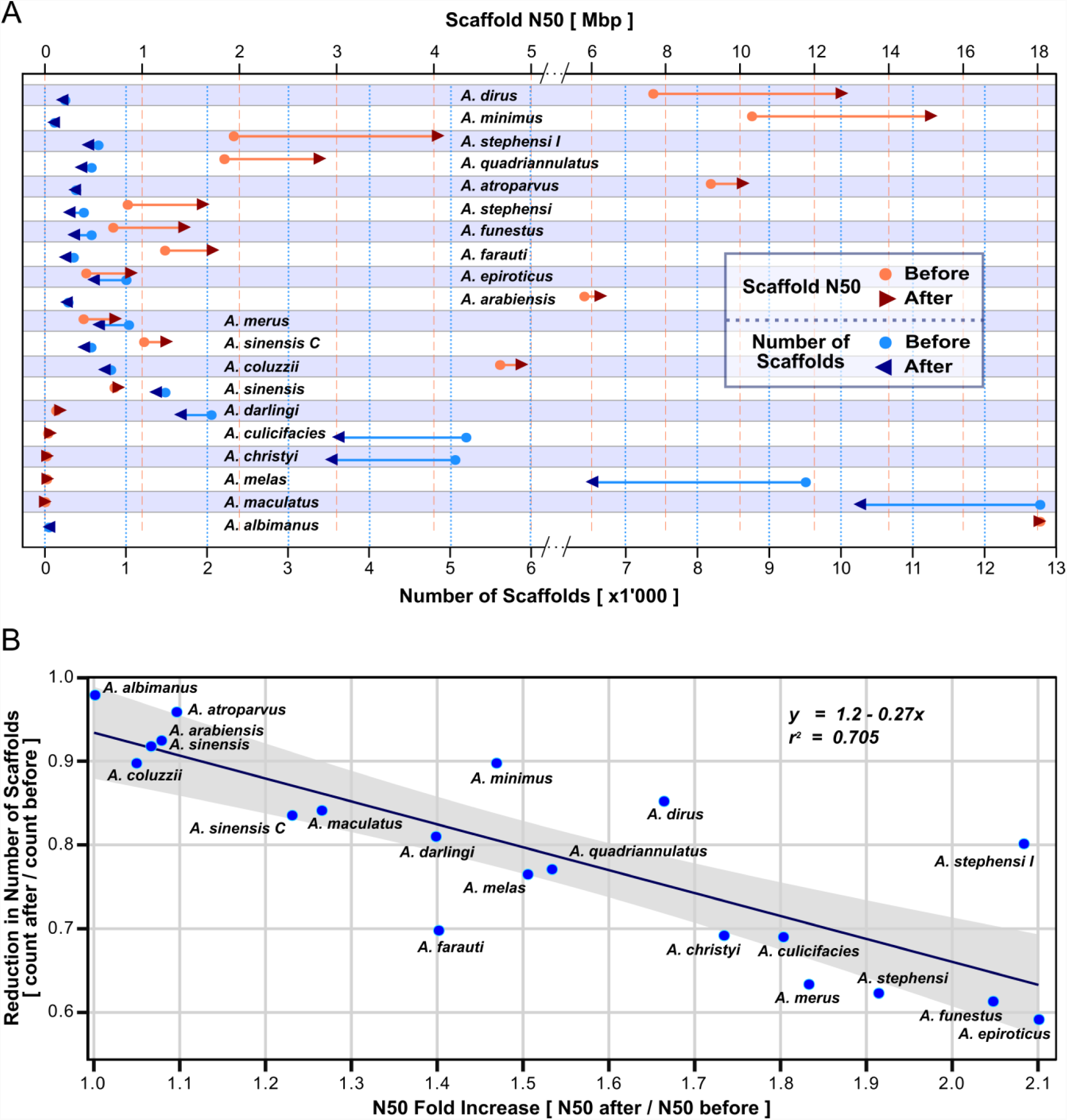
Improved genome assemblies for 20 anophelines from solely synteny-based scaffold adjacency predictions. Results from ADSEQ, GOS-ASM, and ORTHOSTITCH predictions were compared to define two-way consensus adjacencies predicted by at least two of the three approaches, where the third approach did not conflict. These adjacencies were used to build new assemblies with improved contiguities, quantified by comparing before and after scaffold counts and N50 values (half the total assembly length comprises scaffolds of length N50 or longer). The counts, values, and ratios represent only scaffolds with annotated orthologous genes used as the input dataset for the scaffold adjacency predictions. (***A***) Scaffold counts (blues, bottom axis) and N50 values (red/orange, top axis) are shown before (dots) and after (arrowheads) synteny-based improvements were applied. The 20 anopheline assemblies are ordered from the greatest N50 improvement at the top for *Anopheles dirus* to the smallest at the bottom for *Anopheles albimanus*. Note axis scale changes for improved visibility after N50 of 5 Mbp and scaffold count of 6’000. (***B***) Plotting before to after ratios of scaffold counts versus N50 values (counts or N50 after / counts or N50 before superscaffolding of the adjacencies) reveals a general trend of a ∼33% reduction in scaffold numbers resulting in a ∼2-fold increase of N50 values. The line shows the linear regression with a 95% confidence interval in grey. Results for two strains are shown for *Anopheles sinensis*, SINENSIS and Chinese (C), and *Anopheles stephensi*, SDA-500 and Indian (I).

The greatest absolute increases in scaffold N50 values were achieved for *Anopheles dirus* and *A. minimus*, while the greatest absolute reductions in scaffold counts were achieved for *A. christyi, A. culicifacies, A. maculatus*, and *A. melas* (**Fig. 2A**), reflecting the different levels of contiguity of their input assemblies. As no physical mapping data are currently available for these species, and only *A. dirus* and *A. minimus* have supporting RNAseq data, these synteny-based adjacencies represent the only or principal resource from which to build improved assemblies. Reductions in the numbers of scaffolds that comprise each assembly varied from 1’890 fewer for the rather fragmented *A. melas* assembly to just one fewer for the already relatively contiguous *A. albimanus* assembly. Even without large reductions in the numbers of scaffolds, when a few adjacencies bring together relatively long scaffolds then they can lead to marked improvements in N50 values. For example, *A. dirus* and *A. minimus* improved with N50 increases of 5.1 Mbp and 4.8 Mbp and only 36 and 12 fewer scaffolds, respectively.

The general trend indicates that reducing the number of scaffolds by a third leads to a doubling of N50 values (**Fig. 2B**). Exemplifying this trend, *Anopheles epiroticus* showed the greatest relative reduction in the number of scaffolds (40%) and achieved a 2.1-fold N50 increase. Notable exceptions include *A. farauti*, which showed a 1.4-fold N50 increase with a 30% reduction in the number of scaffolds, while *A. dirus* and *A. stephensi* (Indian) achieved 1.66-fold and 2.08-fold N50 increases with only 14% and 19% reductions in the number of scaffolds, respectively. Using only three-way consensus adjacencies led to more conservative improvements, while employing a liberal union of all non-conflicting adjacencies resulted in a trend of a ∼30% scaffold reduction to double N50 values (**Supplementary Online Material: Figures S5, S6**). The enhanced contiguities of these anopheline assemblies based solely on synteny-predicted scaffold adjacencies demonstrate that while the results clearly depend on the quality of the input assemblies, applying synteny-based approaches can achieve substantial improvements.

### Consensus adjacencies from complementary synteny-based methods

To characterise the contributions from each of the synteny-based methods the resulting scaffold adjacency predictions were examined with the Comparative Analysis and Merging of Scaffold Assemblies (CAMSA) tool (Aganezov and Alekseyev 2017) (**Supplementary Online Material: Table S4**). Although each of the computational methods aims to predict scaffold adjacencies based on gene collinearity, they differ in some of their underlying assumptions and in their implementations that identify, score, and infer the most likely scaffold neighbours (**see Methods**). Following traditional meta-assembly-like methods, the comparisons leveraged these differences to identify subsets of well-supported consensus adjacency predictions that were subsequently used for superscaffolding (**Fig. 3**).

**Figure 3.**
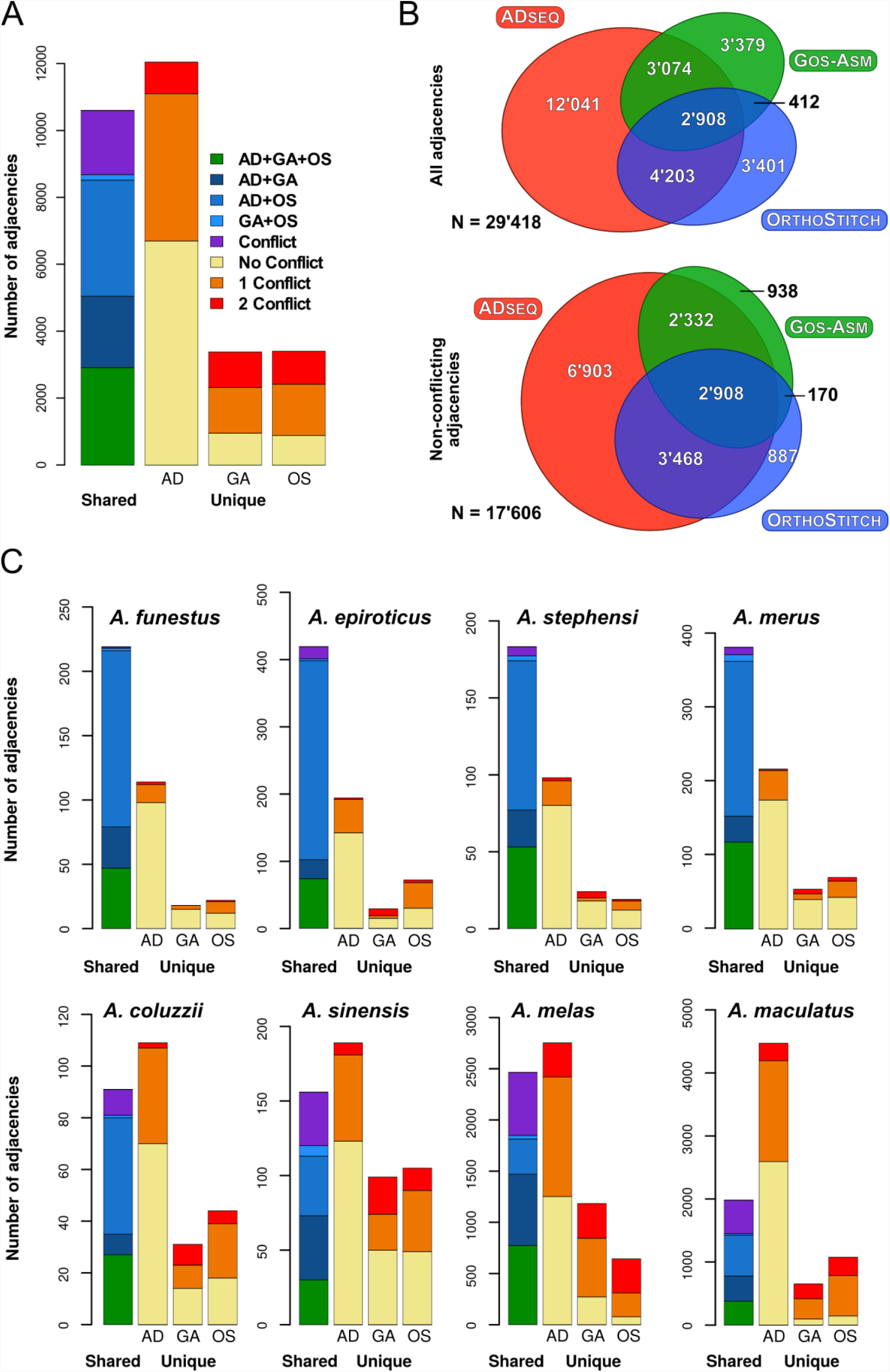
Comparisons of synteny-based scaffold adjacency predictions from ADSEQ (AD), GOS-ASM (GA), and ORTHOSTITCH (OS). Bar charts show counts of predicted adjacencies (pairs of neighbouring scaffolds) that are shared amongst all three methods (green), or two methods without (blues) and with (purple) third method conflicts, or that are unique to a single method and do not conflict (yellow) or do conflict with predictions from one (orange) or both (red) of the other methods. (***A***) Results of all adjacencies summed across all 20 anopheline assemblies. (***B***) Area-proportional Euler diagrams showing (top) the extent of the agreements amongst the three methods for all 29’418 distinct scaffold adjacencies, and (bottom) the extent of the agreements amongst the three methods for the 17’606 distinct and non-conflicting scaffold adjacencies (the liberal union sets), both summed over all 20 assemblies. (***C***) Individual results of adjacencies for representative anopheline assemblies, four with more than 50% agreement (top row), and four with lower levels of agreement (bottom row). Colours for each fraction are the same as in panel A, y-axes vary for each assembly with maxima of 120 for *Anopheles coluzzii* to 5’000 for *Anopheles maculatus*. Results for *Anopheles stephensi* are for the SDA-500 strain.

For the full set of assemblies, GOS-ASM and ORTHOSTITCH predicted about 10’000 oriented adjacencies each, with just over twice as many predictions from ADSEQ. Comparing all predictions identified almost 30’000 distinct scaffold adjacencies, 36% of which were supported by at least two methods; this fraction comprised 10% that were in three-way agreement and a further 20% that were in two-way agreement with no conflicts with the third method (**Fig. 3; Supplementary Online Material: Figure S7**). The larger sets of predictions from ADSEQ resulted in much higher proportions of unique adjacencies. Adjacencies in three-way agreement constituted just under a third of GOS-ASM and ORTHOSTITCH predictions, and just 13% of the more numerous ADSEQ predictions. From the liberal union sets of all non-conflicting adjacencies for all assemblies, the adjacencies in three-way agreement made up a sixth of the total (**Fig. 3B**). Considering only the two-way consensus sets of adjacencies used for the synteny-based assembly improvements, the three-way consensus adjacencies increased to a third of the total, and 54% of GOS-ASM, 44% of ORTHOSTITCH, and 33% of ADSEQ predictions. Of these two-way consensus adjacencies 98% were supported by ADSEQ, 74% by ORTHOSTITCH, and 61% by GOS-ASM. Thus, comparing the results from the three methods and employing a two-way agreement with no third-method conflict filter resulted in greatly improved levels of adjacency agreements.

For the individual assemblies, more than half of the distinct scaffold adjacencies were in agreement for *A. epiroticus, Anopheles merus*, and both the *A. stephensi* assemblies, with *A. funestus* achieving the highest consistency at 58% (**Fig. 3C; Supplementary Online Material: Figure S8**). Some of the most fragmented input assemblies produced some of the largest sets of distinct adjacency predictions but the agreement amongst these predictions was generally lower than the other assemblies. For example, *A. maculatus* was the least contiguous input assembly and produced more than 8’000 distinct predictions, of which only 18% showed at least two-way agreement with no conflicts (**Fig. 3C; Supplementary Online Material: Figure S8**).

### Enhanced superscaffolding with physical mapping and RNA sequencing data

Combining the synteny-based results with physical mapping data from a subset of the anophelines allowed for enhanced superscaffolding as well as independent validations of the synteny-based predictions and their consensus sets. Building cytogenetic photomaps and conducting extensive FISH experiments mapped 31 *A. albimanus* scaffolds (Artemov et al. 2017), 46 *A. atroparvus* scaffolds (Artemov et al. 2015; Neafsey et al. 2015; Artemov et al. 2018a), 202 *A. funestus* scaffolds (Sharakhov et al. 2002, 2004; Xia et al. 2010; Neafsey et al. 2015) (including additional mapping for this study), 52 *A. sinensis* scaffolds (Chinese) (Wei et al. 2017), 99 *A. stephensi* (SDA-500) scaffolds (Neafsey et al. 2015), and 118 *A. stephensi* (Indian) scaffolds (Jiang et al. 2014) (including additional mapping for this study) (**see Methods; Supplementary Online Material: Figure S9 and Tables S5, S6**). The scaffold adjacencies identified from these physical mapping data, i.e. pairs of neighbouring mapped scaffolds, were compared with adjacencies predicted by each of the three methods and the CAMSA-generated consensus sets (**Supplementary Online Material: Table S7**). *A. funestus* validations confirmed 12-17% of the different sets of synteny-based adjacencies and highlighted conflicts with just 4-8%, while for *A. atroparvus* five of the 15 two-way consensus synteny-based predictions were confirmed by physical mapping and only one conflict was identified (**Fig. 4A**). Examining the identified conflicts in detail revealed that most were resolvable. As not all scaffolds were targeted for physical mapping, neighbouring scaffolds on the physical maps could have shorter unmapped scaffolds between them that were identified by the synteny-based approaches. For *A. funestus*, five conflicts were resolved because the synteny-based neighbour was short and not used for physical mapping and an additional four conflicts were resolved by switching the orientation of physically mapped scaffolds, which were anchored by only a single FISH probe and therefore their orientations had not been confidently determined.

**Figure 4.**
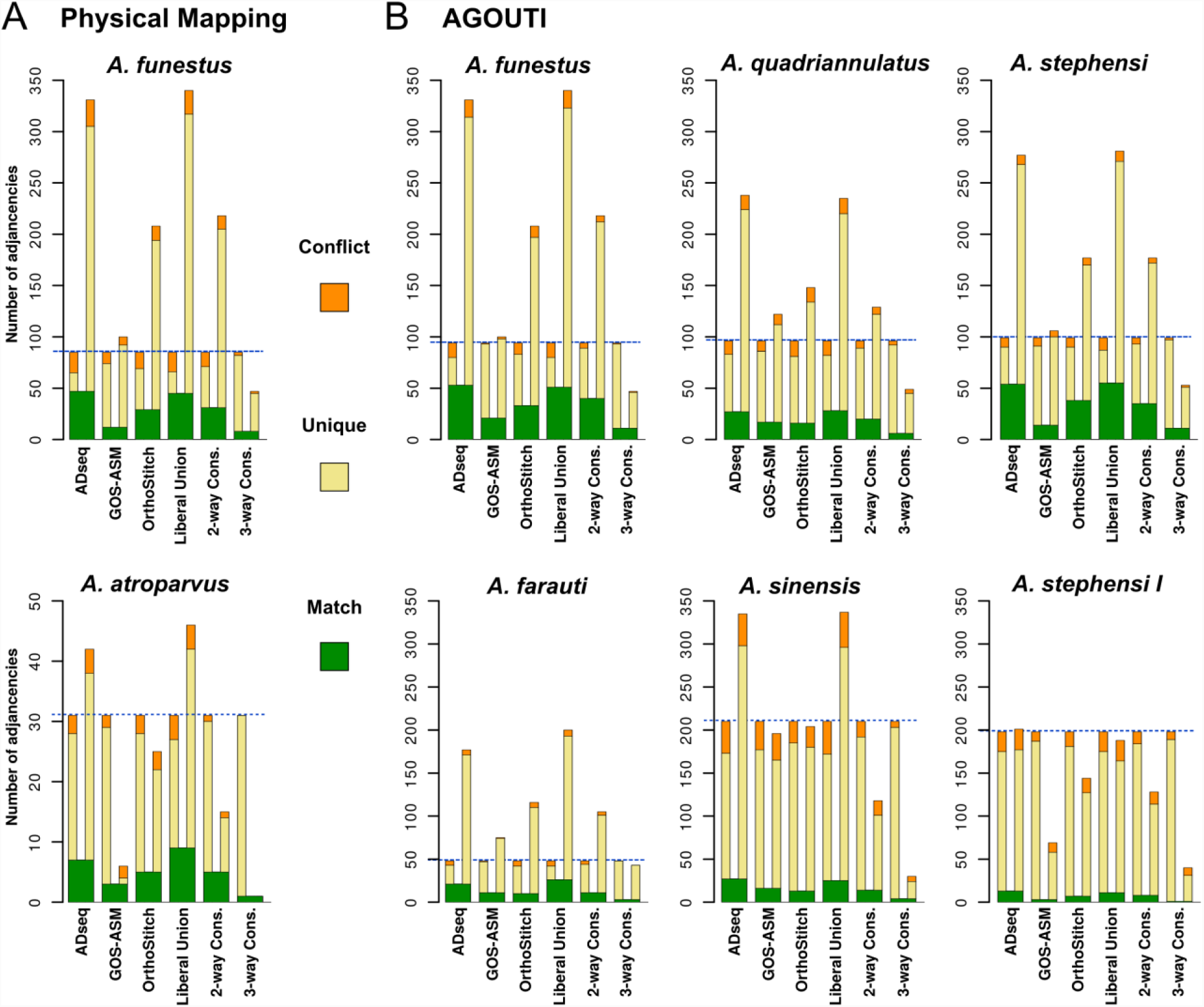
Comparisons of synteny-based scaffold adjacency predictions with physical mapping and RNA sequencing data. The bar charts show counts from each set of synteny-based scaffold adjacency predictions compared with the adjacencies from the physical mapping (***A***) or RNAseq AGOUTI-based (***B***) sets. The synteny-based sets comprise predictions from three different methods, ADSEQ, GOS-ASM, and ORTHOSTITCH, as well as their Liberal Union (all non-conflicting predictions), their two-way consensus (2-way Cons. predicted by two methods and not conflicting with the third method), and their three-way consensus (3-way Cons. predicted by all three methods). Adjacencies that are exactly matching form the green base common to both sets in each comparison, from which extend bars showing physical mapping or AGOUTI adjacency counts (left) and synteny-based adjacency counts (right) that are unique (yellow) or conflicting (orange) in each comparison. Blue dashed lines highlight the total adjacencies for the physical mapping or AGOUTI sets. For comparison all y-axes are fixed at a maximum of 350 adjacencies, except for *Anopheles atroparvus*. Results for two strains are shown for *Anopheles stephensi*, SDA-500 and Indian (I).

Transcriptome data from RNAseq experiments enabled further superscaffolding and validations of the synteny-based predictions and their consensus sets. The Annotated Genome Optimization Using Transcriptome Information (AGOUTI) tool (Zhang et al. 2016) employs RNAseq data to identify adjacencies when individual transcripts (or paired-end reads) reliably map to scaffold extremities. Using available mapped paired-end RNAseq data from VECTORBASE (Giraldo-Calderón et al. 2015), scaffold adjacencies predicted for 13 anophelines ranged from just two for *A. albimanus* to 210 for *A. sinensis* (SINENSIS) (**Supplementary Online Material: Table S8**). These AGOUTI-based scaffold adjacencies were compared with the adjacencies predicted by each of the three methods and the CAMSA-generated consensus sets (**Fig. 4B; Supplementary Online Material: Table S9**). Across all 13 assemblies, 18% of AGOUTI-based scaffold adjacencies supported the two-way consensus synteny-based adjacencies, 75% were unique to the AGOUTI sets, and only 7% were in conflict. Nearly 200 AGOUTI-based scaffold adjacencies for *A. stephensi* (Indian) confirmed only eight and conflicted with 14 of the two-way consensus set adjacencies (**Fig. 4B**). In contrast, about half as many AGOUTI-based scaffold adjacencies each for *A. stephensi* (SDA-500) and *A. funestus* confirmed four to five times as many two-way consensus set adjacencies and conflicted with only five and six, respectively. Notably, 68% of the AGOUTI-based scaffold adjacencies that produced conflicts with the two-way consensus set adjacencies comprised scaffolds with no annotated orthologues. These cases can be resolved by noting that only scaffolds with orthologous genes were used for synteny-based predictions: therefore, the inferred neighbouring scaffolds could have shorter non-annotated scaffolds between them that were identified by AGOUTI. Such non-annotated scaffolds were also numerous amongst the adjacencies that were unique to AGOUTI where for 66% either one or both scaffolds had no annotated orthologues.

### Superscaffold comparisons with new genome assemblies

A new *A. funestus* assembly, designated AfunF2-IP, was generated as part of this study by merging approximately 70X of PacBio sequencing data with the reference assembly (AfunF1), with subsequent scaffolding using the original Illumina sequencing data (**see Methods; Supplementary Online Material: Figure S10 and Table S10**). This AfunF2-IP assembly for *A. funestus* enabled the validation of the scaffold adjacency predictions for the AfunF1 assembly by examining collinearity between the two assemblies. AfunF1 scaffolds were ordered and oriented based on their alignments to AfunF2-IP scaffolds and the resulting 321 alignment-based scaffold adjacencies were then compared with the synteny-based and AGOUTI predictions as well as with the physical mapping adjacencies to identify supported, unique, and conflicting adjacencies (**Fig. 5; Supplementary Online Material: Figure S11 and Table S11**). Each of the three synteny method prediction sets, as well as the two-way consensus and liberal union sets, had 14-17.5% in common with the alignment-based scaffold adjacencies, fewer than a quarter in conflict, and almost two thirds that were neither supported nor in conflict (**Supplementary Online Material: Table S11**). The physical mapping adjacencies had generally more support, but also more conflicts as about half disagreed with the alignment-based adjacencies. Several disagreements were easily resolved by comparing these conflicts with those identified from the synteny-based adjacencies and confirming that switching the orientation of physically mapped scaffolds corrected the relative placements of these scaffolds, e.g. **Fig. 5 inset (i)**. Similarly to the comparisons with the physical mapping and RNAseq data presented above, apparent conflicts with the alignment-based adjacencies can also arise because using genome alignment data considered all alignable scaffolds while physical mapping targeted only large scaffolds and synteny methods did not consider scaffolds with no annotated orthologues (i.e. short scaffolds). This is exemplified in **Fig. 5 inset (ii)** where the alignment data placed a short scaffold between two scaffolds predicted to be neighbours by ADSEQ, ORTHOSTITCH, and physical mapping data. Skipping such short scaffolds (<5 Kbp) to define a smaller set of alignment-based adjacencies considering only the longer scaffolds resulted in increased support for the synteny-based sets of 19-23%, and most notably up to 39% for the physical mapping adjacencies, while only marginally increasing support for AGOUTI predictions from 15% to 17% (**Supplementary Online Material: Table S11**).

**Figure 5.**
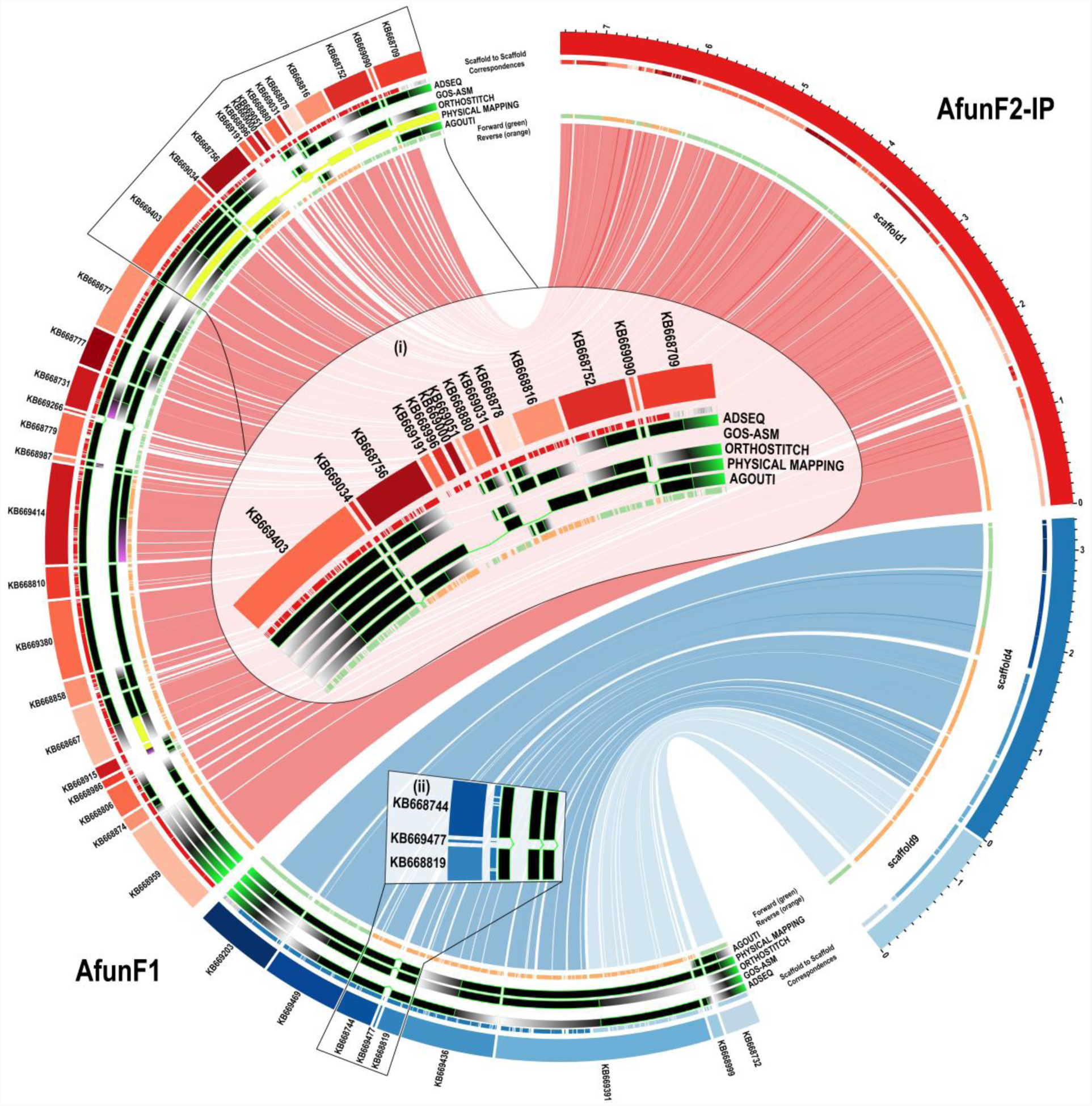
Whole genome alignment comparisons of selected *Anopheles funestus* AfunF1 and AfunF2-IP scaffolds. The plot shows correspondences of three AfunF2-IP scaffolds (right) with AfunF1 (left) scaffolds based on whole genome alignments, with links coloured according to their AfunF2-IP scaffold. Putative adjacencies between AfunF1 scaffolds are highlighted with tracks showing confirmed neighbours (black with bright green borders), supported neighbours with conflicting orientations (yellow), scaffolds with putative adjacencies that conflict with the alignments (purple gradient), scaffolds without putative adjacencies and thus no conflicts with the alignments (grey gradient) for: from outer to inner tracks, ADSEQ, GOS-ASM, ORTHOSTITCH, physical mapping, and AGOUTI. The innermost track shows alignments in forward (green) and reverse (orange) orientations. The outermost track shows alignments coloured according to the corresponding scaffold in the other assembly (light grey if aligned to scaffolds not shown). **Inset (i)** shows how corrected orientations of physically mapped scaffolds agree with the other methods. **Inset (ii)** shows how the alignments identified a short scaffold that was placed between two scaffolds identified by three other methods.

Re-scaffolding of the initial *A. farauti* (AfarF1) and *A. merus* (AmerM1) assemblies employed large-insert ‘Fosill’ sequencing libraries and reduced the numbers of scaffolds from 550 to 310 and 2’753 to 2’027 and increased N50 values from 1’197 Kbp to 12’895 Kbp and 342 Kbp to 1’490 Kbp, respectively (Neafsey et al. 2015). The availability of these re-scaffolded assemblies enabled the validation of the synteny-based and AGOUTI-based scaffold adjacency predictions for the AfarF1 and AmerM1 assemblies by examining corresponding scaffolds from the AfarF2 and AmerM2 assemblies (**see Methods; Supplementary Online Material: Figure S12**). The comparisons identified full support for the majority (87% and 82%) of the two-way synteny consensus set adjacencies and unresolvable conflicts for just 5% and 10%, while the AGOUTI-based adjacencies achieved similarly high levels of full support (81% and 67%), but with slightly greater proportions of conflicts (**Supplementary Online Material: Table S12**).

### Updated cytogenetic photomaps and physical genome maps for *A. funestus* and *A. stephensi*

The collated data allowed for comprehensive updates of the previously published chromosomal photomaps from ovarian nurse cells for *A. funestus* (Sharakhov et al. 2002) and for *A. stephensi* (Sharakhova et al. 2006). The existing images of *A. funestus* polytene chromosomes of the five arms common to all anophelines (X, 2R, 2L, 3R, and 3L) were further straightened to facilitate linear placements of the genomic scaffolds on the photomap (**Fig. 6**). Major structural updates to the *A. funestus* cytogenetic photomap included reversal of the order of divisions and subdivisions within the 3La inversion to follow the standard 3L+^a^ arrangement, and merging of two small subdivisions with larger neighbouring subdivisions: 5D to 6 and 34D to 34C. The previous physical genome map of the AfunF1 assembly included 104 scaffolds and spanned 35% of the assembly (Neafsey et al. 2015). The extensive additional physical mapping performed for *A. funestus*, together with the new AfunF2-IP assembly and sequence alignment-based comparisons with the AfunF1 assembly, enabled an updated physical genome map to be built (**Fig. 6**). The 126 previously FISH-mapped (Sharakhov et al. 2002, 2004; Xia et al. 2010) and 66 newly FISH-mapped DNA markers (**Supplementary Online Material: Figure S9**) were located with BLAST searches to 139 AfunF1 scaffolds and then compared with AfunF2-IP scaffolds using whole genome pairwise alignments (**see Methods**). The placement of scaffolds along the photomap took advantage of comparisons with the synteny-based scaffold adjacency predictions and with the AfunF1-AfunF2-IP whole genome pairwise alignments. Synteny-or alignment-based scaffold neighbours were added to the genome map when they were short and thus had not been used for physical mapping. Additionally, scaffolds which were anchored with only a single FISH probe (i.e. with undetermined orientations) were reoriented when synteny-or alignment-based scaffold adjacencies provided supporting evidence to correct their relative placements on the map. The resulting physical genome map for *A. funestus* includes 202 AfunF1 scaffolds spanning 61% of the assembly (**Supplementary Online Material: Table S6**), with a further 100 neighbouring scaffolds (additional 12% of the assembly) after incorporating the synteny-based and AGOUTI-based adjacencies. For *A. stephensi* (Indian), structural updates to the cytogenetic photomap (Sharakhova et al. 2006) included changing the order of lettered subdivisions on arms 2L and 3L to match the order of numbered divisions (**Fig. 7**). The previous physical genome map of the AsteI2 assembly included 86 scaffolds and spanned 62% of the assembly (Jiang et al. 2014). The additional FISH probes allowed for 43 scaffolds to be oriented and placed a total of 118 scaffolds on the cytogenetic photomap spanning 79% of the assembly (**Fig. 7**) with a further 90 neighbouring scaffolds (additional 5% of the assembly) after incorporating all reconciled adjacencies.

**Figure 6.**
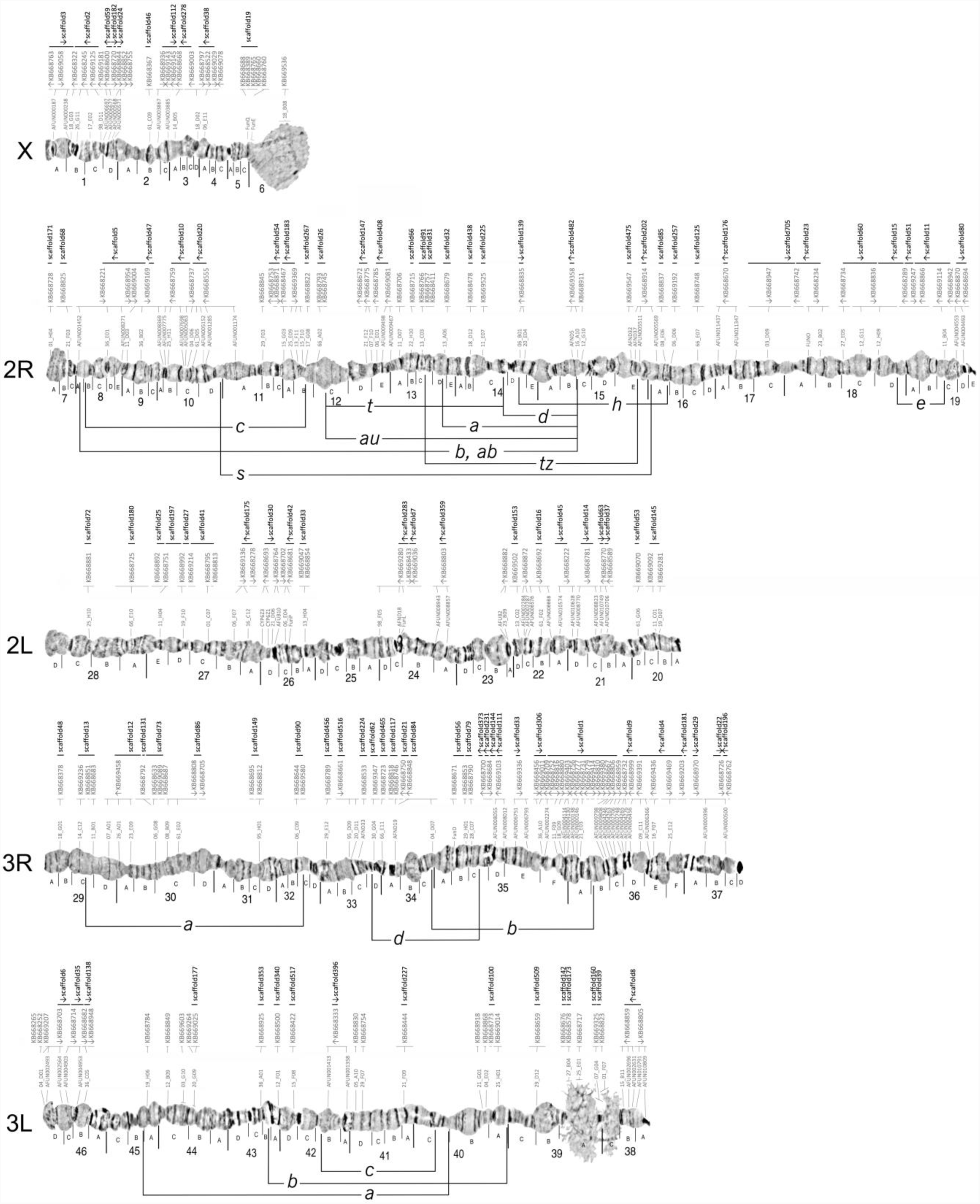
The *Anopheles funestus* cytogenetic photomap of polytene chromosomes with anchored scaffolds from the AfunF1 and AfunF2-IP assemblies. FISH-mapped DNA markers (grey probe identifiers directly above each chromosome) show the density of physical mapping along the chromosome arm subdivisions (labelled with letters A, B, C, etc. directly below each chromosome) and divisions (labelled with numbers 1-46 below the subdivision labels). Scaffolds from the AfunF1 (KB66XXXX identifiers, grey font and thin horizontal lines) and AfunF2-IP (scaffoldXX identifiers, black font and thick horizontal lines) assemblies are ordered along the photomap above each chromosome. Orientation of the scaffolds in the genome, if known, is shown by the arrows below each of the scaffold identifiers. Known polymorphic inversions are shown for chromosome arms 2R, 3R, and 3L.

**Figure 7.**
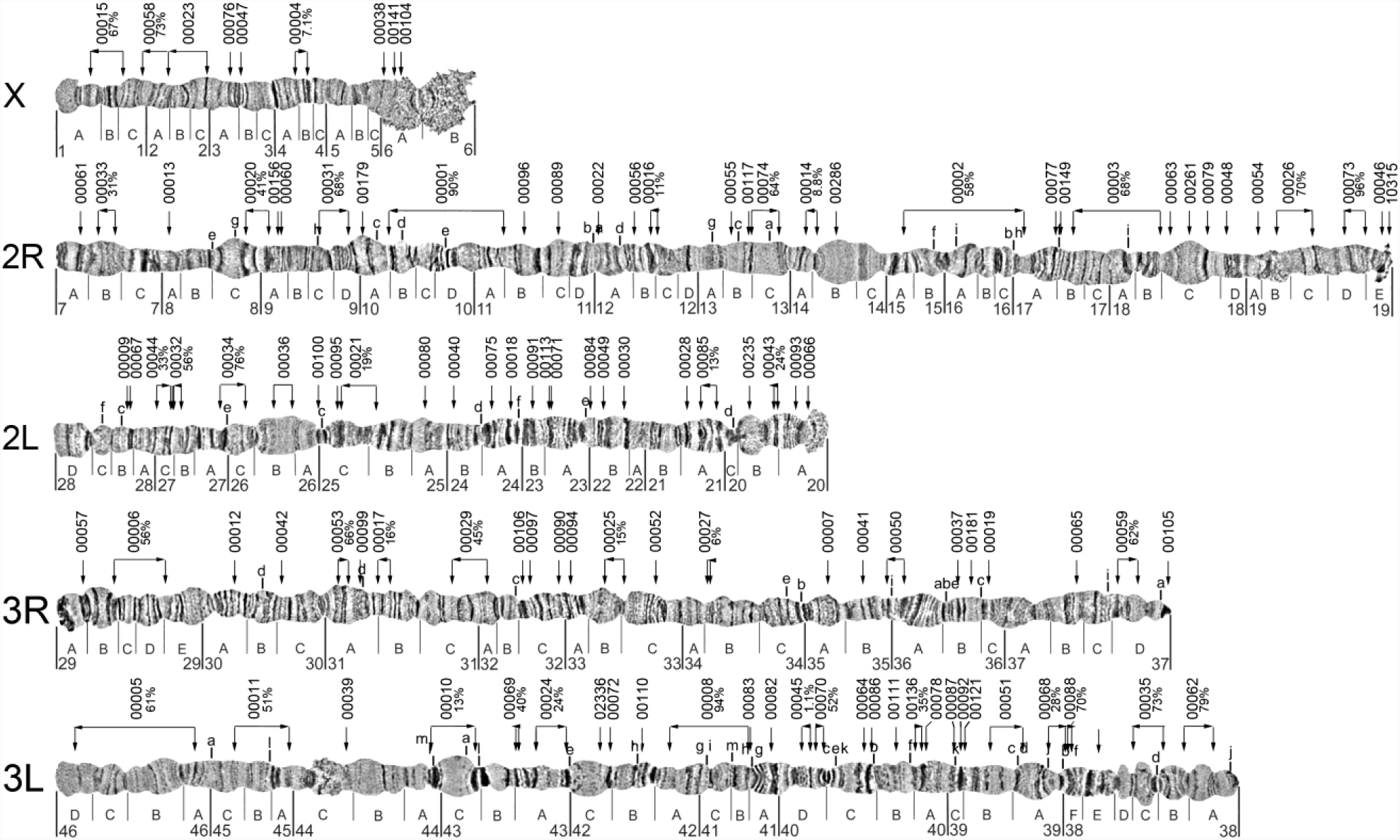
The *Anopheles stephensi* cytogenetic photomap of polytene chromosomes with anchored scaffolds from the AsteI2 assembly. The updated cytogenetic photomap is shown with chromosome arm subdivisions (labelled with letters A, B, C, etc. directly below each chromosome) and divisions (labelled with numbers 1-46 below the subdivision labels). Locations of known polymorphic inversions are indicated with lowercase letters above chromosome arms 2R, 2L, 3R, and 3L. The AsteI2 assembly identifiers of the 118 mapped scaffolds are shown above each chromosome arm (scaffold identifiers are abbreviated e.g. ‘scaffold_00001’ is shown on the map as ‘00001’), and the locations of FISH probes used to map the scaffolds are shown with downward-pointing arrows. For scaffolds with two mapped FISH probes, the orientations along the genome map are shown with horizontal arrows below each of the scaffold identifiers, with labels indicating the proportion (%) of each scaffold located between the probe pairs.

## Discussion

Integrating synteny-based scaffold adjacency predictions with additional supporting data for subsets of the anophelines enabled superscaffolding with chromosome anchoring and arm assignments to produce 20 new *Anopheles* reference genome assemblies (**Fig. 1; Tables 1 and 2**). Consensus predictions were used to build the improved assemblies for which the general trend showed that a reduction in the total number of orthologue-bearing scaffolds of about a third could double the scaffold N50 (**Fig. 2**). Notably, when the scaffolds involved were long, even a handful of adjacencies could greatly increase the N50 value, however, the numerous adjacencies for the rather fragmented input assemblies improved their contiguity but led to only minor improvements in N50 values. For the six assemblies with starting N50 values of between 340 Kbp and 840 Kbp (considering all scaffolds, not only those with orthologues), the average improvement was just under 400 Kbp, demonstrating what can be achieved using only synteny-based approaches. By way of comparison, the honeybee genome assembly upgrade relied on millions of reads from ∼20X SOLiD and ∼5X Roche 454 sequencing to improve the scaffold N50 by 638 Kbp from 359 Kbp to 997 Kbp (Elsik et al. 2014). Thus, while the *Anopheles* results varied considerably depending on the input assemblies, using only gene synteny-based adjacencies from a combined analysis of the results of three methods achieved substantial contiguity improvements for many assemblies.

The comparisons of predicted adjacencies from the three synteny-based methods (**Fig. 3**) highlight the challenge of inferring accurate adjacencies as well as the importance of employing multiple approaches. Only 10% of all distinct scaffold adjacencies were predicted by all three methods, but building the two-way consensus sets increased this three-method agreement more than three-fold, and almost all the two-way consensus adjacencies were supported by ADSEQ, nearly three-quarters by ORTHOSTITCH, and three-fifths by GOS-ASM. Consensus building can therefore take advantage of differences amongst the employed methods to achieve the goal of identifying a subset of well-supported adjacencies. Synteny block delineation, which then allows for scaffold adjacencies to be predicted, is itself a complex task where results from different anchor-based approaches can vary considerably (Liu et al. 2018a). Several key differences distinguish the three methods applied to the *Anopheles* assemblies, for example, GOS-ASM employs only single-copy orthologues so any gene duplications are excluded from the ancestral genome reconstructions, whereas the other two methods do consider paralogues. Furthermore, both GOS-ASM and ADSEQ are ‘phylogeny-aware’ algorithms as they use the species tree topology, and ADSEQ additionally employs individual gene trees for each orthologous group. In contrast, ORTHOSTITCH does not take phylogenies into account and instead relies on enumerating levels of support across the dataset to score putative adjacencies. These differences affect the sensitivity and specificity of the methods, reflected by the more numerous predictions from ADSEQ that can explore complex gene evolutionary histories within the species tree topology, versus the smaller sets of adjacencies from GOS-ASM, which excludes complexities introduced by gene duplications, and ORTHOSTITCH that simplifies the search by not imposing any evolutionary model. So while the consensus approach applied to the predictions across all the anophelines results in reduced sensitivities, it takes advantage of the different underlying assumptions and algorithmic implementations of each method to identify well-supported sets of scaffold adjacencies.

The input data are another factor that may influence the number of predicted adjacencies, the level of agreement amongst different methods, and the achievable contiguity improvements. An assembly with many short scaffolds with annotated orthologues may achieve numerous adjacency predictions, e.g. *A. maculatus* with 1’454 consensus adjacencies, but an assembly with such low contiguity is less likely to provide support for putative adjacencies in other assemblies. The evolutionary divergence of the set of species, as well as the total number of species, to which these methods are applied would also impact their ability to recover reliable adjacencies, because the complexity of the task of inferring synteny blocks is greatly reduced if the input orthology dataset consists mainly of near-universal single-copy orthologues. As gene duplications and losses accumulate over time the proportion of near-universal single-copy orthologues will shrink, and even amongst those that are maintained translocations and genomic shuffling events will add to the steady erosion of the evolutionary signals on which these methods rely. Rearrangements may also be more or less common in different genomic contexts, e.g. the *Osiris* (Shah et al. 2012) and *TipE* (Li et al. 2011) gene clusters have been noted for their unusually high synteny conservation across insects, or in different species, e.g. the well-known *Hox* gene cluster is largely collinear across animals but may be found with disorganized, split, or atomised arrangements (Duboule 2007). Genomic shuffling rates may also vary among different lineages—e.g. lepidopteran genomes appear to have reduced levels of gene rearrangements (Kanost et al. 2016)—so seemingly equally divergent (in terms of time to last common ancestor) sets of species may be differentially amenable to superscaffolding through synteny delineation.

The availability of alternative datasets with scaffold adjacency information for subsets of the anophelines allowed for several comparisons with the predictions based solely on synteny inferences to be performed. Although generally few adjacencies were obtained from the physical mapping data (because only larger scaffolds were selected for mapping they may not be direct neighbours) the comparisons were able to identify support for many synteny-based adjacencies (**Fig. 4A**). Several conflicts were also identified; however, most of these were due to the fact that the synteny-based neighbour was a short scaffold that had not been targeted for physical mapping and could be positioned between the two much larger physically mapped scaffolds, thus, they are not truly conflicts. Importantly, other conflicts involved only the relative orientation of neighbouring scaffolds and occurred with scaffolds that were anchored with only a single FISH probe and whose orientations had thus not been confidently determined. In these cases the synteny-based adjacencies therefore provided key complementary information and helped to correct the orientations of the physically mapped scaffolds. Comparisons with RNAseq-based AGOUTI-predicted adjacencies also provided support for many synteny-based predictions (**Fig. 4B**). Two-thirds of the adjacencies unique to AGOUTI were between scaffolds where one or both had no annotated orthologues. As AGOUTI is not restricted to large scaffolds preferred for physical mapping or scaffolds with annotated orthologues required for synteny-based approaches, it can provide complementary predictions that capture shorter non-annotated scaffolds that would otherwise not be recovered. While this would not substantially improve N50 values it is nonetheless important for improving gene annotations as correcting such assembly breaks could allow for more complete gene models to be correctly identified.

The *A. funestus* PacBio-based AfunF2-IP assembly scaffolds facilitated the alignment-based ordering and orientation of AfunF1 scaffolds for comparisons with the adjacency predictions and physical mapping data (**Fig. 5**). These supported up to almost a quarter of *A. funestus* two-way consensus synteny adjacencies and about 40% of the physical mapping adjacencies. Importantly, most were neither supported nor in conflict, and conflicts generally occurred when the alignment-based adjacencies included short scaffolds that were not considered by the synteny-based or physical mapping approaches, and thus could be resolved. For *A. farauti* and *A. merus*, the genome-alignment-based comparisons of their initial assemblies with the re-scaffolded AfarF2 and AmerM2 assemblies provided much higher levels of support for the two-way consensus synteny adjacencies, with very few conflicts. This reflects the radically different approaches between re-scaffolding, where the additional ‘Fosill’ library data served to build longer scaffolds from the initial scaffolds, versus the Illumina-PacBio hybrid re-assembly of *A. funestus*. These comparisons therefore validate many of the synteny-based adjacency predictions while conceding that short intervening scaffolds may be overlooked due to the limitations of having to rely on scaffolds with annotated orthologues.

As modern long-read and long-range sequencing technologies are capable of producing highly contiguous assemblies (Saha 2019), it is conceivable that many fragmented draft genomes will be completely superseded by new independently built high-quality reference assemblies. For example, single-molecule sequencing technologies were recently employed to produce assemblies of 15 *Drosophila* species, 14 of which already had previously reported sequenced genomes (Miller et al. 2018), and a high-quality *de novo* assembly from a single *A. coluzzii* mosquito (Kingan et al. 2019). Alternatively, re-sequencing to obtain proximity data to use in conjunction with contigs from draft assemblies can also achieve high-quality references to replace the fragmented initial versions, e.g. (Putnam et al. 2016; Dudchenko et al. 2017).

Furthermore, although reference-assisted assembly approaches may mask true genomic rearrangements (Liu et al. 2018a), high-quality chromosome-level genomes of very close relatives can be used to improve draft assemblies, often employing alignment-based comparisons such as assisted assembly tools (Gnerre et al. 2009), reference-assisted chromosome assembly (Kim et al. 2013), CHROMOSOMER (Tamazian et al. 2016), the Reference-based Genome Assembly and Annotation Tool (Liu et al. 2018b), or the RAGOUT 2 reference-assisted assembly tool (Kolmogorov et al. 2018). What role then is there for comparative genomics approaches that use evolutionary signals to predict scaffold adjacencies in draft assemblies?

Firstly, while recognising that downward trending costs of many new technologies are making sequencing-based approaches more accessible to even the smallest of research communities, the costs and time associated with experimental finishing or re-sequencing efforts remain non-trivial and acquired expertise is needed for high-quality sample preparation and library building. Furthermore, the disappointing reality is that re-sequencing and re-scaffolding does not always lead to vastly improved assemblies, albeit an anecdotal reality because failures are not reported in the published literature. Secondly, hybrid assembly approaches benefit from the complementarity of the different types of input data that they employ, and our comparisons show that synteny-based adjacencies can further complement the experimental data. In this regard, even if synteny-based results are not directly included in such hybrid approaches, they can nevertheless serve as a benchmark against which to quantify the effectiveness of different combinations of approaches (or different parameters used) and help guide re-assembly procedures towards producing the best possible improved assemblies. Thirdly, reference-assisted assembly approaches work best with good quality closely related reference and outgroup genomes, which are not always available. The anophelines analysed here shared a common ancestor some 100 million years ago and only about 9% of the *A. gambiae* (PEST) genome was alignable to the most distant relatives (Neafsey et al. 2015). Previous comparisons of *Ae. aegypti* and *A. gambiae* revealed that almost 80% of their single-copy orthologues were retained in the same genomic neighbourhood (Waterhouse et al. 2008), and using protein sequence alignments identifies recognisable orthologues for about 80% of genes between the most distant pairs of anophelines. Multi-species gene synteny-based approaches are therefore well-suited to the analysis of datasets such as the 21 *Anopheles* assemblies. Finally, our results show how physical mapping datasets can be augmented or even corrected through comparisons with synteny-based scaffold adjacency predictions. Where subsets of scaffolds have already been mapped to chromosomes (**Figs. 6 and 7; Table 2**), adding neighbouring scaffolds from synteny-based predictions can add to the overall total proportion anchored without more labour-intensive experimental work. Superscaffolding also reduces the total numbers of scaffolds to be mapped and thus allows for greater proportions of draft assemblies to be anchored using fewer markers. Comprehensive anchoring in multiple species in turn allows for greater confidence from cross-species comparisons to assign non-anchored scaffolds to chromosome arms. Chromosome anchoring and arm assignments have facilitated investigations such as rates of gene translocations between chromosome arms (Neafsey et al. 2015), the genetics of saltwater tolerance (Smith et al. 2015) or resting behaviour and host preference (Main et al. 2016), chromosome arm-specific patterns of polymorphism (Kamdem et al. 2017), sex-biased gene expression (Papa et al. 2017), dosage compensation (Deitz et al. 2018), or the evolution of sex chromosomes (Pease and Hahn 2012; Vicoso and Bachtrog 2015). The new assemblies with enhanced chromosome mapping achieved for the anophelines therefore represent greatly improved genomic resources for future studies.

## Conclusions

Our three-method synteny-based scaffold adjacency prediction workflow is relatively easily implemented and may flexibly include results from additional adjacency predictors or, as evidenced with our various types of comparison datasets, alternative sources of adjacency information. Rather than prescribing a panacea to cure all assembly ailments, we conclude that the components of this workflow may be adapted, substituted, extended, or simplified according to the needs and resources of draft genome assembly improvement projects. Assessing the performance of three comparative genomics approaches and comparing their results with available experimental data demonstrates their utility as part of assembly improvement initiatives, as well as highlighting their complementarity to experimental approaches. The consensus predicted scaffold adjacencies can lead to substantial improvements of draft assemblies without requiring additional sequencing-based support, and they can add to and improve physical mapping efforts and chromosome arm assignments. These evolutionarily guided methods therefore augment the capabilities of any genome assembly toolbox with approaches to assembly improvements or validations that will help to propel the draft assemblies from similar species-clusters along the journey towards becoming ‘finished’ reference genomes.

## Methods

### Synteny-based scaffold adjacency predictions

The synteny-based prediction tools require as input both delineated orthology and genomic location data for the annotated genes from each assembly. All gene annotations were retrieved from VECTORBASE (Giraldo-Calderón et al. 2015) and orthology data were retrieved from ORTHODB V9 (Zdobnov et al. 2017): versions of the genome assemblies and their annotated gene sets are detailed in **Supplementary Online Material: Table S2**, along with counts of scaffolds, genes, and orthologues. With an average of 11’832 orthologues (standard deviation 1’075), including 10’708 orthologous groups with genes from more than half of the 21 anophelines, these data provide a comprehensive set of genomic markers for gene synteny-based approaches. ADSEQ analysis first builds reconciled gene trees for each orthologous group (gene family), then for pairs of gene families for which extant genomic adjacencies are observed, or suggested by sequencing data, a duplication-aware parsimonious evolutionary scenario is computed, via Dynamic Programming (DP), that also predicts extant adjacencies between genes at the extremities of contigs or scaffolds. This DP algorithm also accounts for scaffolding scores obtained from paired-end reads mapped onto contigs and provides a probabilistic score for each predicted extant adjacency, based on sampling optimal solutions (Anselmetti et al. 2018). ADSEQ was applied across the full anopheline input dataset to predict scaffold adjacencies (**Supplementary Online Material: Table S3**). GOS-ASM (Gene order scaffold assembler) employs an evolutionary rearrangement analysis strategy on multiple genomes utilizing the topology of the species phylogenetic tree and the concept of the breakpoint graph (Aganezov and Alekseyev 2016). Fragmented genomes with missing assembly ‘links’ between assembled regions are modelled as resulting from artificial ‘fissions’ caused by technological fragmentation that breaks longer contiguous genomic regions (chromosomes) into scaffolds (Aganezov et al. 2015). Assembling these scaffolds is therefore reduced to a search for technological ‘fusions’ that revert non-evolutionary ‘fissions’ and glue scaffolds back into chromosomes. GOS-ASM was applied to the full anopheline input dataset to predict such scaffold ‘fusions’ (**Supplementary Online Material: Table S3**). The ORTHOSTITCH approach was first prototyped as part of the investigation of greater synteny conservation in lepidopteran genomes (Kanost et al. 2016), and subsequently further developed as part of this study to include a scoring system and additional consistency checks. Searches are performed to identify orthologues (both single-copy and multi-copy orthologues are considered) at scaffold extremities in a given assembly that form neighbouring pairs in the other compared assemblies, thereby supporting the hypothesis that these scaffolds should themselves be neighbours. ORTHOSTITCH was applied to the full anopheline input dataset to predict scaffold adjacencies (**Supplementary Online Material: Figures S3, S4 and Table S3**). The CAMSA tool (Aganezov and Alekseyev 2017) was used to compare and merge scaffold assemblies produced by the three methods by identifying adjacencies in three-way and two-way agreement (with no third method conflict) (**Supplementary Online Material: Table S4**). CAMSA was also used to build merged assemblies using only conservative three-way consensus adjacencies, and using liberal unions of all non-conflicting adjacencies. Quantifications of assembly improvements considered only scaffolds with annotated orthologous genes (because the synteny-based methods rely on orthology data) to count the numbers of scaffolds and compute scaffold N50 values before and after merging (**Fig. 2; Supplementary Online Material: Figures S5, S6**). The results of the CAMSA merging procedure were used to quantify all agreements and conflicts amongst the different sets of predicted adjacencies (**Fig. 3; Supplementary Online Material: Figures S7, S8 and Table S4**). A DOCKER container is provided that packages ADSEQ, GOS-ASM, ORTHOSTITCH, and CAMSA, as well as their dependencies, in a virtual environment that can run on a Linux server. See **Supplementary Online Material** for further details for all synteny-based predictions and their comparisons, and the DOCKER container.

### Integration of physical mapping and RNA sequencing data

Methods for chromosomal mapping of scaffolds (Sharakhova et al. 2019; Artemov et al. 2018b) are detailed for *A. albimanus* (Artemov et al. 2017), *A. atroparvus* (Artemov et al. 2015; Neafsey et al. 2015; Artemov et al. 2018a), *A. stephensi* (SDA-500) (Neafsey et al. 2015), *A. stephensi* (Indian) (Jiang et al. 2014), and *A. sinensis* (Chinese) (Wei et al. 2017). *A. funestus* mapping built on previous results (Sharakhov et al. 2002, 2004; Xia et al. 2010) with additional FISH mapping (**Supplementary Online Material: Figure S9**) to further develop the physical map by considering several different types of mapping results. *A. stephensi* mapping also extended previous efforts (Sharakhova et al. 2010) by aligning FISH probes to the AsteI2 scaffolds with BLAST, and designing and hybridising new probes targeting specific scaffolds to increase the coverage. The usable scaffold pair adjacencies are presented in **Supplementary Online Material: Table S5**, and the definitive mapped *A. funestus* scaffolds in **Supplementary Online Material: Table S6**. These adjacencies were compared with the CAMSA-generated two-way consensus assemblies, as well as the predictions from each method and the conservative and liberal consensus assemblies (**Fig. 4A; Supplementary Online Material: Table S7**). The RNAseq-based adjacency predictions used genome-mapped paired-end sequencing data for 13 of the anophelines available from VECTORBASE (Giraldo-Calderón et al. 2015) (Release VB-2017-02), including those from the *Anopheles* 16 Genomes Project (Neafsey et al. 2015) and an *A. stephensi* (Indian) male/female study (Jiang et al. 2015). AGOUTI (Zhang et al. 2016) analyses were performed to identify transcript-supported scaffold adjacencies for these 13 anophelines (**Supplementary Online Material: Table S8**). These adjacencies were compared with the CAMSA-generated two-way consensus assemblies, as well as the predictions from each method and the conservative and liberal consensus assemblies (**Fig. 4B; Supplementary Online Material: Table S9**). See **Supplementary Online Material** for further details for physical mapping and AGOUTI adjacencies and their comparisons.

### Building the new assemblies

The new assemblies were built using the different datasets available for each of the anophelines (**Supplementary Online Material: Figure S1**): synteny data only for six, *A. christyi, A. coluzzii, A. culicifacies, A. darlingi, A. maculatus*, and *A. melas*; synteny and AGOUTI data for eight, *A. arabiensis, A. dirus, A. epiroticus, A. farauti, A. merus, A. minimus, A. quadriannulatus*, and *A. sinsensis* (SINENSIS); synteny and physical mapping data for *A. sinensis* (Chinese); synteny, AGOUTI, and physical mapping data for four, *A. albimanus, A. atroparvus, A. stephensi* (SDA-500), *A. stephensi* (Indian); and synteny, AGOUTI, physical mapping data, and the new PacBio-based assembly for *A. funestus*. The new *A. arabiensis* assembly additionally incorporated scaffold orders determined by alignments to the *A. gambiae* (PEST) X chromosome from (Fontaine et al. 2015) and to autosomes provided by Xiaofang Jiang and Brantley Hall. The new *A. funestus* assembly generated as part of this study was based on approximately 70X of PacBio sequencing data polished with QUIVER (from PacBio’s SMRT Analysis software suite). This was combined with the reference assembly (AfunF1) using METASSEMBLER (Wences and Schatz 2015) to generate a merged assembly, and this merged assembly was then scaffolded with SSPACE (Boetzer et al. 2011) using the original Illumina sequencing data, and designated the *A. funestus* AfunF2-IP assembly. The AfunF2-IP assembly improves on the reference AfunF1 assembly at contig level but not at scaffold level (**Supplementary Online Material: Figure S10 and Table S10**). Where AfunF2-IP scaffolds span the ends of AfunF1 scaffolds they provide support for AfunF1 scaffold adjacencies. Thus, whole genome alignments of the two assemblies were performed using LASTZ (Harris 2007) and used to identify corresponding genomic regions that enabled the alignment-based ordering and orientation of AfunF1 scaffolds, which were then compared with the synteny-based, physical mapping based, and AGOUTI-based, adjacencies (**Fig. 5, Supplementary Online Material: Figure S11 and Table S11**). Using the AfunF1 assembly as the basis, and incorporating evidence from the AfunF2-IP assembly through scaffold correspondences established from the whole genome alignments, the physical mapping data and the synteny-based and AGOUTI-based adjacency predictions were integrated to build the new reference assembly for *A. funestus*. The comprehensive update to the photomap employed BLAST searches to identify positions of the physically mapped DNA markers within the AfunF1 and AfunF2-IP assemblies, and whole genome pairwise alignments to reconcile these two assemblies with the new photomap. Whole genome alignments of versions 1 and 2 assemblies for *A. farauti*, and *A. merus* were used to delineate corresponding scaffolds and identify supported, unsupported, and conflicting adjacencies (**Supplementary Online Material: Figure S12 and Table S12**). Reconciling all adjacencies produced the resolved sets of scaffold adjacencies and superscaffolds that were used to build all the new assemblies and the definitive chromosome anchoring data for seven assemblies. These updated assemblies, their correspondingly updated gene annotations, the orthology data used as input for the gene synteny-based approaches, and the definitive anchoring data, were employed to assign non-anchored scaffolds to chromosome arms (**Supplementary Online Material: Table S13**). See **Supplementary Online Material** for further details on the workflow to integrate different adjacency predictions and build the new assemblies, the PacBio assembly generation, the genome alignment based comparisons of the AfunF1 and AfunF2-IP assemblies, the lift-over of gene annotations to the new assemblies, and the assignment of non-anchored scaffolds and superscaffolds to chromosome arms.

## Abbreviations

AD: ADSEQ
AGO: AGOUTI-based
AGOUTI: annotated genome optimization using transcriptome information tool
ALN: alignment-based
CAMSA: comparative analysis and merging of scaffold assemblies tool
DP: dynamic programming
FISH: fluorescence in situ hybridization
GA: GOS-ASM
GOS-ASM: Gene order scaffold assembler
Kbp: kilobasepairs
Mbp: megabasepairs
OS: ORTHOSTITCH
PacBio: Pacific Biosciences
PB: PacBio-based
PHY: physical-mapping-based
RNAseq: RNA sequencing
QTL: quantitative trait loci
SYN: synteny-based.

## Acknowledgements

The authors acknowledge Marcia Kern for technical assistance with the physical mapping, Vasily Sitnik for assistance with data submission to VectorBase, and the Montpellier Bioinformatics Biodiversity platform service for ADSEQ data computation and analyses. Physical mapping and PacBio sequencing of *A. funestus* were supported by the United States (US) National Institutes of Health (NIH) National Institute of Allergy and Infectious Diseases (NIAID) grant R21 AI112734 to NJB, with SJE and IVS as co-investigators. IVS was supported by the US NIH NIAID grants R21AI099528 and R21AI135298 and by the US Department of Agriculture National Institute of Food and Agriculture Hatch project 223822. SA and MAA were supported by the US National Science Foundation (NSF) grant IIS-1462107. SA was supported by the US NSF grants CCF-1053753 and DBI-1350041 and by US NIH grants U24CA211000 and R01-HG006677. YA, SB and ET were supported by the French Agence Nationale pour la Recherche Ancestrome project ANR-10-BINF-01-01. SK and AMP were supported by the Intramural Research Program of the NIH National Human Genome Research Institute 1ZIAHG200398. CC was supported by a Mitacs Globalink grant, the Natural Sciences and Engineering Research Council of Canada Discovery Grant RGPIN-249834, and a resource allocation from Compute Canada. MWH and SVZ were supported by US NSF grant DEB-1249633. RMW, LR, MJMFR, and RF were supported by Novartis Foundation for medical-biological research grant #18B116 and Swiss National Science Foundation grant PP00P3_170664.

## Author contributions

RMW and IVS conceived the study. SA and MAA developed and implemented GOS-ASM and CAMSA. YA, SB, ET and CC developed and implemented ADSEQ. RMW developed and implemented ORTHOSTITCH. SJE contributed to synteny-based analyses. JL, PG, MK, AP, MVS, MFU and IVS carried out physical mapping experiments. RMW, MWH and SVZ performed AGOUTI analyses. RMW, SA, LR and MJMFR compared synteny-based with physical mapping and AGOUTI adjacencies. PacBio *funestus* sequencing and data production: PIH, SJE and NJB; assembly: SK and AMP; and assembly comparisons: RMW, JL, MVS and IVS. Reconciliation to produce updated assemblies: RMW, JL, LR, MJMFR, RF, DL, GM and IVS. Chromosome arm assignment: RF. Docker container: MJMFR. The manuscript was written by RMW with input from all authors. Author contributions are further detailed in **Supplementary Online Material**. All authors read and approved the manuscript.

