## Supplementary Online Material for "Evolutionary superscaffolding and chromosome anchoring to improve *Anopheles* genome assemblies"

#### Author emails & ORCIDs

| Name | Email | ORCID |
|---|---|---|
| Robert M. Waterhouse | | 0000-0003-4199-9052 |
| Sergey Aganezov | | 0000-0003-2458-8323 |
| Yoann Anselmetti | | 0000-0002-6689-1163 |
| Jiyoung Lee | | 0000-0003-1702-874X |
| Livio Ruzzante | | 0000-0002-8693-8678 |
| Maarten J.M.F. Reijnders | | 0000-0002-5657-4762 |
| Romain Feron | | 0000-0001-5893-6184 |
| Sèverine Bérard | | 0000-0002-3029-0964 |
| Phillip George | | NA |
| Matthew W. Hahn | | 0000-0002-5731-8808 |
| Paul I. Howell | | 0000-0002-3834-4621 |
| Maryam Kamali | | 0000-0001-9955-0683 |
| Sergey Koren | | 0000-0002-1472-8962 |
| Daniel Lawson | | 0000-0001-7765-983X |
| Gareth Maslen | | 0000-0001-7318-3678 |
| Ashley Peery | | NA |
| Adam M. Phillippy | | 0000-0003-2983-8934 |
| Maria V. Sharakhova | | 0000-0002-5790-3548 |
| Eric Tannier | | 0000-0002-3681-7536 |
| Maria F. Unger | | NA |
| Simo V. Zhang | | 0000-0003-2154-2549 |
| Max A. Alekseyev | | 0000-0002-5140-8095 |
| Nora J. Besansky | | 0000-0003-0646-0721 |
| Cedric Chauve | | 0000-0001-9837-1878 |
| Scott J. Emrich | | 0000-0002-4804-7436 |
| Igor V. Sharakhov | | 0000-0003-0752-3747 |

### [1] Data analysis and adjacency reconciliation workflow overview

*Robert M. Waterhouse*

Details of all steps are presented in the following sections, here we provide an overview of the production of different sets of scaffold adjacencies for each of the anophelines and the different workflows that were followed to reconcile all the data to build the new assemblies (**Figure S1**). The simplest workflow (**A**, six assemblies) was used for *A. christyi*, *A. coluzzii*, *A. culicifacies*, *A. darlingi*, *A. maculatus*, and *A. melas*, for which only consensus synteny predictions were produced. Workflow **B** (eight assemblies) reconciled the synteny-based two-way consensus sets with the adjacency predictions from RNA sequencing (RNAseq) data using the AGOUTI tool (Zhang et al. 2016) to build new assemblies for *A. arabiensis*, *A. dirus*, *A. epiroticus*, *A. farauti*, *A. merus*, *A. minimus*, *A. quadriannulatus*, and *A. sinsensis* (SINENSIS). Workflow **C** (four assemblies) additionally incorporated reconciliations with the available physical mapping data for *A. albimanus*, *A. atroparvus*, *A. stephensi* (SDA-500), and *A. stephensi* (Indian). Workflow **D** was applied to *A. funestus* to also incorporate reconciliations with the adjacencies produced from comparing the reference assembly (AfunF1) with the new Pacific Biosciences (PacBio) assembly (AfunF2-IP). And finally, workflow **E** was adopted for *A. sinensis* (Chinese) that employed just the synteny-based two-way consensus set and the physical mapping data. Finally, chromosome mapping data from *A. arabiensis* were combined with the workflow B results to produce the new chromosome-anchored assembly.

We employed gene orthology data delineated using ORTHODB (Zdobnov et al. 2017), but alternative methodologies may be used to define orthologous relations amongst the annotated gene sets of the species to be analysed. With gene orthology data and genomic location data from VECTORBASE (Giraldo-Calderón et al. 2015) prepared, we performed adjacency predictions with GOS-ASM (Aganezov and Alekseyev 2016) and ORTHOSTITCH (this study) directly, while ADSEQ (Anselmetti et al. 2015, 2018) first required building sequence alignments and reconciled trees before scaffold neighbours were predicted (see the following sections for details). We then employed the CAMSA tool (Aganezov and Alekseyev 2017) for comparative analyses of the results from our different scaffold adjacency predictions to automatically build the most confident merged-scaffold assembly, and we used CAMSA's interactive visualisation framework to inspect conflicts in the assembly graph. For the species with no validation datasets we employed a simple two-way consensus approach with no third-method conflicts to define the final adjacencies. For the other species, all conflicts identified between the two-way consensus adjacencies and the alternative sources of adjacency information were manually resolved, the most complex being for *A. funestus* with the reconciliation of synteny, RNAseq (AGOUTI), PacBio-AfunF2-IP-alignment, and physical mapping data, and the construction of a new cytogenetic photomap.

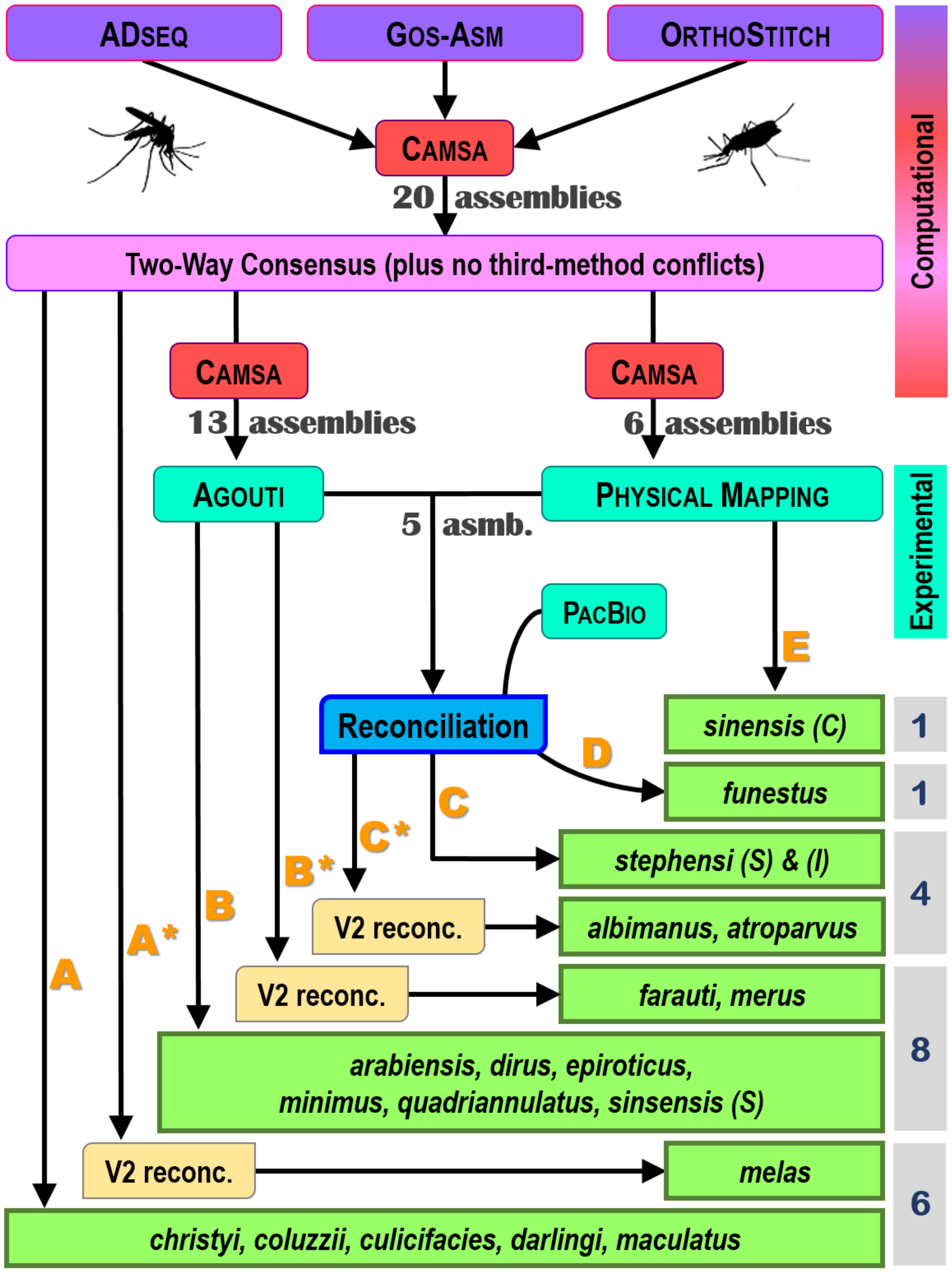

**Figure S1. Workflows applied to upgrade the 20 anopheline assemblies**

A: two-way synteny only. B: two-way synteny and AGOUTI. C: two-way synteny, AGOUTI, and physical mapping data. D: two-way synteny, AGOUTI, physical mapping data, and PacBio sequencing data. E: two-way synteny and physical mapping data. Asterisks (*) indicate additional reconciliation with version 2 assemblies (V2 reconc.) for a subset of species.

### [2] Superscaffolding and chromosome arm assignments

*Robert M. Waterhouse, Livio Ruzzante, Maarten J.M.F. Reijnders, Romain Feron*

The integrated approach to reconciling the different sources of scaffold adjacencies with available experimental data outlined above and detailed in the sections below improved assembly contiguity through building well-supported superscaffolds (**Table 1, main text**), as well as enhancing the anchoring of ordered and oriented scaffolds to chromosome arms (**Table 2, main text**), and enabling the assignment of non-anchored scaffolds and superscaffolds to chromosome arms (**Table S1**). The resulting superscaffolds had total spans ranging from more than 200 Mbps for *A. arabiensis* to fewer than 20 Mbps for *A. maculatus*, reflecting the contiguity of the input assemblies and the availability of complementary datasets to support superscaffolding (**Figure S2**). For ten assemblies the total span of superscaffolds comprised more than half the total assembly size, and they made up more than a quarter of a further seven assemblies (**Figure S2**).

The enhanced chromosome anchoring for a subset of the anophelines (**Table 2, main text**) and the chromosomal-level assembly for *A. gambiae* PEST together allowed for the assignment of non-anchored scaffolds and superscaffolds to chromosome arms. Enumerating shared orthologues between non-anchored scaffolds and the eight species with chromosome-anchored scaffolds (see section [12] below for details) enabled assignments with support from multiple species (**Table S1**).

**Table S1. Assignment of scaffolds and superscaffolds to chromosome arms**
Scaffold counts and proportions of the 20 updated assemblies with chromosome arm assignments.

| Species | Assembly Version | Assigned Scaffolds or Superscaffolds [Also Anchored] | % Assembly Assigned |
|---|---|---|---|
| *Anopheles albimanus* | AalbS3 | 7 [7] | 97 |
| *Anopheles arabiensis* | AaraD2 | 5 [5] | 88 |
| *Anopheles atroparvus* | AatrE4 | 10 [10] | 88 |
| *Anopheles christyi* | AchrA2 | 154 | 8 |
| *Anopheles coluzzii* | AcolM2 | 65 | 85 |
| *Anopheles culicifacies* | AculA2 | 286 | 19 |
| *Anopheles darlingi* | AdarC4 | 247 | 48 |
| *Anopheles dirus* | AdirW2 | 36 | 91 |
| *Anopheles epiroticus* | AepiE2 | 215 | 78 |
| *Anopheles farauti* | AfarF3 | 29 | 97 |
| *Anopheles funestus* | AfunF2 | 136 [81] | 90 |
| *Anopheles maculatus* | AmacM2 | 2 | 0 |
| *Anopheles melas* | AmelC3 | 106 | 4 |
| *Anopheles merus* | AmerM3 | 119 | 75 |
| *Anopheles minimus* | AminM2 | 22 | 96 |
| *Anopheles quadriannulatus* | AquaS2 | 105 | 80 |
| *Anopheles sinensis* | AsinS3 | 222 | 56 |
| *Anopheles sinensis (Chinese)* | AsinC3 | 165 [29] | 70 |
| *Anopheles stephensi* | AsteS2 | 150 [71] | 89 |
| *Anopheles stephensi (Indian)* | AsteI3 | 72 [60] | 83 |

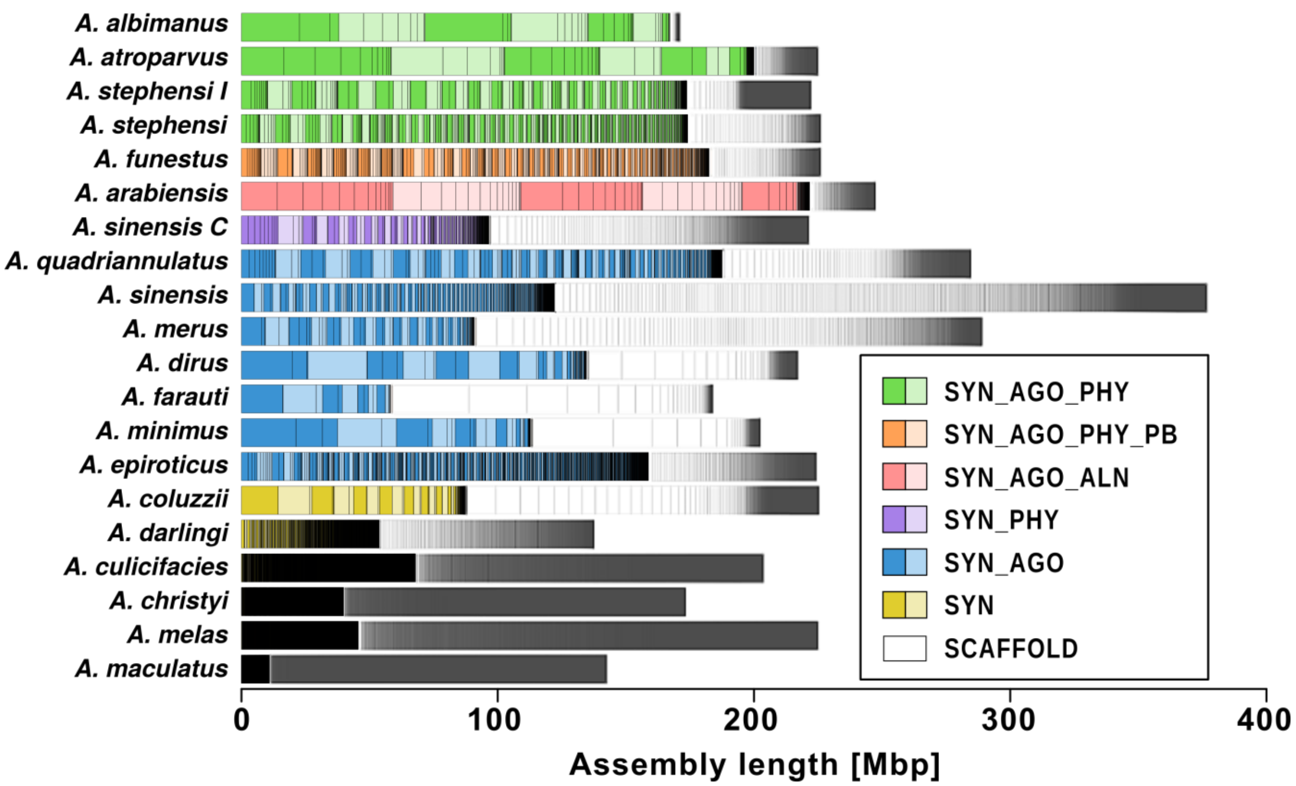

**Figure S2. Superscaffolding genomic spans of 20 anopheline genome assemblies**

Superscaffolds are shown as stacked bars of alternating dark and light colours with lines within each superscaffold indicating the sizes (y-axis, basepairs) of their constituent scaffolds, and with superscaffolds and scaffolds ordered from the largest (left) to the smallest (right). The stacked bars continue with scaffolds that are not part of superscaffolds in grey, again ordered from the largest to the smallest. The assemblies are grouped and coloured according to the types of data and approaches used to perform the superscaffolding as presented in the legend and in main text Table 1. Approaches: synteny-based (SYN), and/or RNAseq AGOUTI-based (AGO), and/or alignment-based (ALN), and/or physical mapping-based (PHY), and/or PacBio sequencing-based (PB). Results for two strains are shown for *Anopheles sinensis*, SINENSIS and Chinese (C), and *Anopheles stephensi*, SDA-500 and Indian (I).

### [3] Sources of input data for predicting adjacencies

*Robert M. Waterhouse*

The orthology data used as inputs for each of the three synteny-based methods were retrieved from ORTHODB V9.1 (www.orthodb.org) (Zdobnov et al. 2017). These orthologous groups included all the anophelines apart from *A. sinensis* SINENSIS strain and *A. stephensi* Indian strain, so proteins from the gene sets of these two anophelines were mapped to the ORTHODB anopheline orthologous groups using the complete species mapping approach of ORTHODB. The protein sequences used by ORTHODB, and the gene annotations required for the adjacency predictions, were retrieved from VECTORBASE (Giraldo-Calderón et al. 2015). The versions of the genome assemblies and their annotated gene sets are detailed in **Table S2**, along with counts of scaffolds, genes, and orthologues.

**Table S2. Assembly and orthology input data**

Genome assembly versions, scaffold counts, gene set versions, gene counts, and ORTHODB orthologous groups (from ORTHODB V9.1) across 21 anophelines used as input data for the synteny-based scaffold adjacency predictions.

| Species | Assembly | Scaffolds | Gene set | Total genes | Genes in orthogroups | Scaffolds with genes | Scaffolds with orthologs |
|---|---|---|---|---|---|---|---|
| *Anopheles albimanus* | AalbS1 | 204 | AalbS1.3 | 12085 | 10637 | 57 | 52 |
| *Anopheles arabiensis* | AaraD1 | 1214 | AaraD1.3 | 13333 | 12132 | 340 | 289 |
| *Anopheles atroparvus* | AatrE1 | 1371 | AatrE1.3 | 13789 | 12249 | 476 | 384 |
| *Anopheles christyi* | AchrA1 | 30369 | AchrA1.2 | 10738 | 10103 | 5173 | 5064 |
| *Anopheles coluzzii* | AcolM1 | 10521 | AcolM1.2 | 14710 | 12998 | 1124 | 816 |
| *Anopheles culicifacies* | AculA1 | 16162 | AculA1.2 | 14335 | 13002 | 5715 | 5200 |
| *Anopheles darlingi* | AdarC3 | 2221 | AdarC3.2 | 10457 | 9871 | 2161 | 2055 |
| *Anopheles dirus* | AdirW1 | 1266 | AdirW1.3 | 12840 | 11488 | 302 | 250 |
| *Anopheles epiroticus* | AepiE1 | 2673 | AepiE1.3 | 12181 | 11549 | 1053 | 1004 |
| *Anopheles farauti* | AfarF1 | 550 | AfarF1.2 | 13217 | 12146 | 376 | 355 |
| *Anopheles funestus* | AfunF1 | 1392 | AfunF1.2 | 13344 | 11616 | 619 | 575 |
| *Anopheles gambiae* | AgamP4 | 8 | AgamP4.2 | 12843 | 12240 | 7 | 7 |
| *Anopheles maculatus* | AmacM1 | 47797 | AmacM1.2 | 14835 | 11777 | 12776 | 10297 |
| *Anopheles melas* | AmelC1 | 20281 | AmelC1.2 | 16149 | 14718 | 8855 | 8223 |
| *Anopheles merus* | AmerM1 | 2753 | AmerM1.2 | 13886 | 13076 | 1078 | 1036 |
| *Anopheles minimus* | AminM1 | 678 | AminM1.3 | 12663 | 11436 | 142 | 121 |
| *Anopheles quadriannulatus* | AquaS1 | 2823 | AquaS1.3 | 13484 | 12055 | 647 | 576 |
| *Anopheles sinensis* | AsinS2 | 10448 | AsinS2.1 | 12869 | 11037 | 1825 | 1486 |
| *Anopheles sinensis (Chinese)* | AsinC2 | 9592 | AsinC2.1 | 19352 | 11594 | 702 | 573 |
| *Anopheles stephensi* | AsteS1 | 1110 | AsteS1.3 | 13227 | 11645 | 502 | 479 |
| *Anopheles stephensi (Indian)* | AsteI2 | 23371 | AsteI2.2 | 11789 | 11100 | 906 | 660 |

### [4] ADSEQ: scaffolding genomes using gene trees, synteny and sequencing data

*Yoann Anselmetti, Sèverine Bérard, Eric Tannier, Cedric Chauve*

**Gene trees.** Gene trees contain the information about how genes, and the traits they are related to, evolve along the history of the species. They give access to information about adaptations by substitutions, gene gains and losses, duplications, transfers. Gene trees can also be used to detect co-evolutionary elements in genomes. Moreover, gene trees are useful to reconstruct ancestral genomes and provide better assemblies for extant species, as shown in (Duchemin et al. 2017; Anselmetti et al. 2015, 2018). However, the quality of the results highly depends on the quality of the gene trees. For the *Anopheles* genomes, genes were clustered into ORTHODB orthologous groups (**Table S2**), and multiple alignments were computed for each group using MUSCLE v.3.8.425 (Edgar 2004). These were then used as input for RAXML (Stamatakis 2014) phylogenetic tree estimations, in a large-scale automatic effort. A substantial number of branches are probably incorrect, since multiple sequence alignments often do not contain enough signal to fully resolve the gene tree. We applied the gene tree correction program TREERECS (https://gitlab.inria.fr/Phylophile/Treerecs) to correct these gene trees and our preliminary analysis shows that the corrected trees are of better quality (in terms of ancestral gene content) than the original ones.

**Scaffolding extant and ancestral genomes.** In a second step we used these improved reconciled gene trees to reconstruct jointly ancestral and extant gene adjacencies. The general approach is described in our recent papers (Anselmetti et al. 2015, 2018): we consider pairs of gene families for which extant adjacencies (synteny) is observed and compute, from the reconciled gene trees, a duplication-aware parsimonious evolutionary scenario in terms of adjacency gain/breaks that can also create extant adjacencies between genes at the extremities of contigs/scaffolds. The method has been modified to include sequencing data for the inference of potential extant scaffolding adjacencies, thus it is based on a combination of evolutionary signal and sequence data. We used all sequencing data available for the 21 anophelines to associate a prior score to potential extant scaffolding adjacencies with the scaffolder BESST (Sahlin et al. 2014). The new method, using both sentence evolution and sequencing data is called ADSEQ; it includes a probabilistic version of the algorithm that allows sampling of optimal solutions uniformly and to associate to potential scaffolding (both extant and ancestral) a posterior score defined as the frequency of observing the adjacencies in this sample. Finally, if adjacency conflicts are observed (e.g. the same contig extremity is deemed to be adjacent to more than one other contig extremity), we use a Maximum Weight Matching algorithm to resolve these conflicts, using the posterior score of the adjacencies as edge weights. Resulting counts of predicted scaffold adjacencies for each of the anopheline assemblies are presented in **Table S3**.

**Data and code availability**. Input data and results obtained with ADSEQ are available from the GitHub repository https://github.com/YoannAnselmetti/DeCoSTAR_pipeline in the directory named "21Anopheles_dataset". This contains a pipeline written in snakemake, a python workflow management system (Köster and Rahmann 2012), allowing users to generate input data required for ADSEQ and execute it from standard genomic format files.

### [5] GOS-ASM: multi-genome rearrangement-based gene order scaffolder

*Sergey Aganezov, Max A. Alekseyev*

GOS-ASM (Aganezov and Alekseyev 2016) (https://github.com/aganezov/gos-asm, Gene order scaffold assembler) starts with an assumption that the constructed scaffolds that make up the input assemblies are accurate and long enough to allow for the identification of orthologous genes. The scaffolds can then be represented as ordered sequences of oriented genes and the scaffold assembly problem can be posed as the reconstruction of the global gene order (along genome chromosomes) from the gene sub-orders defined by the scaffolds. Such gene sub-orders are viewed as the result of both evolutionary events and artificial "technological" fragmentation in the genome. Evolutionary events that change gene orders are genome rearrangements, most common of which are reversals, fusions, fissions, and translocations. Technological fragmentation is modelled by artificial "fissions" that break genomic chromosomes into scaffolds. Scaffold assembly can therefore be reduced to the search for "fusions" that revert technological "fissions" and glue scaffolds back into chromosomes. This observation inspired us to employ the genome rearrangement analysis techniques for scaffolding purposes. Rearrangement analysis of multiple genomes relies on the concept of the breakpoint graph and utilizes the topology of the organisms' phylogenetic tree. While traditionally the breakpoint graph is constructed for complete genomes, it can also be constructed for fragmented genomes, where we treat scaffolds as "chromosomes". We demonstrate that the breakpoint graph of multiple genomes possesses an important property that its connected components are robust with respect to the technological genome fragmentation. In other words, connected components of the breakpoint graph mostly retain information about the complete genomes, even when the breakpoint graph is constructed on their scaffolds. We thus use the topology of the species phylogenetic tree and the structure of the connected components in the corresponding breakpoint graph to reconstruct the "reverse evolution" of the input genomes along the branches of the phylogenetic tree, distinguishing between signatures of evolutionary and technological fissions. Identified technological fissions are then used as guidance for the gluing of input scaffolds back into complete chromosomes. Resulting counts of predicted scaffold adjacencies from applying GOS-ASM to the full set of anopheline assemblies are presented in **Table S3**.

### [6] ORTHOSTITCH: scaffold adjacencies from conserved orthologous neighbours

*Robert M. Waterhouse*

Using gene orthology data from cross-species comparisons, ORTHOSTITCH identifies genes located at scaffold extremities and evaluates the evidence from the locations of orthologous genes from other species to predict likely scaffold adjacencies. The analysis proceeds in a stepwise manner, first identifying the most likely neighbour for each scaffold end and then requiring best neighbours to be reciprocal in order to identify putative adjacencies. The evaluations are not limited to single-copy orthologues as analyses of all paralogues are performed such that all possible neighbour relationships are examined. Putative neighbours at scaffold extremities are scored by how many of the species with orthologues show the same neighbour relationship (**Figure S3**), requiring at least two species to do so. ORTHOSTITCH was developed as part of the synteny-focused analyses of the comparative analysis of the *Manduca sexta* genome (Kanost et al. 2016), it is described in detail below and the code is available from the GitLab project page: https://gitlab.com/rmwaterhouse/OrthoStitch

| Species | Scaffold | GeneID[GroupID]#neighbours | | GeneID[GroupID]#neighbours | Scaffold |
|---|---|---|---|---|---|
| AFUNE | KB668690[+] | AFUN010217[EOG09170540]1 | ---x>+-x--> | AFUN000326[EOG091701PP]1 | KB668920[+] |
| AALBI | KB672397 | AALB002374[EOG09170540]2 | --x--o--x-- | AALB002371[EOG091701PP]2 | KB672397 |
| AARAB | KB704451 | AARA004744[EOG09170540]2 | ---x---x--- | AARA004745[EOG091701PP]2 | KB704451 |
| AATRO | KI421897 | AATE015926[EOG09170540]2 | ---x---x--- | AATE019573[EOG091701PP]2 | KI421897 |
| ACHRI | KB698096 | ACHR007688[EOG09170540]0 | -x-- .... | noortho[NOOG] | na |
| ACOLU | scf_1925491386 | ACOM037460[EOG09170540]2 | ---x---x--- | ACOM037465[EOG091701PP]2 | scf_1925491386 |
| ACULI | KI423732 | ACUA013460[EOG09170540]0 | -x-- --x- | ACUA014152[EOG091701PP]1 | KI424031 |
| ADARL | na | noortho[NOOG] | .... --x- | ADAC004066[EOG091701PP]2 | scaffold_20 |
| ADIRU | KB672868 | ADIR002642[EOG09170540]2 | ---x---x--- | ADIR002641[EOG091701PP]2 | KB672868 |
| AEPIR | KB672164 | AEPI009033[EOG09170540]2 | -x-- --x- | AEPI005575[EOG091701PP]2 | KB671247 |
| AFARA | KI421545 | AFAF012506[EOG09170540]2 | ---x---x--- | AFAF012254[EOG091701PP]2 | KI421545 |
| AGAMB | 2R | AGAP002925[EOG09170540]2 | ---x---x--- | AGAP002926[EOG091701PP]2 | 2R |
| AMACU | AXCL01014811 | AMAM000110[EOG09170540]0 | -x-- --x- | AMAM010795[EOG091701PP]0 | AXCL01051139 |
| AMELA | KI429153 | AMEC010445[EOG09170540]1 | ---x---x--- | AMEC021444[EOG091701PP]2 | KI429153 |
| AMERU | KI438982 | AMEM010727[EOG09170540]2 | ---x---x--- | AMEM012929[EOG091701PP]2 | KI438982 |
| AMINI | KB663832 | AMIN000747[EOG09170540]2 | ---x---x--- | AMIN000748[EOG091701PP]2 | KB663832 |
| AQUAD | KB666065 | AQUA009280[EOG09170540]2 | ---x---x--- | AQUA009279[EOG091701PP]2 | KB666065 |
| ASINC | AS2_scf7180000695544 | ASIC004586[EOG09170540]2 | --x--o--x-- | ASIC004611[EOG091701PP]2 | AS2_scf7180000695544 |
| ASINS | KI916183 | ASIS000523[EOG09170540]2 | --x--o--x-- | ASIS000668[EOG091701PP]2 | KI916183 |
| ASTEI | scaffold_00001 | ASTEI00055[EOG09170540]2 | ---x---x--- | ASTEI00054[EOG091701PP]2 | scaffold_00001 |
| ASTES | KB665265 | ASTE007167[EOG09170540]2 | ---x---x--- | ASTE007166[EOG091701PP]2 | KB665265 |

**Figure S3. Example of ORTHOSTITCH adjacency evidence**

This putative adjacency is identified in *A. funestus* (AFUNE, blue) with both scaffolds in the forward orientation, where orthologous genes from 12 other anophelines support the neighbour relationship (green). In three other anophelines one or more intervening genes disrupt the neighbour relationship of these pairs of orthologues (orange). In the remaining five anophelines there are no orthologues or the orthologues have no neighbouring genes and thus they offer neither support nor evidence against the putative neighbour relationship (yellow), or there are orthologues with neighbours but they do not support the putative adjacency (purple). So this adjacency is supported by evidence from 12 species out of a possible 16 for scaffold KB668690 and out of a possible 18 for scaffold KB668920, giving a synteny score of 0.71 and a universality score of 0.85 with a final adjacency score of 0.60.

ORTHOSTITCH requires as input an anchor groups file and an anchor locations file. The anchor groups file may be generated from any orthology delineation procedure, and consists of just three columns of data: the orthologous group identifier, the gene identifier, and the species identifier. The anchor locations file may be generated from general feature format (GFF) or general transfer format (GTF) files that indicate the genomic locations of annotated features (genes) for each assembly. Like GFF or GTF files, the anchor locations file consists of nine columns, with only the coding sequence (CDS) lines selected from GFF or GTF files, and with the 'source' column ($2^{nd}$ column) containing the species identifier, and with the 'attribute' column ($9^{th}$ column) containing only the gene identifier. The gene and species identifiers used in both the groups file and the locations file must match exactly, and gene identifiers must be unique across the complete dataset of all species. The anchor locations file may contain the locations of genes that are not present in the anchor groups file, i.e. some genes with known locations may not have been assigned to any orthologous group, however, the anchor groups file may not contain any genes that are not present in the anchor locations files, i.e. all genes in orthologous groups must have known locations.

ORTHOSTITCH options allow for the genomic location of each anchor gene to be set as the start, middle, or end of the input coding sequence genomic coordinates, and the analyses can be run using only genes with orthologues or with all genes in the locations file. All predicted adjacencies are further classified into confident, and superconfident subsets. Confident adjacencies require more than a third of comparison species to have orthologues and more than a third of those that do have orthologues to support the predicted scaffold adjacency. Superconfident adjacencies additionally require the same of their upstream or downstream neighbours. The adjacency score for each pair of putatively neighbouring scaffolds is computed as the product of a synteny score ***(S)*** and a universality score ***(U)***, based on the numbers of species with orthologues that support the adjacency where *Sup* = the number of supporting species, *Pos* = the number of possible species, and *Tot* = the total number of species thus:

$$S = \frac{\left(\frac{Sup1}{Pos1} + \frac{Sup2}{Pos2}\right)}{2} \qquad U = \frac{\left(\frac{Pos1 + Pos2}{2}\right)}{Tot - 1}$$

Orthology data from ORTHODB v9 (Zdobnov et al. 2017), were used to produce the input anchor groups file and the anchor locations were produced from GFF files from VECTORBASE (Giraldo-Calderón et al. 2015) (see **Table S2**). The ORTHOSTITCH (v1.6) analysis was run using data from all 21 available anophelines with the options of anchor locations set to 'middle' and using all annotated genes, and the resulting adjacency counts are presented in **Table S3**.

The performance of ORTHOSTITCH in terms of the ability to recover true adjacencies was assessed using the same input dataset from the 21 anophelines with the introduction of artificial scaffold/chromosome breaks. Four different types of randomly positioned scaffold/chromosome-splitting breaks were introduced and analysed separately, (i) between any neighbouring pair of orthologues; (ii) between neighbouring orthologue pairs both from orthologous groups containing at least a third of the 21 species; (iii) between neighbouring orthologue pairs both from orthologous groups with more than half of the 21 species, a gene-to-species ratio of no more than 1.5 (i.e. limiting the numbers of duplicated copies), and restricted to scaffolds/chromosomes with at least 25 orthologues in total (i.e. avoiding splitting shorter scaffolds); and (iv) the same as (iii) but also requiring the neighbouring pair to have been part of the supporting sets that defined the superconfident adjacencies in **Table S3** (i.e. known to provide synteny support). 100 random scaffold/chromosome breaks were introduced and then analysed to predict putative adjacencies and assess how many of the artificially introduced breaks were recovered as predicted adjacencies, repeated 100 times for each of the four different types of neighbouring orthologues. ORTHOSTITCH options were selected as for the complete analysis above, with anchor locations set to 'middle' and using all annotated genes. Median recoveries were 76%, 81%, 88%, and 97% for the four types, respectively (**Figure S4**). Thus for similar datasets ORTHOSTITCH is expected to be able to recover about three quarters of true adjacencies, when the genes at the scaffold extremities have orthologues in more than a third or more than half the species then recovery levels are expected to increase, and when these orthologues provide synteny support then the adjacencies are almost always recovered.

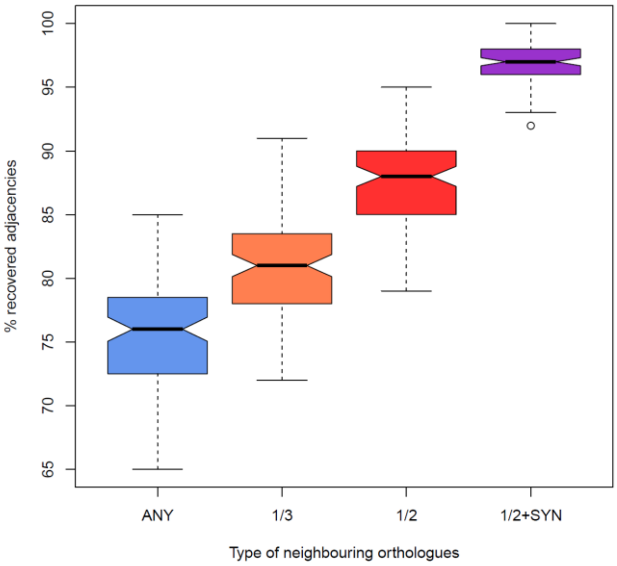

**Figure S4. Performance of ORTHOSTITCH adjacency recovery**

For each of four different types of neighbouring orthologues (ANY, 1/3, 1/2, 1/2+SYN, see text for details), a total of 100 random scaffold/chromosome breaks were introduced into the gene locations data. These were then analysed to predict putative adjacencies and assess how many introduced breaks were recovered as predicted adjacencies. This was repeated 100 times for each of the four different types of neighbouring orthologues.

**Table S3. Synteny-based adjacency predictions**

Counts of predicted adjacencies from running three methods across 21 anophelines.

| Species | ADSEQ | GOS-ASM | ORTHOSTITCH | | |
|---|---|---|---|---|---|
| | | | All | Confident | Superconfident |
| *Anopheles albimanus* | 1 | 2 | 4 | 4 | 3 |
| *Anopheles arabiensis* | 48 | 11 | 33 | 27 | 7 |
| *Anopheles atroparvus* | 42 | 6 | 29 | 24 | 2 |
| *Anopheles christyi* | 3220 | 1176 | 2031 | 1820 | 371 |
| *Anopheles coluzzii* | 199 | 71 | 134 | 114 | 14 |
| *Anopheles culicifacies* | 3594 | 2055 | 1821 | 1620 | 373 |
| *Anopheles darlingi* | 678 | 290 | 684 | 551 | 109 |
| *Anopheles dirus* | 63 | 19 | 42 | 36 | 10 |
| *Anopheles epiroticus* | 608 | 143 | 471 | 369 | 190 |
| *Anopheles farauti* | 177 | 75 | 119 | 92 | 36 |
| *Anopheles funestus* | 331 | 100 | 211 | 167 | 78 |
| *Anopheles gambiae* | 0 | 0 | 0 | 0 | 0 |
| *Anopheles maculatus* | 6366 | 1859 | 2411 | 2284 | 74 |
| *Anopheles melas* | 5081 | 3080 | 2181 | 2001 | 350 |
| *Anopheles merus* | 590 | 220 | 422 | 326 | 114 |
| *Anopheles minimus* | 19 | 7 | 14 | 8 | 4 |
| *Anopheles quadriannulatus* | 238 | 122 | 163 | 130 | 38 |
| *Anopheles sinensis* | 336 | 196 | 218 | 190 | 10 |
| *Anopheles sinensis (Chinese)* | 158 | 166 | 78 | 60 | 14 |
| *Anopheles stephensi* | 277 | 106 | 182 | 134 | 64 |
| *Anopheles stephensi (Indian)* | 201 | 69 | 155 | 124 | 43 |

### [7] CAMSA: comparative analysis and merging of scaffold assemblies

*Robert M. Waterhouse, Sergey Aganezov, Livio Ruzzante, Maarten J.M.F. Reijnders, Max A. Alekseyev*

The CAMSA tool automates the process of comparing and merging scaffold assemblies produced by alternative methods as well as providing interactive visualisations that enable detailed manual inspections of the scaffold adjacency agreements and conflicts identified during the merging process (Aganezov and Alekseyev 2017). CAMSA allows working with both oriented and (partially) un-oriented scaffold assemblies under the same unifying framework, thus greatly simplifying the downstream analysis process when working with data produced by both computational and wet-lab based methods. CAMSA (version 1.1.0b14, https://github.com/compbiol/CAMSA) was applied to the predicted adjacencies from each of the three synteny-based methods to produce three consensus sets for each of the 20 anopheline assemblies: conservative three-way consensus adjacency sets, two-way consensus adjacency sets with no third-method conflicts, and liberal union sets of all non-conflicting adjacencies. Pre-filtering of the predicted adjacencies first removed any pairs of scaffolds where one or both remained un-oriented (i.e., semi-un-oriented assembly pairs were removed). Thus common adjacencies must agree both at the level of being predicted neighbours and their relative orientations. Conflicting adjacencies occur when one or both scaffolds in a pair predicted by one method are predicted to be paired with a different scaffold (or the same scaffold but the opposite orientation) by another method. The remaining unique and non-conflicting adjacencies from each method formed part of the liberal union sets.

Adjacencies in three-way and two-way agreement in the resulting CAMSA-produced consensus sets (**Table S4**) were used to build the synteny-improved assemblies and compute scaffold N50 values and counts before and after merging. As the synteny-based methods rely on orthologous anchors as their input data they cannot predict adjacencies for scaffolds with no annotated orthologous genes, thus N50 values and counts were computed based only on scaffolds with annotated orthologues (**Fig. 2, main text; Figures S5 and S6**). Linear regressions plotted with 95% confidence intervals computed with the geom_smooth() function from the R package ggplot2, specifying the 'lm' method.

**Table S4. Synteny-based adjacency agreements**

Counts of input (All) and filtered (Use) adjacencies from three synteny-based methods and their agreements or conflicts, reported two-way agreements are required not to conflict with the third method.

| Species | ADSEQ | | GOS-ASM | | ORTHOSTITCH | | 3-Way Agreement | 2-Way Agreement | ADSEQ & GOS-ASM | GOS-ASM & ORTHOSTITCH | ADSEQ & ORTHOSTITCH |
|---|---|---|---|---|---|---|---|---|---|---|---|
| | All | Use | All | Use | All | Use | | | | | |
| *Anopheles albimanus* | 1 | 1 | 2 | 2 | 4 | 4 | 0 | 1 | 0 | 1 | 0 |
| *Anopheles arabiensis* | 48 | 48 | 11 | 11 | 33 | 29 | 2 | 19 | 3 | 0 | 16 |
| *Anopheles atroparvus* | 42 | 42 | 6 | 6 | 29 | 25 | 1 | 14 | 1 | 0 | 13 |
| *Anopheles christyi* | 3220 | 3220 | 1176 | 1176 | 2031 | 1937 | 474 | 1041 | 238 | 17 | 786 |
| *Anopheles coluzzii* | 199 | 199 | 71 | 71 | 134 | 123 | 27 | 54 | 8 | 1 | 45 |
| *Anopheles culicifacies* | 3594 | 3594 | 2055 | 2055 | 1821 | 1742 | 659 | 910 | 494 | 29 | 387 |
| *Anopheles darlingi* | 678 | 678 | 290 | 290 | 684 | 640 | 102 | 281 | 31 | 23 | 227 |
| *Anopheles dirus* | 63 | 63 | 19 | 19 | 42 | 39 | 9 | 27 | 5 | 0 | 22 |
| *Anopheles epiroticus* | 608 | 608 | 143 | 143 | 471 | 456 | 74 | 327 | 28 | 3 | 296 |
| *Anopheles farauti* | 177 | 177 | 75 | 75 | 119 | 116 | 43 | 62 | 12 | 2 | 48 |
| *Anopheles funestus* | 331 | 331 | 100 | 100 | 211 | 208 | 47 | 171 | 32 | 2 | 137 |
| *Anopheles gambiae* | 0 | 0 | 0 | 0 | 0 | 0 | 0 | 0 | 0 | 0 | 0 |
| *Anopheles maculatus* | 6366 | 6366 | 1859 | 1859 | 2411 | 2312 | 377 | 1076 | 401 | 26 | 649 |
| *Anopheles melas* | 5081 | 5081 | 3080 | 3080 | 2181 | 2116 | 773 | 1075 | 696 | 36 | 343 |
| *Anopheles merus* | 590 | 590 | 220 | 220 | 422 | 413 | 118 | 254 | 35 | 9 | 210 |
| *Anopheles minimus* | 19 | 19 | 7 | 7 | 14 | 14 | 3 | 9 | 3 | 0 | 6 |
| *Anopheles quadriannulatus* | 238 | 238 | 122 | 122 | 163 | 148 | 49 | 81 | 23 | 2 | 56 |
| *Anopheles sinensis* | 336 | 335 | 196 | 196 | 218 | 204 | 30 | 90 | 43 | 7 | 40 |
| *Anopheles sinensis (Chinese)* | 158 | 158 | 166 | 166 | 78 | 77 | 27 | 65 | 45 | 5 | 15 |
| *Anopheles stephensi* | 277 | 277 | 106 | 106 | 182 | 177 | 53 | 124 | 24 | 3 | 97 |
| *Anopheles stephensi (Indian)* | 201 | 201 | 69 | 69 | 155 | 144 | 40 | 90 | 12 | 2 | 76 |

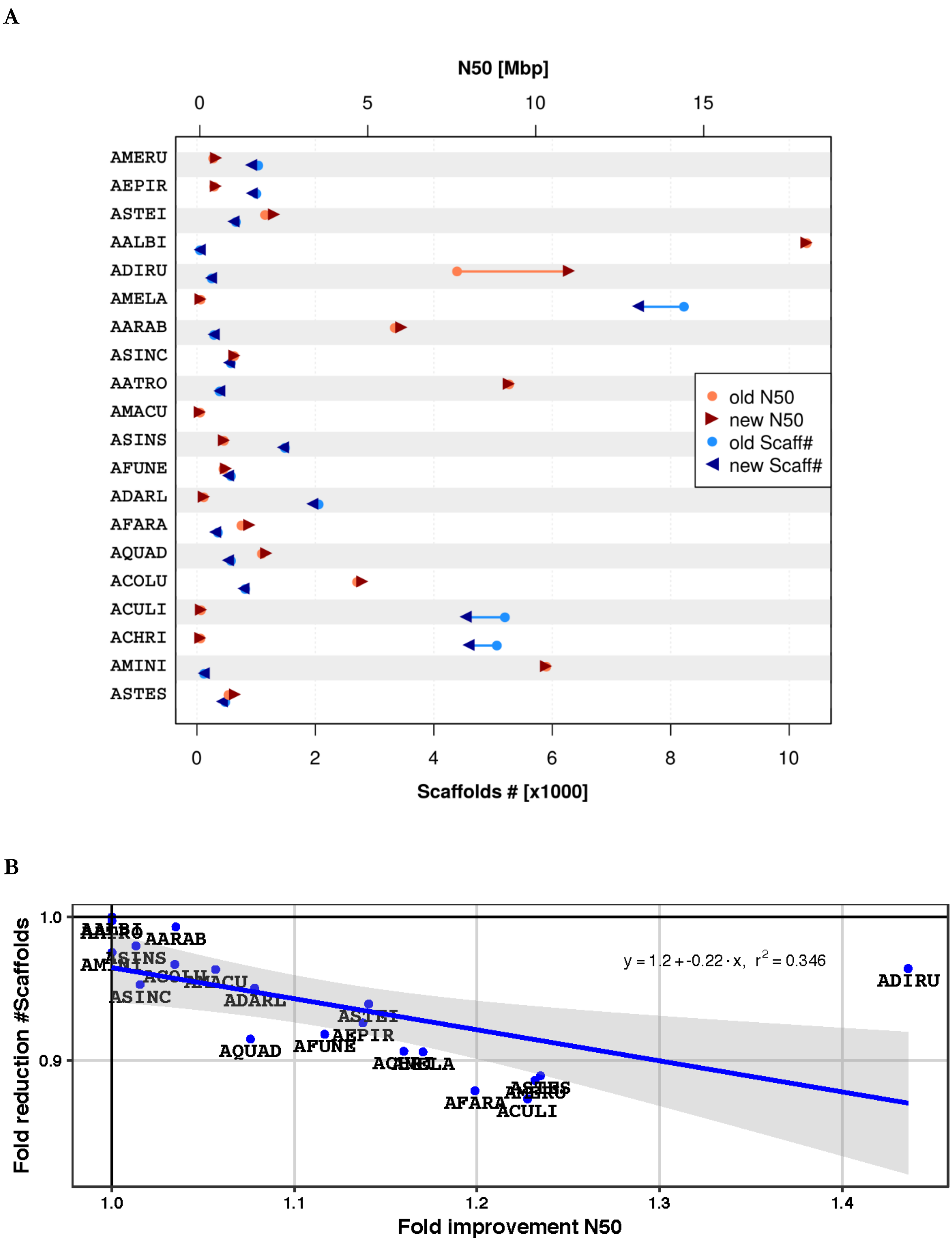

**Figure S5. Assembly improvements based on conservative set synteny predictions**

For details see Fig. 2, main text.

A

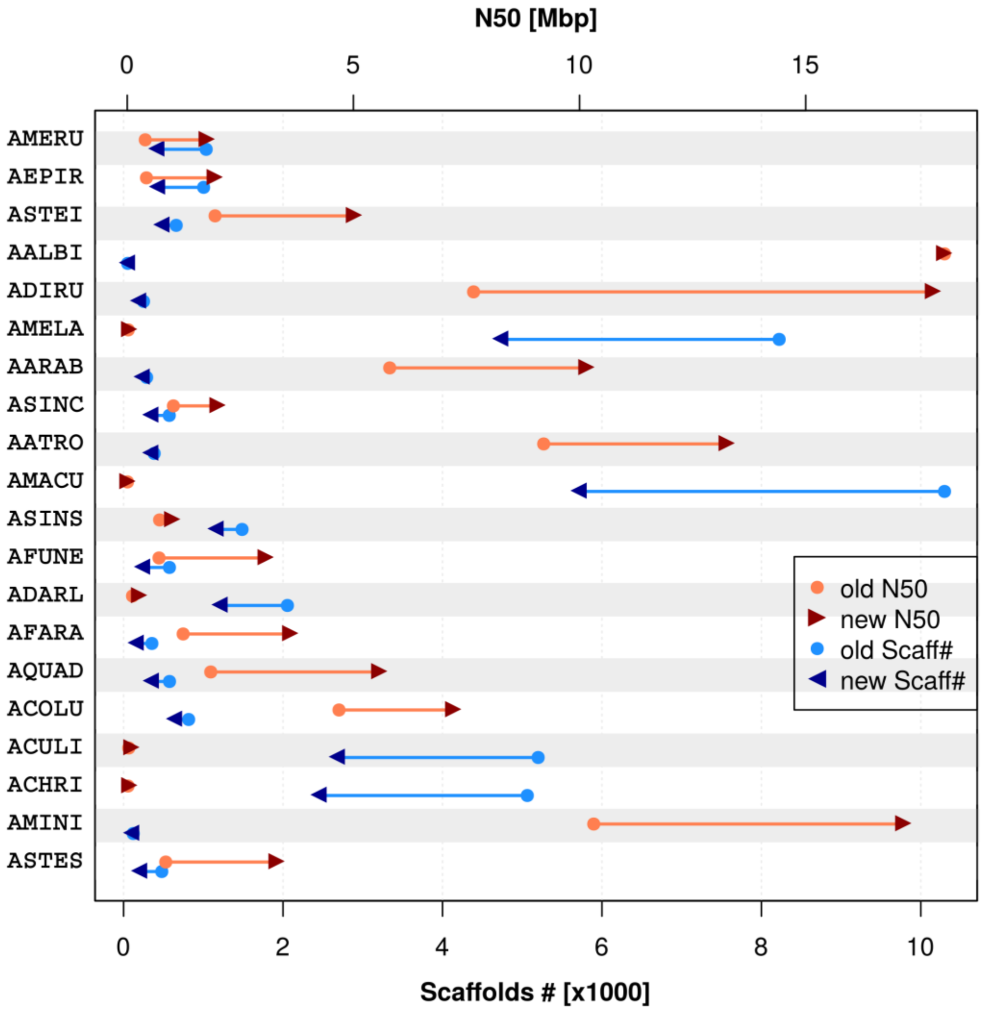

B

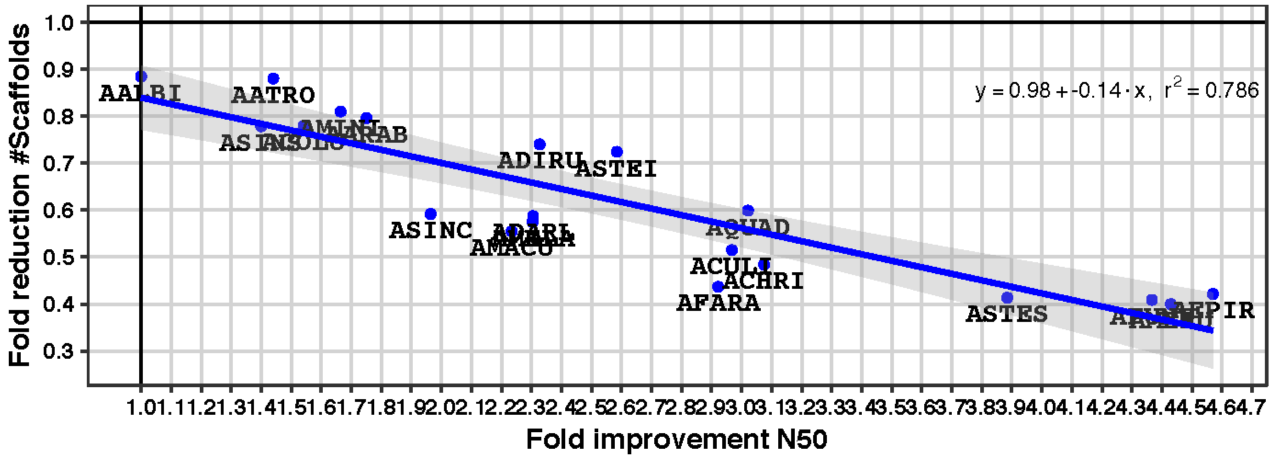

**Figure S6. Assembly improvements based on liberal union set synteny predictions**
For details see Fig. 2, main text.

### Synteny-based method comparisons

Comparing the CAMSA-produced two-way consensus sets with the input adjacencies from each of the three methods quantified agreements (**Table S4**) as well as conflicting and unique adjacencies predicted by each method for each assembly (**Fig. 3, main text; Figure S7**). A total of 29'418 distinct scaffold adjacencies were identified from the combined results of all 42'923 predictions from the three methods. These were classified according to whether they were in three-way agreement, in two-way agreement with no third-method conflict, in two-way agreement but with conflict(s), unique to an individual method with no conflict(s) with the other methods, or unique to an individual method but with conflict(s).

Comparing all 42'923 predictions identified 29'418 distinct scaffold adjacencies, 36% of which were supported by at least two methods. Overall, 10% of the distinct adjacencies were predicted by all three methods, and a further 26% were predicted by two methods but this was reduced to 20% when adjacencies that conflicted with the third method were removed. These 8'878 supported predictions were used to build the two-way consensus sets of scaffold adjacencies for synteny-based assembly improvements presented in **Fig. 2**. Main text **Fig. 3B** shows the overlaps amongst the three methods, plotted as an area-proportional Euler diagram with EULERAPE v3.0.0 (Micallef and Rodgers 2014). Adjacencies in three-way agreement made up 30% of GOS-ASM and 27% of ORTHOSTITCH predictions, and 13% of ADSEQ predictions (as there were about double the number of ADSEQ predictions compared with the other two methods). The much larger total number of ADSEQ predictions resulted in a higher proportion of unique adjacencies (54%) compared with GOS-ASM (35%) and ORTHOSTITCH (31%). Pairwise method comparisons: ADSEQ supported 61% of GOS-ASM and 65% of ORTHOSTITCH predictions; ORTHOSTITCH supported 32% of ADSEQ and 34% of GOS-ASM adjacencies; and GOS-ASM supported 27% of ADSEQ and 30% of ORTHOSTITCH predictions.

Considering only the liberal union sets of all non-conflicting adjacencies, the adjacencies in three-way agreement made up 16.5% of the total, 45.6% of GOS-ASM, 39.1% of ORTHOSTITCH, and 18.6% of ADSEQ predictions (**Fig. 3B, main text**). From the two-way consensus adjacency sets with no third-method conflicts, three-way consensus adjacencies made up 32.8% of the total, 53.8% of GOS-ASM, 44.4% of ORTHOSTITCH, and 33.4% of ADSEQ predictions (**Fig. 3B, main text**). These two-way consensus adjacencies that were employed to build the new superscaffolded assemblies were therefore supported by ADSEQ (98.1%), and/or ORTHOSTITCH (73.7%), and/or GOS-ASM (60.9%), with a third being supported by all three methods. Thus, comparing the results from the three methods and employing a two-way agreement with no third-method conflict filter improved the overall level of three-way agreement from a tenth to a third.

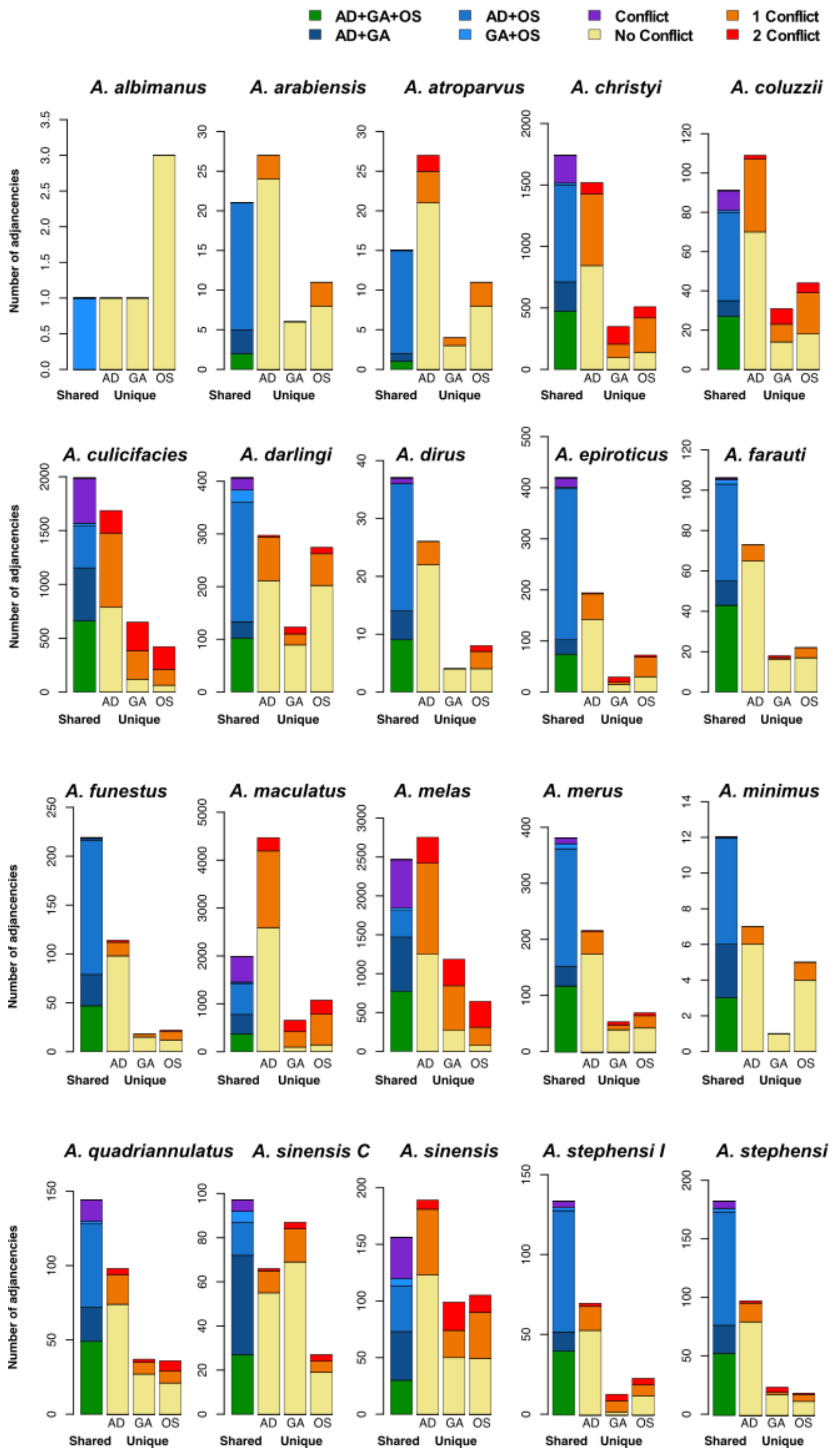

**Figure S7. Comparisons of adjacency results from three synteny-based methods**

Comparisons of synteny-based scaffold adjacency predictions from ADSEQ (AD), GOS-ASM (GA), and ORTHOSTITCH (OS). Bar charts show counts of predicted adjacencies (pairs of neighbouring scaffolds) that are shared amongst all three methods (green), or two methods without (blues) and with (purple) third method conflicts, or that are unique to a single method and do not conflict (yellow) or do conflict with predictions from one (orange) or both (red) of the other methods. Note variable maxima for y-axes.

Examining the results from each individual assembly (selected assemblies shown **Fig. 3C, main text**; all assemblies shown in **Figures S7 and S8**), showed generally good agreement for at least eight of the assemblies (more than 48% of distinct adjacencies were found to be in at least two-way agreement with no third-method conflict), with *A. funestus* achieving the highest consistency at 58%. Some of the most fragmented input assemblies produced the some of the largest sets of distinct adjacency predictions but the agreement amongst these predictions was generally lower than the other assemblies, e.g. *A. maculatus* with 8'179 distinct adjacencies of which only of which only 18% showed at least two-way agreement with no conflicts (**Figure S8**). *A. albimanus* showed a very low level of agreement (16.7%), but this is primarily because of the very few predicted adjacencies: just six distinct adjacencies with only one being shared between two of the methods.

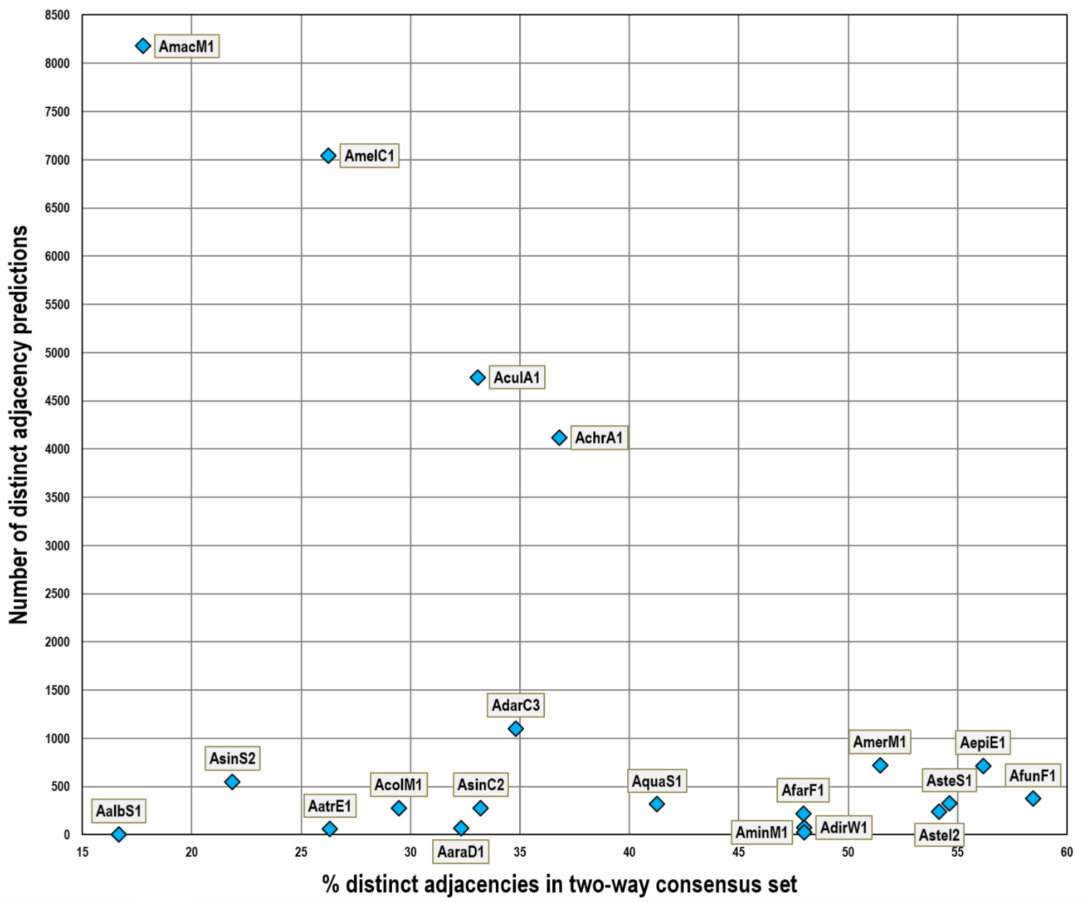

**Figure S8. Proportions of synteny-based adjacencies in agreement for each assembly**
Comparisons of the number of distinct adjacencies and the proportion of which were common to at least two methods with no third method conflict. Two-way consensus adjacencies made up 48% or more of the distinct predictions for eight assemblies, while some of the most fragmented assemblies with the most predicted adjacencies showed lower levels of agreement. For AalbS1 (*Anopheles albimanus*), only one of the six distinct adjacency predictions was in the two-way consensus set. See **Table S2** for the species that corresponds to each assembly identifier.

### [8] Physical mapping data from six anophelines

*Jiyoung Lee, Phillip George, Maryam Kamali, Ashley Peery, Maria V. Sharakhova, Maria F. Unger, Igor V. Sharakhov*

Methods of chromosomal mapping of scaffolds (Sharakhova et al. 2019; Artemov et al. 2018b) are detailed for *A. albimanus* (Artemov et al. 2017), *A. atroparvus* (Artemov et al. 2015; Neafsey et al. 2015; Artemov et al. 2018a), *A. sinensis* Chinese strain (Wei et al. 2017), *A. stephensi* SDA-500 strain (Neafsey et al. 2015), and *A. stephensi* Indian strain (Jiang et al. 2014). *A. stephensi* mapping added to existing mapping data (Sharakhova et al. 2006, 2010), and *A. funestus* mapping built on previous results (Sharakhov et al. 2002, 2004; Xia et al. 2010) to further develop the physical map as described in detail below. Counts of mapped scaffolds and the resulting scaffold adjacencies, i.e. pairs of neighbouring mapped scaffolds, are summarised in **Table S5**.

**Table S5. Physically mapped scaffolds from six anophelines**
Counts of physically mapped scaffolds and adjacencies available for six of the anophelines.

| Species | Number of Mapped Scaffolds | Usable Scaffold Pair Adjacencies | Reference(s) |
|---|---|---|---|
| *Anopheles albimanus* | 31 | 31 | (Artemov et al. 2017) |
| *Anopheles atroparvus* | 46 | 31 | (Artemov et al. 2015; Neafsey et al. 2015; Artemov et al. 2018a) |
| *Anopheles funestus* | 202 | 85 | (Sharakhov et al. 2002, 2004; Xia et al. 2010; Neafsey et al. 2015) & this study |
| *Anopheles sinensis (Chinese)* | 52 | 20 | (Wei et al. 2017) |
| *Anopheles stephensi* | 99 | 3 | (Neafsey et al. 2015) |
| *Anopheles stephensi (Indian)* | 118 | 6 | (Jiang et al. 2014) & this study |

**Mosquito strain and ovary preservation:**

The FUMOZ strain of *A. funestus* was maintained in the insectary of the Eck Institute, the University of Notre Dame USA. The strain was originally colonized from the Matolo Province of Mozambique, and deposited at the Malaria Research and Reference Reagent Resource (MR4) at the Biodefense and Emerging Infections Research Resources Repository (BEI) under catalogue number MRA-1027. Mosquitoes were raised in a growth chamber at 27°C, with a 12-hour cycle of light and darkness. Approximately 20-21 hours post-blood feeding, ovaries of adult females were pulled out and fixed in Carnoy's solution (3 : 1 ethanol : glacial acetic acid by volume). Ovaries were preserved in fixative solution from 24 h up to 1 month at -20°C.

**Chromosome preparation:**

Isolated ovaries were bathed in a drop of 50% propionic acid for 5 minutes and squashed as previously described (Sharakhova et al. 2014). The quality of the preparation was assessed with an Olympus CX41 phase contrast microscope (Olympus America Inc., Melville, NY). High-quality chromosome preparations were then flash frozen in liquid nitrogen and immediately placed in cold 50% ethanol. After that, preparations were dehydrated in an ethanol series (50%, 70%, 90%, and 100%) and air-dried. Unstained chromosomes were observed using an Olympus BX41 phase contrast microscope with attached CCD camera Qcolor5 (Olympus America Inc., Melville, NY).

**Probe preparation and fluorescence *in situ* hybridization:**

Gene-specific primers were designed to amplify unique exon sequences from the beginning and end of each scaffold using the primer-BLAST program (Ye et al. 2012) available at NCBI (http://www.ncbi.nlm.nih.gov/tools/primer-blast/). The primer design was based on gene annotations from the AfunF1 genome assembly available at VECTORBASE (https://www.vectorbase.org/organisms/anopheles-funestus/fumoz/afunf1) (Giraldo-Calderón et al. 2015). PCR was performed using 2X Immomix DNA polymerase (Bioline USA Inc., MA, USA) and a standard Immomix amplification protocol. Amplified fragments were labelled with fluorescein, Cy3 or Cy5 dyes (GE Health Care, UK Ltd, Buckinghamshire, UK and Enzo Biochem, Enzo Life Sciences Inc., Farmingdale, NY) using a Random Primers DNA Labelling System (Invitrogen, Carlsbad, CA, USA). By combining different dyes in one reaction, we labelled and used up to four probes corresponding to two genomic scaffolds in the same FISH experiment. FISH was performed according to the previously described standard protocol (Sharakhova et al. 2014). DNA probes were hybridized to the chromosomes at 39°C for 10-15 hours in a hybridization solution (50% Formamide; 10% Sodium Dextran sulfate, 0.1% Tween 20 in 2XSSC, pH 7.4). Chromosome preparations were washed in 0.2X SSC (Saline-Sodium Citrate: 0.03M Sodium Chloride, 0.003M Sodium Citrate) and counterstained with DAPI in ProLong Gold Antifade Mountant (Thermo Fisher Scientific Inc., USA).

**Linking Illumina scaffolds with PacBio contigs and PacBio merged scaffolds:**

Illumina scaffolds of the *A. funestus* Anop_fune_FUMOZ_V1 assembly were downloaded from GENBANK (https://www.ncbi.nlm.nih.gov/assembly/GCA_000349085.1). PacBio contigs which were longer than one million bp were aligned to the Illumina scaffolds using BLASTN 2.2.31+ (http://blast.ncbi.nlm.nih.gov/Blast.cgi) with default settings. The PacBio assembly was generated with approximately 70X of PacBio sequencing data and polished by Quiver (see section below on 'Building the PacBio-based *Anopheles funestus* assembly'). To see the reverse alignment relationships, we used Illumina scaffolds as query sequences and align them to PacBio merged scaffolds with the standalone BLASTN 2.2.30+ program installed on a server, and we built BLAST databases. The PacBio merged scaffolds were obtained by merging PacBio contigs with the Illumina assembly using METASSEMBLER (Wences and Schatz 2015) and then by scaffolding with SSPACE (Boetzer et al. 2011) using available Illumina sequencing data. The BLASTN was performed with the 97% identity and 1e-50 e-value thresholds. Illumina and PacBio merged scaffolds longer than 0.2 million bps were chosen for FISH (**Figure S9**). CIRCOLETTO was used to visualize sequence similarity between linked Illumina scaffolds with merged PacBio scaffolds, their order and orientation (Darzentas 2010). Illumina scaffolds were ordered and oriented within large PacBio contigs and merged PacBio scaffolds, and the resulted arrangements were anchored to chromosomes by FISH as described above.

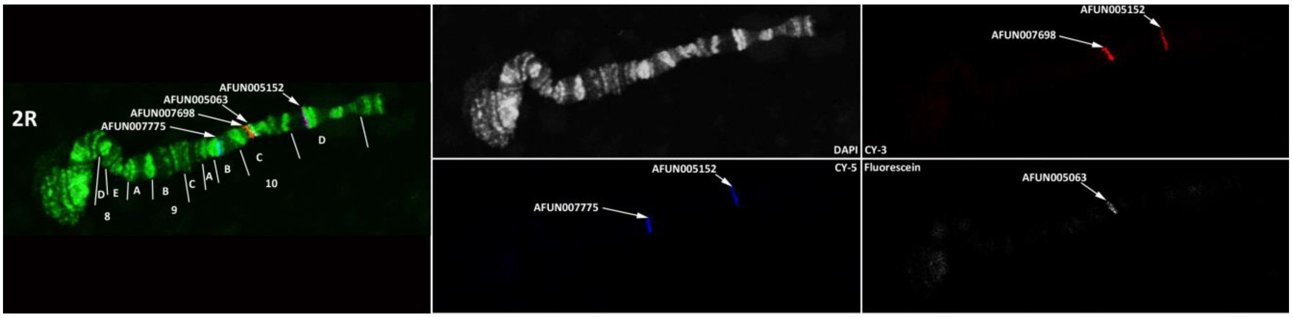

**Figure S9. Fluorescence *in situ* hybridization (FISH) mapping in *Anopheles funestus*.**
Multicolour FISH of four DNA probes designed based on gene sequences. Polytene chromosomes are from ovarian nurse cells of *A. funestus*.

**Chromosome mapping:**

Illumina scaffolds and merged Illumina-PacBio arrangements were anchored to chromosomes by several different ways. (1) Scaffolds without adjacency and orientation were placed on chromosomes with only one FISH probe. (2) Oriented scaffolds without adjacency were placed on chromosomes with at least two FISH probes, but they did not have any neighbours. (3) Scaffolds with adjacency but without orientation consisted of two or several neighbouring scaffolds mapped with one FISH probe each. Alternatively, several Illumina scaffolds were predicted to be adjacent within a PacBio contig or PacBio-merged scaffolds by BLAST but the whole assembly was anchored to chromosome by only one FISH probe. (4) Ordered and oriented scaffolds were placed on chromosomes by multiple FISH probes (**Figure S9**) or their adjacency and orientation were inferred from the alignment to a mapped and oriented PacBio contigs or PacBio-merged scaffolds. The resulting physical genome map for *A. funestus* includes 202 AfunF1 scaffolds (**Table S6**).

**Table S6. Physically mapped *Anopheles funestus* scaffolds on the cytogenetic map**

Chromosomal locations and orientation (if determined) of AfunF1 genomic scaffolds on the *Anopheles funestus* cytogenetic map from 126 previously FISH-mapped (Sharakhov et al. 2002, 2004; Xia et al. 2010) and 66 newly FISH-mapped DNA markers. Note that these mappings incorporate additions and corrections that were made during the reconciliation process and thus there are some differences with the 'input' physical mapping data.

| AfunF1 scaffolds | Scaffold orientation | Chromosome region | Scaffold size | Method of placement to the map |
|---|---|---|---|---|
| KB668763 | - | X:1A | 305818 | PACBIO OVERLAP |
| KB669058 | + | X:1AB | 2367365 | MAPPED |
| KB668322 | - | X:1B | 615127 | MAPPED |
| KB668245 | - | X:1C | 629234 | MAPPED |
| KB669125 | - | X:1C | 833292 | MAPPED |
| KB669181 | - | X:1C | 789512 | PACBIO OVERLAP |
| KB668600 | - | X:1D | 671960 | MAPPED |
| KB668720 | + | X:1D | 419310 | PACBIO OVERLAP |
| KB668844 | + | X:1D | 215519 | PACBIO OVERLAP |
| KB668852 | + | X:1D | 216557 | MAPPED |
| KB668755 | + | X:2A | 379841 | PACBIO OVERLAP |
| KB668367 | ? | X:2B | 636359 | MAPPED |
| KB668936 | + | X:2BC | 1176300 | MAPPED |
| KB669143 | - | X:2C | 12834 | PACBIO OVERLAP |
| KB669145 | - | X:3A | 12715 | PACBIO OVERLAP |
| KB668668 | - | X:3AB | 583467 | MAPPED |
| KB669003 | - | X:3CD | 1206901 | MAPPED |
| KB668797 | + | X:3D | 250364 | PACBIO OVERLAP |
| KB668522 | + | X:4A | 703988 | MAPPED |
| KB669029 | + | X:4A | 88731 | PACBIO OVERLAP |
| KB669078 | - | X:4AB | 51937 | PACBIO OVERLAP |
| KB668688 | ? | X:5C | 429614 | MAPPED |
| KB668389 | ? | X:5C | 547300 | PACBIO OVERLAP |
| KB668765 | ? | X:6 | 305606 | PACBIO OVERLAP |
| KB668660 | ? | X:6 | 504041 | PACBIO OVERLAP |
| KB668760 | ? | X:6 | 504041 | MAPPED |
| KB669536 | ? | X:6 | 625123 | MAPPED |
| KB668728 | ? | 2R:7A | 333164 | MAPPED |
| KB668825 | ? | 2R:7BC | 1174813 | MAPPED |
| KB668221 | + | 2R:8AE | 3832769 | MAPPED |
| KB668954 | + | 2R:9A | 66343 | PACBIO OVERLAP |
| KB669004 | + | 2R:9A | 57566 | PACBIO OVERLAP |
| KB669169 | + | 2R:9A-10B | 1771395 | MAPPED |
| KB668759 | - | 2R:10BC | 1353732 | MAPPED |
| KB668737 | + | 2R:10C | 1261231 | MAPPED |
| KB668555 | - | 2R:10D-11A | 1493947 | MAPPED |
| KB668845 | ? | 2R:11B | 188083 | MAPPED |
| KB668753 | - | 2R:11C | 343795 | PACBIO OVERLAP |
| KB668871 | + | 2R:11C | 218729 | PACBIO OVERLAP |
| KB668467 | - | 2R:12A | 563073 | MAPPED |
| KB669369 | + | 2R:12B | 804489 | MAPPED |
| KB668822 | ? | 2R:12B | 194798 | MAPPED |
| KB668793 | ? | 2R:12C | 254162 | MAPPED |
| KB668745 | ? | 2R:12C | 346721 | PACBIO OVERLAP |

| | | | | |
|---|---|---|---|---|
| KB668672 | - | 2R:12D | 584724 | MAPPED |
| KB668775 | - | 2R:12D | 355205 | PACBIO OVERLAP |
| KB668785 | - | 2R:12E | 408060 | MAPPED |
| KB669081 | - | 2R:12E-13A | 1051734 | MAPPED |
| KB668706 | ? | 2R:13A | 572386 | MAPPED |
| KB668715 | ? | 2R:13B | 364622 | MAPPED |
| KB668766 | ? | 2R:13C | 300363 | MAPPED |
| KB668757 | ? | 2R:13C | 303086 | PACBIO OVERLAP |
| KB668411 | ? | 2R:13CD | 533316 | PACBIO OVERLAP |
| KB668679 | ? | 2R:13D | 436776 | MAPPED |
| KB668478 | ? | 2R:14B | 598359 | MAPPED |
| KB669525 | ? | 2R:14C | 624454 | MAPPED |
| KB668835 | - | 2R:14D | 178699 | MAPPED |
| KB669358 | - | 2R:15B | 786284 | MAPPED |
| KB668911 | ? | 2R:15C | 97490 | MAPPED |
| KB669547 | ? | 2R:15E | 618077 | MAPPED |
| KB668914 | + | 2R:15E-16A | 1038133 | MAPPED |
| KB668837 | ? | 2R:16A | 1126662 | MAPPED |
| KB669192 | ? | 2R:16B | 922711 | MAPPED |
| KB668748 | ? | 2R:16C | 1398093 | MAPPED |
| KB668670 | - | 2R:17AB | 1428115 | MAPPED |
| KB668947 | + | 2R:17C-18A | 2413216 | MAPPED |
| KB668742 | - | 2R:18A | 317854 | PACBIO OVERLAP |
| KB668234 | + | 2R:18AB | 602066 | MAPPED |
| KB668734 | - | 2R:18C | 433875 | MAPPED |
| KB668836 | + | 2R:18CD | 2772343 | MAPPED |
| KB668289 | - | 2R:18D | 644640 | PACBIO OVERLAP |
| KB669247 | + | 2R:19A | 782462 | PACBIO OVERLAP |
| KB668866 | - | 2R:19B | 140173 | PACBIO OVERLAP |
| KB669114 | - | 2R:19C | 878476 | MAPPED |
| KB668942 | - | 2R:19C | 101067 | PACBIO OVERLAP |
| KB668870 | - | 2R:19CD | 1070199 | MAPPED |
| KB668694 | - | 2R:19DE | 522677 | PACBIO OVERLAP |
| KB669281 | ? | 2L:20BC | 897800 | MAPPED |
| KB669092 | ? | 2L:20C | 887634 | MAPPED |
| KB669070 | ? | 2L:20D | 943178 | MAPPED |
| KB668589 | + | 2L:21BC | 555648 | PACBIO OVERLAP |
| KB668770 | + | 2L:21C | 1246493 | MAPPED |
| KB668781 | - | 2L:21CD | 1311501 | MAPPED |
| KB668222 | - | 2L:21D-22A | 1876834 | MAPPED |
| KB668692 | - | 2L:22AC | 1835194 | MAPPED |
| KB668872 | - | 2L:22C | 248811 | MAPPED |
| KB669502 | ? | 2L:22D | 1948688 | MAPPED |
| KB668882 | + | 2L:23A | 121550 | MAPPED |
| KB668803 | + | 2L:24AB | 1311425 | MAPPED |
| KB669036 | + | 2L:24B | 850007 | PACBIO OVERLAP |
| KB668433 | - | 2L:24B | 552028 | PACBIO OVERLAP |
| KB669280 | + | 2L:24CD | 1738428 | MAPPED |
| KB668854 | ? | 2L:26A | 204352 | PACBIO OVERLAP |
| KB669047 | ? | 2L:26A | 999242 | MAPPED |
| KB668681 | + | 2L:26C | 1609593 | MAPPED |
| KB668702 | - | 2L:26C | 460558 | PACBIO OVERLAP |
| KB668764 | - | 2L:26CD | 327039 | MAPPED |
| KB668693 | + | 2L:26D | 518197 | MAPPED |
| KB668278 | - | 2L:27A | 702492 | PACBIO OVERLAP |
| KB669136 | - | 2L:27AB | 882720 | MAPPED |

| | | | | |
|---|---|---|---|---|
| KB668813 | ? | 2L:27C | 258639 | PACBIO OVERLAP |
| KB668795 | ? | 2L:27C | 281738 | MAPPED |
| KB669214 | ? | 2L:27CD | 874018 | PACBIO OVERLAP |
| KB668992 | ? | 2L:27D | 953533 | MAPPED |
| KB668751 | ? | 2L:27E | 335862 | MAPPED |
| KB668892 | ? | 2L:27E | 1102954 | PACBIO OVERLAP |
| KB668725 | ? | 2L:28A | 3134932 | MAPPED |
| KB668881 | ? | 2L:28C | 976588 | MAPPED |
| KB668378 | ? | 3R:29B | 571908 | MAPPED |
| KB669236 | ? | 3R:29C | 713413 | MAPPED |
| KB668851 | ? | 3R:29C | 164446 | PACBIO OVERLAP |
| KB668683 | ? | 3R:29CD | 458860 | MAPPED |
| KB669458 | - | 3R:29D-30A | 679292 | MAPPED |
| KB668792 | ? | 3R:30AC | 1431544 | MAPPED |
| KB668633 | ? | 3R:30C | 577339 | MAPPED |
| KB669089 | ? | 3R:30C | 19591 | PACBIO OVERLAP |
| KB668687 | ? | 3R:30C | 589177 | PACBIO OVERLAP |
| KB668808 | + | 3R:30C | 233839 | MAPPED |
| KB668705 | + | 3R:30CD | 400982 | PACBIO OVERLAP |
| KB668695 | ? | 3R:31C | 480392 | PACBIO OVERLAP |
| KB668812 | ? | 3R:31CD | 217759 | MAPPED |
| KB668644 | ? | 3R:32B | 475666 | MAPPED |
| KB669580 | ? | 3R:32B | 640366 | PACBIO OVERLAP |
| KB668789 | ? | 3R:33A | 318576 | MAPPED |
| KB668661 | + | 3R:33C | 494022 | MAPPED |
| KB668533 | ? | 3R:33C | 534943 | MAPPED |
| KB669347 | ? | 3R:33D | 864369 | MAPPED |
| KB668723 | ? | 3R:33D | 382759 | MAPPED |
| KB668818 | ? | 3R:34A | 257953 | PACBIO OVERLAP |
| KB668746 | ? | 3R:34A | 323294 | MAPPED |
| KB668750 | - | 3R:34B | 390086 | PACBIO OVERLAP |
| KB668848 | - | 3R:34BC | 1452984 | MAPPED |
| KB668671 | ? | 3R:35B | 710502 | MAPPED |
| KB668853 | ? | 3R:35B | 306342 | MAPPED |
| KB668790 | ? | 3R:35C | 381342 | MAPPED |
| KB668700 | - | 3R:35CD | 478136 | PACBIO OVERLAP |
| KB668684 | + | 3R:35D | 499467 | PACBIO OVERLAP |
| KB669103 | - | 3R:35DE | 1272063 | MAPPED |
| KB669336 | + | 3R:35E | 1014753 | MAPPED |
| KB668456 | + | 3R:35EF | 708050 | MAPPED |
| KB669011 | - | 3R:35F | 39851 | MAPPED |
| KB668709 | - | 3R:35F | 491865 | MAPPED |
| KB668752 | - | 3R:35F | 447341 | PACBIO OVERLAP |
| KB668816 | + | 3R:35F | 290561 | MAPPED |
| KB668880 | + | 3R:35F | 119321 | MAPPED |
| KB669403 | + | 3R:35F | 972476 | MAPPED |
| KB669034 | + | 3R:35F | 31323 | PACBIO OVERLAP |
| KB668756 | - | 3R:35F | 460866 | PACBIO OVERLAP |
| KB668777 | - | 3R:36A | 381111 | MAPPED |
| KB668731 | - | 3R:36A | 405878 | MAPPED |
| KB668779 | - | 3R:36A | 398220 | PACBIO OVERLAP |
| KB668987 | - | 3R:36AB | 46664 | PACBIO OVERLAP |
| KB669414 | + | 3R:36B | 997879 | PACBIO OVERLAP |
| KB668810 | + | 3R:36B | 310469 | PACBIO OVERLAP |
| KB669380 | + | 3R:36B | 770134 | PACBIO OVERLAP |
| KB668858 | + | 3R:36B | 254377 | PACBIO OVERLAP |

| | | | | |
|---|---|---|---|---|
| KB668667 | + | 3R:36B | 580684 | MAPPED |
| KB668806 | + | 3R:36C | 292235 | MAPPED |
| KB668874 | + | 3R:36C | 134456 | MAPPED |
| KB668959 | + | 3R:36D | 938304 | MAPPED |
| KB668732 | + | 3R:36D | 346637 | MAPPED |
| KB668999 | - | 3R:36D | 106097 | PACBIO OVERLAP |
| KB669391 | - | 3R:36DE | 2124680 | MAPPED |
| KB669436 | - | 3R:36E | 911299 | MAPPED |
| KB668819 | - | 3R:36E | 222314 | PACBIO OVERLAP |
| KB668744 | - | 3R:36E | 359052 | PACBIO OVERLAP |
| KB669469 | + | 3R:36F | 758275 | MAPPED |
| KB669203 | + | 3R:36F | 774987 | PACBIO OVERLAP |
| KB668970 | + | 3R:37A | 879108 | PACBIO OVERLAP |
| KB668726 | + | 3R:37AB | 1292245 | MAPPED |
| KB668762 | - | 3R:37C | 351750 | PACBIO OVERLAP |
| KB668805 | - | 3L:38A | 229600 | MAPPED |
| KB668859 | + | 3L:38B | 1041161 | MAPPED |
| KB668823 | ? | 3L:38C | 194609 | MAPPED |
| KB669325 | ? | 3L:39A | 699719 | MAPPED |
| KB668717 | ? | 3L:39A | 402095 | MAPPED |
| KB668578 | ? | 3L:39A | 505528 | MAPPED |
| KB668676 | ? | 3L:39A | 439401 | PACBIO OVERLAP |
| KB668659 | ? | 3L:39B | 1545011 | MAPPED |
| KB669014 | ? | 3L:40A | 892505 | MAPPED |
| KB668773 | ? | 3L:40A | 296798 | PACBIO OVERLAP |
| KB668868 | ? | 3L:40B | 141784 | MAPPED |
| KB668918 | ? | 3L:40B | 93149 | MAPPED |
| KB668444 | ? | 3L:41A | 1547433 | MAPPED |
| KB668754 | ? | 3L:41D | 348546 | MAPPED |
| KB668830 | ? | 3L:41D | 181046 | MAPPED |
| KB668333 | + | 3L:42AB | 1525199 | MAPPED |
| KB668422 | ? | 3L:42D | 532383 | MAPPED |
| KB668500 | ? | 3L:43A | 547302 | MAPPED |
| KB668925 | ? | 3L:43B | 940405 | MAPPED |
| KB669025 | ? | 3L:44B | 910511 | MAPPED |
| KB669264 | ? | 3L:44B | 32220 | MAPPED |
| KB669603 | ? | 3L:44C | 2248 | MAPPED |
| KB668849 | ? | 3L:44C | 159962 | MAPPED |
| KB668784 | ? | 3L:45A | 266933 | MAPPED |
| KB668948 | - | 3L:46B | 917728 | PACBIO OVERLAP |
| KB668682 | - | 3L:46B | 470092 | MAPPED |
| KB668714 | - | 3L:46BC | 1305928 | MAPPED |
| KB668703 | - | 3L:46CD | 1311118 | MAPPED |
| KB669207 | ? | 3L:46D | 62723 | MAPPED |
| KB668252 | ? | 3L:46D | 2000 | MAPPED |
| KB668265 | ? | 3L:46D | 1954 | MAPPED |

As for the comparisons of the synteny-based results, CAMSA was used to compare the two-way consensus sets, as well as the conservative three-way consensus sets and the liberal union sets of all non-conflicting adjacencies, with the physical mapping adjacencies from each of the six assemblies and quantify agreements as well as conflicting and unique adjacencies (**Table S7**).

For *A. albimanus*, the two-way consensus synteny-based predictions produced only a single adjacency, and this was confirmed by the physical mapping data. Five of the 15 two-way consensus synteny-based predictions were confirmed by physical mapping of *A. atroparvus* scaffolds and only one conflict (resolved) was identified (**Fig. 4A, main text**). The mapped scaffolds for the *A. stephensi* assemblies resulted very few adjacencies, the three SDA-500 strain adjacencies were all in conflict with synteny-based predictions, and of the six Indian strain adjacencies three were shared and one was in conflict with the two-way consensus synteny-based predictions. These conflicts were resolved by correcting the orientations of the physically mapped scaffolds, as the probe designs meant that mapping misorientations were possible.

Comparing the 20 *A. sinensis* (Chinese) mapped scaffolds confirmed three of the synteny-based adjacencies, but none of these were in the consensus sets, and identified conflicts with just two of the 92 two-way consensus adjacencies, both of which were resolved as they involved scaffolds that had not been selected for physical mapping. And finally, *A. funestus* presented the most adjacencies from both physical mapping and the synteny-based predictions where 12-17% of the different sets of synteny-based adjacencies were confirmed and just 4-8% were in conflict (**Fig. 4A, main text**). Amongst the 14 physically mapped neighbouring pairs that conflicted with 13 synteny-based adjacencies from the two-way consensus set, five conflicts were resolved because the synteny-based neighbour was short and not used for physical mapping. An additional four conflicts were resolved by switching the orientation of physically mapped scaffolds, which were anchored by only a single FISH probe and therefore their orientations were not confidently determined. All but one of these adjacency conflicts were resolved either because the scaffolds involved had not been selected for physical mapping or because the orientation determined by physical mapping was not confident and was thus inverted.

**Table S7. Physical mapping and synteny-based adjacency comparisons**

Comparisons of physical mapping and synteny-based adjacencies for six of the anophelines.

| Species | Synteny Set | Physical mapping with conflicts | Physical mapping with no conflicts | Common to physical mapping & synteny | Synteny with no conflicts | Synteny with conflicts |
|---|---|---|---|---|---|---|
| *Anopheles albimanus* | 3-way | 0 | 31 | 0 | 0 | 0 |
| | 2-way | 0 | 30 | 1 | 0 | 0 |
| | liberal | 2 | 26 | 3 | 1 | 2 |
| | ADSEQ | 0 | 31 | 0 | 1 | 0 |
| | GOS-ASM | 1 | 29 | 1 | 0 | 1 |
| | ORTHOSTITCH | 1 | 27 | 3 | 0 | 1 |
| *Anopheles atroparvus* | 3-way | 0 | 30 | 1 | 0 | 0 |
| | 2-way | 1 | 25 | 5 | 9 | 1 |
| | liberal | 4 | 18 | 9 | 33 | 4 |
| | ADSEQ | 3 | 21 | 7 | 31 | 4 |
| | GOS-ASM | 2 | 26 | 3 | 1 | 2 |
| | ORTHOSTITCH | 3 | 23 | 5 | 17 | 3 |
| *Anopheles funestus* | 3-way | 3 | 74 | 8 | 37 | 2 |
| | 2-way | 14 | 40 | 31 | 174 | 13 |
| | liberal | 19 | 21 | 45 | 272 | 23 |
| | ADSEQ | 20 | 18 | 47 | 258 | 26 |
| | GOS-ASM | 11 | 62 | 12 | 80 | 8 |
| | ORTHOSTITCH | 16 | 40 | 29 | 165 | 14 |
| *Anopheles sinensis (Chinese)* | 3-way | 0 | 20 | 0 | 27 | 0 |
| | 2-way | 2 | 18 | 0 | 90 | 2 |
| | liberal | 5 | 12 | 3 | 225 | 6 |
| | ADSEQ | 5 | 15 | 0 | 152 | 6 |
| | GOS-ASM | 5 | 13 | 2 | 159 | 5 |
| | ORTHOSTITCH | 1 | 18 | 1 | 75 | 1 |
| *Anopheles stephensi (SDA-500)* | 3-way | 3 | 0 | 0 | 51 | 2 |
| | 2-way | 3 | 0 | 0 | 174 | 3 |
| | liberal | 3 | 0 | 0 | 278 | 3 |
| | ADSEQ | 3 | 0 | 0 | 274 | 3 |
| | GOS-ASM | 3 | 0 | 0 | 104 | 2 |
| | ORTHOSTITCH | 3 | 0 | 0 | 174 | 3 |
| *Anopheles stephensi (Indian)* | 3-way | 0 | 4 | 2 | 38 | 0 |
| | 2-way | 1 | 2 | 3 | 124 | 1 |
| | liberal | 1 | 2 | 3 | 184 | 1 |
| | ADSEQ | 2 | 1 | 3 | 195 | 3 |
| | GOS-ASM | 1 | 3 | 2 | 66 | 1 |
| | ORTHOSTITCH | 1 | 2 | 3 | 140 | 1 |

### [9] RNA sequencing data from 13 anophelines

*Robert M. Waterhouse, Matthew W. Hahn, Simo V. Zhang*

Transcriptome data from RNA sequencing (RNAseq) experiments can provide additional information about putative scaffold adjacencies when individual transcripts (or paired-end reads) reliably map to scaffold extremities. The Annotated Genome Optimization Using Transcriptome Information (AGOUTI) tool (Zhang et al. 2016) employs RNAseq data to identify such adjacencies as well as correcting any fragmented gene models at the ends of scaffolds. AGOUTI v0.3.3-24-g64c2a76 was applied to 13 anopheline assemblies using genome-mapped paired-end RNAseq data available from VECTORBASE (Giraldo-Calderón et al. 2015) (Release VB-2017-02), including those from the *Anopheles* 16 Genomes Project (Neafsey et al. 2015) and an *A. stephensi* (Indian) male/female study (Jiang et al. 2015). These data were downloaded from VECTORBASE in the form of pre-computed BAM files – RNAseq reads aligned to the assemblies using HISAT2 version 2.0.4 (Kim et al. 2015). All BAM files were sorted by read name (required by AGOUTI), and where more than one BAM file was available for a given assembly they were first merged, both sorting and merging was performed using SAMTOOLS version 0.1.19-44428cd (Li et al. 2009). AGOUTI was run in scaffold mode with default parameters, e.g. for *A. dirus* 'python2 agouti.py scaffold -assembly anopheles-dirus.fa -bam AdirW1.sorted.bam -gff anopheles-dirus.gff3 -outdir ADIRU'. The numbers of resulting predicted adjacencies ranged from just two for *A. albimanus* to more than 200 *A. sinensis* (SINENSIS) (**Table S8**).

**Table S8. AGOUTI-based scaffold adjacencies from 13 anophelines**
Assemblies with paired-end RNAseq BAM files from VECTORBASE used to run AGOUTI to predict scaffold adjacencies from transcriptome data.

| Species | Assembly | Gene set | RNAseq Dataset(s) | Adjacencies |
|---|---|---|---|---|
| *Anopheles albimanus* | AalbS1 | AalbS1.4 | SRS259216_Generic_RNAseq_for_gene_prediction_AalbS1 | 2 |
| *Anopheles arabiensis* | AaraD1 | AaraD1.5 | SRS259215_Generic_RNAseq_for_gene_prediction_AaraD1 | 34 |
| *Anopheles atroparvus* | AatrE1 | AatrE1.4 | SRP021065_Generic_RNAseq_for_gene_prediction_AatrE1 | 39 |
| *Anopheles dirus* | AdirW1 | AdirW1.4 | SRP021066_Generic_RNAseq_for_gene_prediction_AdirW1 | 21 |
| *Anopheles epiroticus* | AepiE1 | AepiE1.4 | SRP043018_Generic_RNAseq_for_gene_prediction_AepiE1 | 27 |
| *Anopheles farauti* | AfarF1 | AfarF1.2 | SRP020562_merged_AfarF1 | 48 |
| *Anopheles funestus* | AfunF1 | AfunF1.5 | SRP021067_Generic_RNAseq_for_gene_prediction_AfunF1 | 94 |
| *Anopheles merus* | AmerM1 | AmerM1.2 | SRP020545_merged_AmerM1 | 159 |
| *Anopheles minimus* | AminM1 | AminM1.4 | SRP021068_Generic_RNAseq_for_gene_prediction_AminM1 | 16 |
| *Anopheles quadriannulatus* | AquaS1 | AquaS1.5 | SRS259214_Generic_RNAseq_for_gene_prediction_AquaS1 | 96 |
| *Anopheles sinensis* | AsinS2 | AsinS2.2 | SRP035663_Generic_RNAseq_for_gene_prediction_AsinS2 | 210 |
| *Anopheles stephensi* | AsteS1 | AsteS1.4 | SRP020546_Generic_RNAseq_for_gene_prediction_(AGC)_AsteS1<br>SRP052094-SRP052164_MSQ43_cell_line_AsteS1 | 99 |
| *Anopheles stephensi (Indian)* | AsteI2 | AsteI2.3 | SRS866621-SRS866625_Male_AsteI2<br>SRS866626-SRS866630_Female_AsteI2 | 198 |

As for the comparisons of the physical mapping results with the synteny-based results, CAMSA was used to compare the two-way consensus sets, as well as the conservative three-way consensus sets and the liberal union sets of all non-conflicting adjacencies, with the AGOUTI-based adjacencies from each of the 13 assemblies and quantify agreements as well as conflicting and unique adjacencies (**Table S9**). The AGOUTI-based scaffold adjacencies supported up to 17-20% of two-way consensus synteny-based adjacencies in some species, with generally few conflicts but up to 11% and 14% conflicting for *A. stephensi* (Indian) and *A. sinensis* (SINENSIS), respectively, which had the most AGOUTI-based scaffold adjacencies. Across all 13 assemblies, 18% of AGOUTI-based scaffold adjacencies supported the two-way consensus synteny-based adjacencies, with only 7% in conflict and 75% were unique to the AGOUTI sets.

**Table S9. AGOUTI and synteny-based adjacency comparisons**
Comparisons of AGOUTI and synteny-based adjacencies for 13 of the anophelines.

| Species | Synteny Set | AGOUTI with conflicts | AGOUTI with no conflicts | Common to AGOUTI & synteny | Synteny with no conflicts | Synteny with conflicts |
|---|---|---|---|---|---|---|
| *Anopheles albimanus* | 3-way | 0 | 2 | 0 | 1 | 0 |
| | 2-way | 0 | 2 | 0 | 6 | 0 |
| | liberal | 0 | 2 | 0 | 1 | 0 |
| | ADSEQ | 0 | 2 | 0 | 2 | 0 |
| | GOS-ASM | 0 | 2 | 0 | 4 | 0 |
| | ORTHOSTITCH | 0 | 34 | 0 | 2 | 0 |
| *Anopheles arabiensis* | 3-way | 2 | 32 | 0 | 19 | 2 |
| | 2-way | 5 | 27 | 2 | 50 | 7 |
| | liberal | 5 | 28 | 1 | 40 | 7 |
| | ADSEQ | 1 | 32 | 1 | 9 | 1 |
| | GOS-ASM | 3 | 31 | 0 | 26 | 3 |
| | ORTHOSTITCH | 0 | 39 | 0 | 1 | 0 |
| *Anopheles atroparvus* | 3-way | 1 | 37 | 1 | 13 | 1 |
| | 2-way | 5 | 32 | 2 | 39 | 5 |
| | liberal | 7 | 30 | 2 | 33 | 7 |
| | ADSEQ | 0 | 39 | 0 | 6 | 0 |
| | GOS-ASM | 4 | 34 | 1 | 20 | 4 |
| | ORTHOSTITCH | 0 | 20 | 1 | 8 | 0 |
| *Anopheles dirus* | 3-way | 1 | 18 | 2 | 33 | 1 |
| | 2-way | 3 | 16 | 2 | 60 | 3 |
| | liberal | 1 | 18 | 2 | 60 | 1 |
| | ADSEQ | 1 | 18 | 2 | 16 | 1 |
| | GOS-ASM | 2 | 18 | 1 | 36 | 2 |
| | ORTHOSTITCH | 0 | 25 | 2 | 72 | 0 |
| *Anopheles epiroticus* | 3-way | 2 | 16 | 9 | 391 | 2 |
| | 2-way | 5 | 11 | 11 | 565 | 5 |
| | liberal | 5 | 12 | 10 | 593 | 5 |
| | ADSEQ | 1 | 22 | 4 | 138 | 1 |
| | GOS-ASM | 4 | 14 | 9 | 443 | 4 |
| | ORTHOSTITCH | 0 | 45 | 3 | 40 | 0 |
| *Anopheles farauti* | 3-way | 4 | 33 | 11 | 90 | 4 |
| | 2-way | 6 | 16 | 26 | 167 | 7 |
| | liberal | 5 | 22 | 21 | 150 | 6 |
| | ADSEQ | 1 | 36 | 11 | 63 | 1 |
| | GOS-ASM | 6 | 32 | 10 | 100 | 6 |
| | ORTHOSTITCH | 1 | 82 | 11 | 35 | 1 |
| *Anopheles funestus* | 3-way | 5 | 49 | 40 | 172 | 6 |
| | 2-way | 14 | 29 | 51 | 272 | 17 |
| | liberal | 14 | 27 | 53 | 261 | 17 |
| | ADSEQ | 1 | 72 | 21 | 77 | 2 |
| | GOS-ASM | 11 | 50 | 33 | 164 | 11 |
| | ORTHOSTITCH | 3 | 142 | 14 | 101 | 3 |

| | | | | | | |
|---|---|---|---|---|---|---|
| *Anopheles merus* | 3-way | 13 | 103 | 43 | 318 | 11 |
| | 2-way | 23 | 68 | 68 | 536 | 20 |
| | liberal | 20 | 76 | 63 | 509 | 18 |
| | ADSEQ | 5 | 131 | 23 | 192 | 5 |
| | GOS-ASM | 19 | 99 | 41 | 356 | 16 |
| | ORTHOSTITCH | 0 | 14 | 2 | 1 | 0 |
| *Anopheles minimus* | 3-way | 0 | 14 | 2 | 10 | 0 |
| | 2-way | 2 | 12 | 2 | 19 | 2 |
| | liberal | 2 | 12 | 2 | 15 | 2 |
| | ADSEQ | 0 | 14 | 2 | 5 | 0 |
| | GOS-ASM | 0 | 14 | 2 | 12 | 0 |
| | ORTHOSTITCH | 4 | 86 | 6 | 39 | 4 |
| *Anopheles quadriannulatus* | 3-way | 7 | 69 | 20 | 102 | 7 |
| | 2-way | 14 | 54 | 28 | 192 | 15 |
| | liberal | 13 | 56 | 27 | 197 | 14 |
| | ADSEQ | 10 | 69 | 17 | 95 | 10 |
| | GOS-ASM | 15 | 65 | 16 | 118 | 14 |
| | ORTHOSTITCH | 7 | 199 | 4 | 20 | 6 |
| *Anopheles sinensis (SINENSIS)* | 3-way | 18 | 178 | 14 | 87 | 17 |
| | 2-way | 38 | 147 | 25 | 271 | 41 |
| | liberal | 37 | 146 | 27 | 271 | 37 |
| | ADSEQ | 33 | 161 | 16 | 149 | 31 |
| | GOS-ASM | 25 | 172 | 13 | 167 | 24 |
| | ORTHOSTITCH | 9 | 188 | 1 | 30 | 9 |
| *Anopheles stephensi (Indian)* | 3-way | 14 | 176 | 8 | 106 | 14 |
| | 2-way | 23 | 164 | 11 | 153 | 24 |
| | liberal | 23 | 162 | 13 | 164 | 24 |
| | ADSEQ | 11 | 184 | 3 | 55 | 11 |
| | GOS-ASM | 17 | 174 | 7 | 120 | 17 |
| | ORTHOSTITCH | 2 | 86 | 11 | 40 | 2 |
| *Anopheles stephensi (SDA-500)* | 3-way | 6 | 58 | 35 | 137 | 5 |
| | 2-way | 12 | 32 | 55 | 216 | 10 |
| | liberal | 9 | 36 | 54 | 214 | 9 |
| | ADSEQ | 8 | 77 | 14 | 86 | 6 |
| | GOS-ASM | 9 | 52 | 38 | 132 | 7 |
| | ORTHOSTITCH | 0 | 2 | 0 | 1 | 0 |

### [10] Building the PacBio-based *Anopheles funestus* assembly

*Paul I. Howell, Sergey Koren, Adam M. Phillippy, Nora J. Besansky, Scott J. Emrich*

A new *A. funestus* assembly, AfunF2-IP, was generated using approximately 70X of PacBio sequencing data and polished with QUIVER (PacBio's SMRT Analysis software suite). This was merged with the reference assembly (AfunF1) using METASSEMBLER (Wences and Schatz 2015) to generate a merged assembly. Finally, the merged assembly was scaffolded with SSPACE (Boetzer et al. 2011) using the available Illumina sequencing data. Summary statistics for the reference AfunF1, PacBio only, Illumin+PacBio Merged, and Merged+Scaffolded AfunF2-IP assemblies (using 225Mbp as a genome size) are presented in **Table S10** and **Figure S10** (to compute contig statistics, the scaffolds were split at three consecutive Ns).

At the contig level the new AfunF2-IP assembly is an improvement over the reference AfunF1, e.g. the number of contigs is reduced from 9'880 to 4'170 and the NG50 increases from 47 Kbp to 194 Kbp. However, longer-range scaffolding of these contigs unfortunately failed to produce a better quality scaffold-level assembly. In terms of gene content, analysis with 2'799 dipteran Benchmarking Universal Single-Copy Orthologues (BUSCOs) (Simão et al. 2015; Waterhouse et al. 2018, 2019)indicates that despite the better contigs fewer BUSCOs are found as complete genes in the AfunF2-IP assembly (**Table S10**).

The AfunF1 assembly has a very high level of N's, 15.63% compared with just 0.90% for the AfunF2-IP assembly, reflecting how scaffolding improves N50 measures but mainly by joining contigs with stretches of unknown nucleotides (N's). When the scaffolds are artificially de-scaffolded by splitting them at consecutive runs of 3, 300, and 1'000 Ns the new AfunF2-IP assembly is clearly much better (**Figure S10**). The stringent splitting at N>=3 also indicates the greater integrity of the sequence quality of the AfunF2-IP assembly as this does not result in high fragmentation levels as it does for AfunF1 (i.e. from 3'772 scaffolds to 4'186 contigs for AfunF2-IP but from 1'391 scaffolds to 9'878 contigs for AfunF1).

### Table S10. Comparisons of *Anopheles funestus* assemblies

Statistics describing the *Anopheles funestus* old reference AfunF1, PacBio only, Illumina+PacBip Merged, and Merged+Scaffolded AfunF2-IP assemblies.

| | | scaffolds | contigs | BUSCO Scores (% of 2'799 dipteran BUSCOs) Complete[Single-Copy,Duplicated],Fragmented,Missing |
|---|---|---|---|---|
| **Reference AfunF1** | Count | 1,392 | 9,880 | C:98.4%[S:97.9%,D:0.5%],F:1.0%,M:0.6%, |
| | Total Basepairs | 225,223,604 | 190,015,44 | |
| | NG50 | 671,960 | 47,164 | |
| | Maximum | 3,832,769 | 563,645 | |
| **PacBio only** | Count | N/A | 18,595 | N/A |
| | Total Basepairs | N/A | 445,128,909 | |
| | NG50 | N/A | 147,143 | |
| | Maximum | N/A | 4,813,330 | |
| **Illumina+PacBio Merged** | Count | N/A | 4,653 | N/A |
| | Total Basepairs | N/A | 260,811,249 | |
| | NG50 | N/A | 146,657 | |
| | Maximum | N/A | 3,313,857 | |
| **Merged+Scaffolded AfunF2-IP** | Count | 3,773 | 4,170 | C:92.5%[S:85.8%,D:6.7%],F:4.7%,M:2.8% |
| | Total Basepairs | 263,192,532 | 260,811,631 | |
| | NG50 | 244,910 | 194,030 | |
| | Maximum | 7,451,746 | 3,313,857 | |

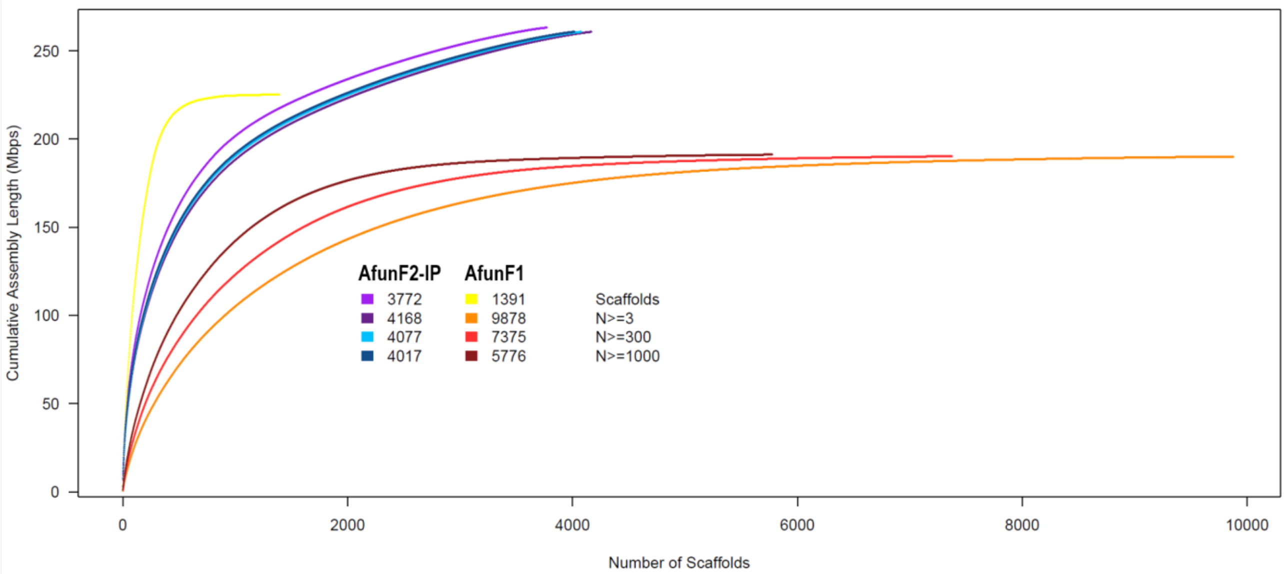

**Figure S10. Cumulative scaffold lengths for *Anopheles funestus* AfunF1 and AfunF2-IP assemblies**

Cumulative assembly length plots for the reference AfunF1 and the new AfunF2-IP *Anopheles funestus* scaffold-level assemblies. Lengths are summed and plotted from the longest to the shortest scaffold for each assembly. These are replotted for each assembly after splitting scaffolds at consecutive runs of 3, 300, and 1'000 Ns, i.e. effectively de-scaffolding them and slicing at ambiguous or low-quality regions.

### [11] Examining collinearity between the two *Anopheles funestus* assemblies

*Robert M. Waterhouse*

Despite the lack of longer-range scaffolding information from the AfunF2-IP assembly, the scaffolds are nonetheless useful for the purposes of identifying potential adjacencies of the AfunF1 scaffolds through whole genome alignment analyses. The first step towards delineating the order and orientation of *A. funestus* AfunF1 scaffolds along those of the AfunF2-IP assembly was to mask each assembly with a library of anopheline repeats using REPEATMASKER (Smit et al. 2015) and then perform a pairwise LASTZ (Harris 2007) whole genome alignment with default parameters. The resulting alignment blocks were then interrogated with a custom Perl script to define alignment blocks of more than 10 basepairs (bps) from AfunF1 allowing for insertions or deletions of no more than 10 bps in either assembly and requiring AfunF1 genomic regions to be unique (basepairs falling in regions that appeared in more than one alignment block were ignored unless the second-best scoring block scored less than 75% of the best-scoring block, in which case only the best-scoring block was considered). This identified a total of 124’926 links connecting 1’098 AfunF1 scaffolds to 2’845 AfunF2-IP scaffolds with a mean length of 1’234 bps, median of 650 bps, and maximum of 31’044 bps.

Links were then bundled into larger link-regions allowing a maximum of 30 Kbps between links from the same pairs of scaffolds with the same orientations. The largest bundle (by genomic span of the bundled links) for each AfunF1 scaffold was used to define the corresponding AfunF2-IP scaffold and its mapping location was set at the midpoint of the bundle’s genomic span on the AfunF2-IP scaffold, thereby ordering and orientating *A. funestus* AfunF1 scaffolds along their corresponding AfunF2-IP scaffolds and producing a final set of 321 alignment-based scaffold adjacencies. Each set of predicted adjacencies, the consensus adjacencies, the physical mapping adjacencies, and the AGOUTI adjacencies were compared with the set of alignment-based scaffold adjacencies (**Table S11**). As the alignments consider scaffolds regardless of whether they were targeted for physical mapping or if they have any annotated orthologues, short un-annotated scaffolds may be ordered and oriented that then result in conflicts with the synteny-based or physical mapping based adjacencies that do not consider such scaffolds. Ignoring short scaffolds (<5 Kbps) or scaffolds with less than 30% aligned sequence reduces the total number of alignment-based scaffold adjacencies by about half to just 154, but this results in additional supported adjacencies being recovered for all the comparison sets (**Table S11**). The ordered and oriented scaffolds were visualised using CIRCOS (Krzywinski et al. 2009) to display alignments greater than 100 bps, and bundled links greater than 3 Kbps and examine the concordance between the different adjacency predictions (**Figure 5, main text; Figure S11**).

**Table S11. Alignment-based adjacency comparisons for *Anopheles funestus***

Comparisons of adjacencies based on alignments of *Anopheles funestus* AfunF1 and AfunF2-IP assemblies with synteny-based, AGOUTI-based, and physical mapping based adjacencies.

| Adjacency Set | Adjacencies | Alignment-based with conflicts | Alignment-based with no conflicts | Common to alignment-based & other | Other with conflicts | Other with no conflicts | Additional Supported adjacencies |
|---|---|---|---|---|---|---|---|
| ADSEQ | 331 | 101 | 162 | 58 | 197 | 76 | 18 |
| GOS-ASM | 100 | 26 | 281 | 14 | 66 | 20 | 5 |
| ORTHOSTITCH | 208 | 66 | 223 | 32 | 130 | 46 | 14 |
| LIBERAL UNION | 340 | 102 | 164 | 55 | 208 | 77 | 18 |
| 2-WAY CONSENSUS | 218 | 61 | 223 | 37 | 136 | 45 | 14 |
| 3-WAY CONSENSUS | 47 | 13 | 303 | 5 | 32 | 10 | 4 |
| PHYSICAL MAPPING | 85 | 65 | 237 | 19 | 24 | 42 | 14 |
| AGOUTI | 94 | 29 | 278 | 14 | 56 | 24 | 2 |

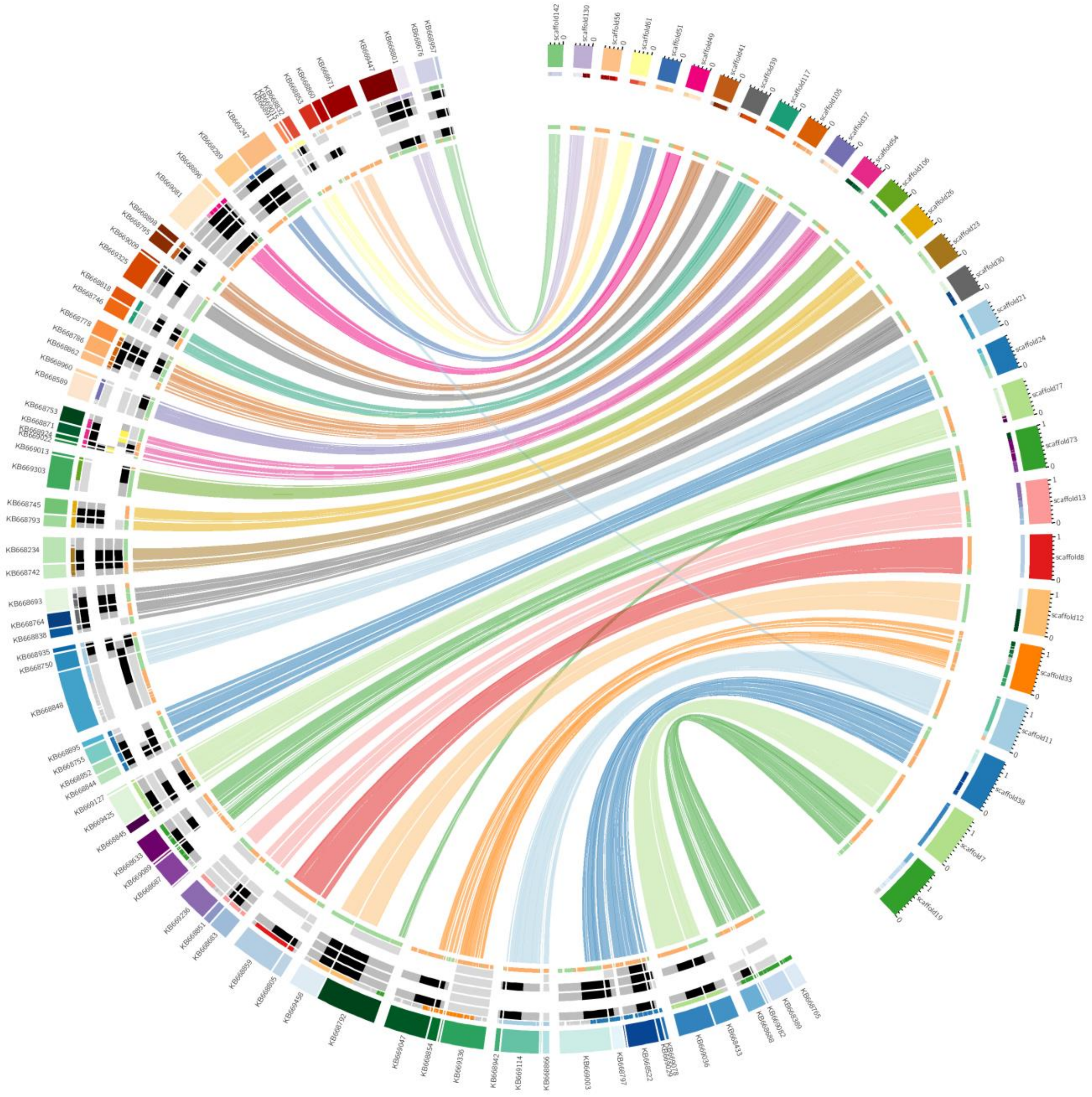

**Figure S11. Collinearity between *Anopheles funestus* AfunF1 and AfunF2-IP scaffolds**

*Anopheles funestus* scaffold adjacencies supported by collinearity with the new AfunF2-IP assembly. The plot shows correspondences of AfunF1 scaffolds with AfunF2-IP scaffolds based on whole genome alignment data, with links coloured according to their AfunF2-IP scaffold. Synteny-based adjacency predictions between AfunF1 scaffolds are highlighted with a track showing confirmed neighbours (black), supported neighbours with conflicting orientations (yellow), scaffolds with predicted adjacencies that are not supported by the alignments (light grey) for: from outer to inner tracks, ADSEQ, GOS-ASM, ORTHOSTITCH, physical mapping, and AGOUTI. The innermost track shows alignments in forward (green) and reverse (orange) orientations. The outermost track shows alignments coloured according to the corresponding scaffold in the other assembly (if they align to scaffold not shown on the plot they appear light grey). AfunF1 scaffolds are labelled KB66XXXX and the AfunF2-IP scaffolds are labelled scaffoldX.

### [12] Reconciliation to build the new assemblies

*Robert M. Waterhouse, Jiyoung Lee, Livio Ruzzante, Maarten J.M.F. Reijnders, Romain Feron, Daniel Lawson, Gareth Maslen, Igor V. Sharakhov*

In order to build the new assemblies for *A. albimanus*, *A. atroparvus*, *A. farauti*, *A. melas*, and *A. merus*, results from the two-way consensus synteny predictions, and AGOUTI and physical mapping data (where available), had to be compared and reconciled with their version 2 reference assemblies. For the published *A. albimanus* AalbS2 assembly, new physical mapping data (also used in this study) was used to improve the assembly by correcting nine misassemblies and anchoring 98% to chromosomes (Artemov et al. 2017). This splitting of the misassembled scaffolds resulted in an increase from 204 AalbS1 scaffolds to 236 AalbS2 scaffolds. The single synteny-based prediction from the two-way consensus set was in agreement with the physical mapping data, as were two of the three adjacencies unique to ORTHOSTITCH, and were therefore already present in the upgraded AalbS2 chromosomal assembly. AGOUTI predicted only two adjacencies, both of which were between very short scaffolds (1'148 bp and 1'012 bp) with no gene annotations and much longer already anchored scaffolds (**Table 2, main text**).

For the published *A. atroparvus* AatrE2 assembly, and later AatrE3, additional physical mapping data (also used in this study) was used to anchor 56 scaffolds (201 Mbps, 89.6% of the assembly) to chromosomes, leaving 1'315 scaffolds unmapped (Artemov et al. 2018a). The *A. melas* AmelC2 assembly was produced from the AmelC1 assembly following the removal of several duplicated scaffolds and regions of scaffolds thereby reducing the number of scaffolds by 52 to 20'229 scaffolds with an unchanged scaffold N50 of 18 Kbps. This affected only 112 scaffolds that were part of 121 adjacencies, and where removed regions made up less than 25% of the original scaffold and they were removed from scaffold ends not involved in any adjacencies then these adjacencies were retained. Thus 95% of AmelC1 adjacencies (97% of scaffolds) were reconciled with the AmelC2 assembly and were used to build the AmelC3 assembly.

The version 2 assemblies for *A. farauti* (AfarF2) and *A. merus* (AmerM2) were derived from re-scaffolding efforts that included the addition of a large-insert 'fosill' sequencing library constructed from high molecular weight DNA, which reduced the numbers of scaffolds from 550 to 310 and 2'753 to 2'027 and increased N50 values from 1'197 Kbps to 12'895 Kbps and 342 Kbps to 1'490 Kbps, respectively. The version 1 assemblies were aligned to the version 2 assemblies using BLAST+ (Camacho et al. 2009) and all scaffolds involved in the synteny-based or AGOUTI-based adjacency predictions were visualised with their corresponding version 2 scaffolds using CIRCOS (Krzywinski et al. 2009). In this way, the predicted adjacencies from version 1 assemblies were assessed to identify adjacencies fully supported by alignments to version 2 scaffolds, e.g. seven *A. farauti* synteny-based two-way consensus set adjacencies confirmed by the alignment with a single AfarF2 scaffold (**Figure S12**). These assessments also identified adjacencies without support from the version 2 assemblies but which were nonetheless not in conflict (i.e. predicted neighbouring scaffolds that were not joined during the re-scaffolding process), supported neighbours but conflicting orientations, and adjacencies where the arrangements in corresponding version 2 scaffolds precluded the possibility of being neighbours (**Table S12**).

**Table S12. Version 2 assembly reconciliations for *Anopheles farauti* and *Anopheles merus***
Reconciliation of adjacencies for *A. farauti* and *A. merus* with their version 2 assemblies.

| Species | Prediction Set | Number of Adjacencies | Fully Supported | Non-Conflicting | Conflicting |
|---|---|---|---|---|---|
| *Anopheles farauti* | Agouti | 48 | 39 (81.2%) | 2 (4.2%) | 7 (14.6%) |
| | Two-way synteny | 105 | 91 (86.7%) | 9 (8.6%) | 5 (4.7%) |
| *Anopheles merus* | Agouti | 159 | 106 (66.7%) | 20 (12.6%) | 33 (20.7%) |
| | Two-way synteny | 372 | 305 (82.0%) | 31 (8.3%) | 36 (9.7%) |

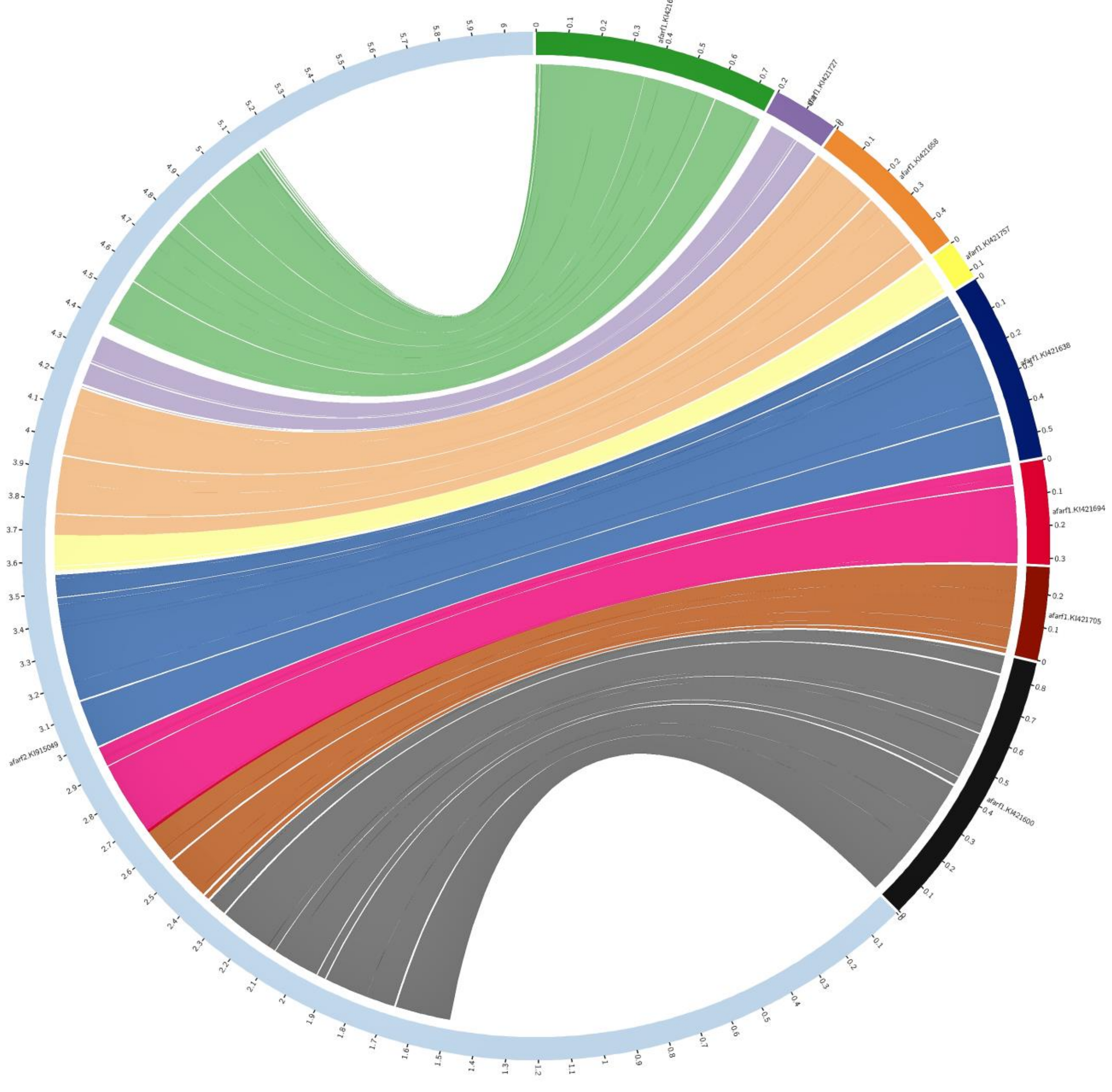

**Figure S12. Collinearity between *Anopheles farauti* AfarF1 and AfarF2 scaffolds**

*Anopheles farauti* AfarF1 scaffold adjacencies supported by collinearity with the subsequent AfarF2 assembly. Seven adjacencies from the *A. farauti* synteny-based two-way consensus set predicted the order and orientation of eight AfarF1 scaffolds that are fully supported by the alignment with a single AfarF2 scaffold. Scaffold lengths are shown in increments of 0.1 Mbps. AfarF2 KI915049 aligned with AfarF1 KI421600, KI421705, KI421694, KI421638, KI421757, KI421658, KI421727, KI421610.

### New assembly FASTA files and annotation 'lift-over' details

The final lists of pairwise adjacencies and the superscaffolds, with superscaffolds presented in a GRIMM-like format (http://grimm.ucsd.edu/GRIMM/grimm_instr.html) were combined with the VECTORBASE (Release VB-2019-02) assembly sequence data (FASTA format) and assembly annotation data (GFF3 and GTF formats) to produce the new updated assemblies and their corresponding annotations. The adjacencies defined the neighbouring scaffolds that were fused together with an insertion of a stretch of 100 N's to indicate a sequence gap, and with reversed scaffold orientations as required by the relative orientations of the pairwise adjacencies and superscaffolds. Coordinate systems for annotated features were updated to reflect the fusions and insertions to create the superscaffolds with all mapped features. Annotation versions used for lift-overs were: AalbS2.6, AaraD1.10, AatrE3.1, AchrA1.7, AcolM1.8, AculA1.6, AdarC3.8, AdirW1.8, AepiE1.7, AfarF2.5, AfunF1.10, AmacM1.5, AmelC2.6, AmerM2.9, AminM1.8, AquaS1.11, AsinS2.5, AsinC2.2, AsteS1.7, AsteI2.3. For the eight assemblies with chromosome-mapped scaffolds and superscaffolds, AGP (A Golden Path) formatted files were built or updated to assign all finalised scaffolds to chromosomal locations. The authors acknowledge the help provided by Vasily Sitnik at VECTORBASE with the process of updating and submitting the new assemblies and annotations and AGP files.

### Chromosome arm assignment using updated assemblies and annotations

Several whole-arm translocations in the anophelines (Neafsey et al. 2015) mean that the five chromosomal elements that make up the X chromosome and the two autosomes correspond to different named chromosome arms in different species (**Table S13**), and thus results are presented as assignments to elements one to five rather than named chromosome arms. Combining orthology data delineated for genes from all 21 assemblies (see section [3] above) and chromosome arm locations for genes from the eight assemblies with chromosomal anchoring data, orthologues of genes on each scaffold were enumerated for each element from each of the eight chromosome-anchored assemblies. To be considered for assignment, the scaffold was required to have a minimum of ten genes with annotated orthologues. The scaffold was then assigned to an element when at least 75% of these orthologues were located on a single element. Confident assignments reported in **Table S1** and main text **Fig. 1** were required to be confirmed by data from at least two species, and conflicting assignments were excluded as they could represent translocation events (assignments with only single-species support or with conflicting species support are reported but flagged as not assigned).

**Table S13. Chromosome arm to element correspondences in anophelines**

For each of the eight assemblies with chromosome anchoring data, the table presents correspondences between chromosomal elements one to five and the named chromosome arms.

| Species | Element 1 | Element 2 | Element 3 | Element 4 | Element 5 |
|---|---|---|---|---|---|
| *A. gambiae* | X | 2R | 2L | 3R | 3L |
| *A. arabiensis* | X | 2R | 2L | 3R | 3L |
| *A. funestus* | X | 2R | 3R | 2L | 3L |
| *A. stephensi* | X | 2R | 3L | 3R | 2L |
| *A. stephensi (Indian)* | X | 2R | 3L | 3R | 2L |
| *A. sinensis (Chinese)* | X | 3R | 2L | 2R | 3L |
| *A. atroparvus* | X | 3R | 2L | 2R | 3L |
| *A. albimanus* | X | 2R | 3L | 2L | 3R |

### [13] Software and database availability

ADSEQ: https://github.com/YoannAnselmetti/ADseq-Anopheles-APBC2018, and https://github.com/YoannAnselmetti/DeCoSTAR_pipeline (Anselmetti et al. 2018)

AGOUTI: https://github.com/svm-zhang/AGOUTI (Zhang et al. 2016)

BESST: https://github.com/ksahlin/BESST, (Sahlin et al. 2014)

BLAST+: ftp://ftp.ncbi.nlm.nih.gov/blast/executables/blast+, (Camacho et al. 2009)

BUSCO: https://busco.ezlab.org, (Waterhouse et al. 2018)

CAMSA: https://github.com/compbiol/CAMSA, (Aganezov and Alekseyev 2017)

CIRCOLETTO: https://github.com/infspiredBAT/Circoletto, (Darzentas 2010)

CIRCOS: http://circos.ca, (Krzywinski et al. 2009)

EULERAPE: http://www.eulerdiagrams.org/eulerAPE, (Micallef and Rodgers 2014)

GOS-ASM: https://github.com/aganezov/gos-asm, (Aganezov and Alekseyev 2016)

HISAT: http://www.ccb.jhu.edu/software/hisat/index.shtml, (Kim et al. 2015)

LASTZ: http://www.bx.psu.edu/miller_lab/dist/README.lastz-1.02.00/README.lastz-1.02.00a.html, (Harris 2007)

METASSEMBLER: https://sourceforge.net/projects/metassembler, (Wences and Schatz 2015)

MUSCLE: https://www.drive5.com/muscle, (Edgar 2004)

ORTHODB: https://www.orthodb.org, (Zdobnov et al. 2017)

ORTHOSTITCH: https://gitlab.com/rmwaterhouse/OrthoStitch, (this study)

PRIMERBLAST: https://www.ncbi.nlm.nih.gov/tools/primer-blast, (Ye et al. 2012)

QUIVER: https://github.com/PacificBiosciences/GenomicConsensus, (PacBio's SMRT Analysis software suite)

RAxML: https://sco.h-its.org/exelixis/web/software/raxml/index.html, (Stamatakis 2014)

REPEATMASKER: http://www.repeatmasker.org, (Smit et al. 2015)

SAMTOOLS: https://github.com/samtools, (Li et al. 2009)

SSPACE: https://www.baseclear.com/services/bioinformatics/basetools/sspace-standard, and https://github.com/nsoranzo/sspace_basic, (Boetzer et al. 2011)

TREERECS: https://gitlab.inria.fr/Phylophile/Treerecs, and https://project.inria.fr/treerecs

VECTORBASE: https://www.vectorbase.org, (Giraldo-Calderón et al. 2015)

**DOCKER container**

A DOCKER container is provided that packages ADSEQ, GOS-ASM, ORTHOSTITCH, and CAMSA, as well as their dependencies, in a virtual environment that can run on a Linux server, this is available from: https://hub.docker.com/r/mreijnders/synteny/

### [14] Main text figure credits

**Figure 1.** Genomic spans of scaffolds and superscaffolds with and without chromosome anchoring or arm assignments for 20 *Anopheles* assemblies. *Robert M. Waterhouse, Livio Ruzzante, Romain Feron*

**Figure 2.** Improved genome assemblies for 20 anophelines from synteny-based scaffold adjacency predictions. *Robert M. Waterhouse, Livio Ruzzante, Maarten J.M.F. Reijnders*

**Figure 3.** Comparisons of synteny-based scaffold adjacency predictions from ADSEQ (AD), GOS-ASM (GA), and ORTHOSTITCH (OS). *Robert M. Waterhouse, Livio Ruzzante*

**Figure 4.** Scaffold adjacency validations with physical mapping and RNA sequencing data. *Robert M. Waterhouse, Maarten J.M.F. Reijnders*

**Figure 5.** Whole genome alignment comparisons of selected *Anopheles funestus* AfunF1 and AfunF2-IP scaffolds. *Robert M. Waterhouse*

**Figure 6.** The *Anopheles funestus* photomap of straightened polytene chromosomes with anchored scaffolds from the AfunF1 and AfunF2-IP assemblies. *Jiyoung Lee, Maria V. Sharakhova, Igor V. Sharakhov*

**Figure 7.** The *Anopheles stephensi* photomap of straightened polytene chromosomes with anchored scaffolds from the AsteI2 assembly. *Jiyoung Lee, Maria V. Sharakhova, Igor V. Sharakhov*
